# Deep molecular characterization linked to drug response profiling of pancreatic ductal adenocarcinoma using patient-derived organoids

**DOI:** 10.1101/2021.08.26.457743

**Authors:** Solange Le Blanc, Naveed Ishaque, Julia Jabs, Tobias Bauer, Sebastian Schuth, Qing Hu, Olivia Debnath, Foo Wei Ten, Carl-Stephan Leonhardt, Anna-Katharina König, Matthias Bieg, Christoph Eckert, Matthias M. Gaida, Michael Volkmar, Daniel Hübschmann, Miriam Schenk, Rienk Offringa, Nathalia A. Giese, Matthias Schlesner, Markus W. Büchler, Roland Eils, Christian Conrad, Oliver Strobel

**Affiliations:** European Pancreas Center, Department of General Surgery, Heidelberg University Hospital, Heidelberg, Germany; Division of Molecular Oncology of Gastrointestinal Tumors, German Cancer Research Center (DKFZ), Heidelberg, Germany; NCT partner site Heidelberg, a clinical-translational cancer research partnership between University Hospital Heidelberg and DKFZ, Germany; Berlin Institute of Health at Charité – Universitätsmedizin Berlin, Digital Health Center; Division of Theoretical Bioinformatics, German Cancer Research Center (DKFZ), Heidelberg, Germany; Merck Healthcare KGaA, Global Research, Darmstadt, Germany; Institute of Pathology, Heidelberg University Hospital, Heidelberg, Germany; Institute of Pathology, University Medical Center Mainz, Mainz, Germany; Computational Oncology, Molecular Precision Oncology Program, National Center for Tumor diseases (NCT) Heidelberg and DKFZ, Heidelberg, Germany; Heidelberg Institute for Stem Cell Technology and Experimental Medicine (HI-STEM), Heidelberg, Germany; Bioinformatics and Omics Data Analytics, German Cancer Research Center (DKFZ), Heidelberg, Germany; Biomedical Informatics, Data Mining and Data Analytics, Augsburg University, Augsburg, Germany; Division of Visceral Surgery, Department of General Surgery, Medical University of Vienna, Vienna, Austria

**Keywords:** Pancreatic cancer, patient-derived organoids, molecular subtypes, drug screening, personalized oncology.

## Abstract

Pancreatic ductal adenocarcinoma (PDAC) is characterized by high drug resistance and poor prognosis. Novel therapeutic and stratification strategies are urgently needed. Here, we present an integration of in-depth genomic and transcriptomic characterization with drug screening and clinical outcome based on a catalogue of 51 patient-derived tumor organoids (PDOs) from resected PDAC. Known PDAC molecular subtypes and their prognostic value are conserved in organoids. Integration of transcriptomic and drug response profiles suggest a metabolism-mediated modulations of drug resistance. Copy number alterations on chromosome 13q and wild-type status of *TP53* emerged as potential novel genomic biomarkers for sensitivity to 5-FU and oxaliplatin treatment, respectively. Functional testing of targeted drugs in PDOs revealed its additional value for genome-driven personalized oncology. Co-deletion of *TP53*/*POLR2A* increased vulnerability to RNA polymerase II inhibition, pointing to a promising target for personalized treatment in PDAC.

**Significance:** Patient-derived PDAC organoids hold great promise as surrogate tumor models for personalized oncology. By integrating highly granular molecular, drug sensitivity and clinical data, we demonstrate that PDOs are valid models for molecular characterization and response prediction that also enable identification of novel drug sensitivity biomarkers and resistance mechanisms in PDAC.

## Introduction

Pancreatic ductal adenocarcinoma (PDAC) is an aggressive malignancy expected to become the second cause of cancer-related deaths by 2030 (1, 2). Surgical resection followed by adjuvant chemotherapy is the best therapeutic option for PDAC patients with resectable disease (3). The current standard adjuvant chemotherapy regimens, gemcitabine plus capecitabine (4) and modified FOLFIRINOX (folinic acid, 5-fluorouracil, irinotecan and oxaliplatin) (5), significantly improve outcome compared to surgery alone. However, the 5-year survival rates remain dismal at around 30% (4). Further evidence indicates that chemotherapy sensitivity might be dependent on molecular subtypes of PDAC (6). Molecular classification strategies for PDAC tumors (7–9) converged into two widely recognized transcriptional subtypes with prognostic value, a classical/progenitor subtype and a basal-like/squamous subtype (10). The basal-like subtype is associated with a more aggressive phenotype, worse response to chemotherapy and poorer clinical outcome (6). However, molecular characterization of PDAC tumors based on tumor biopsy samples is hampered by low epithelial tumor cellularity and high stroma content. In addition, neoadjuvant treatment before surgery further challenges the use of resected samples for high throughput sequencing. This hinders the understanding of the molecular landscape of this disease and limits therapy guidance based on molecular data. Hence, representative models, improved stratification strategies and novel therapeutic targets for resected PDAC patients are urgently needed.

Patient-derived organoids (PDOs) have emerged as a 3D *in vitro* culture system with the potential to retain and reflect the genetic complexity of human cancers (11). The organoid culture system allows for the enrichment and long-term expansion of epithelial tumor cells directly from bulk tumor tissue biopsies, overcoming the limitations of low tumor cellularity, and enables functional testing of drug sensitivities (12, 13).

Previous reports characterizing PDAC-PDO libraries point to their value as tumor avatars for identifying new molecular traits (14) and as predictive tools for personalized oncology (12, 15). However, data on the molecular representability of PDOs of the parental tumor and the molecular stability of PDOs during culture are sparse. More importantly, a comprehensive integration of PDO molecular data and drug response profiling as well as validation of their clinical predictive value are lacking.

In the present work, we generated a catalogue of 51 PDAC-PDOs derived from resected tumors of 44 patients and performed extensive molecular characterization by RNA and whole-genome sequencing. We demonstrate that our PDAC organoids remain largely representative of the original tumors at genomic and transcriptomic levels during culture. Additionally, based on integration of genomic, transcriptomic and drug sensitivity data for 39 PDOs, we identified candidate mechanisms of drug resistance and biomarkers of drug response. Furthermore, PDO drug sensitivity data was correlated with clinical outcomes to interrogate the predictive potential of organoid pharmacotyping.

## Results

### Patient-derived PDAC organoid catalogue and its mutational landscape

We successfully established 51 PDAC-PDO cultures from 44 resected patients (**Fig. 1A**) with an efficiency of approximately 60%, using a modification of the protocol described by Boj et al. (16). The clinicopathological characteristics of the PDAC organoid catalogue are described in **Supplementary Table S1A**. In the majority of cases, one organoid line was generated per patient (n = 39). For patients 077 and 080, three and two organoid lines, respectively, were established from adjacent tumor sites (e.g. aPO, bPO, cPO) while for patients 034, 083 and 093, organoid lines were derived from the primary tumor (PO) and synchronous metastases (MO). A total of 11 organoid lines (21.6%) were derived from patients which had undergone neoadjuvant treatment. The majority of the PDOs displayed a mono-luminal morphology while a few lines formed predominantly multi-luminal and solid structures (**Fig. 1B** and **C**, “Morphology PDO”). Strikingly, in 84% of the cases, the predominant morphology of the epithelial architecture of the original tumor was closely recapitulated by the PDO (**Fig. 1C**, “Morphology adjacent tumor” and “Morphology PDO”).

**Figure 1.**
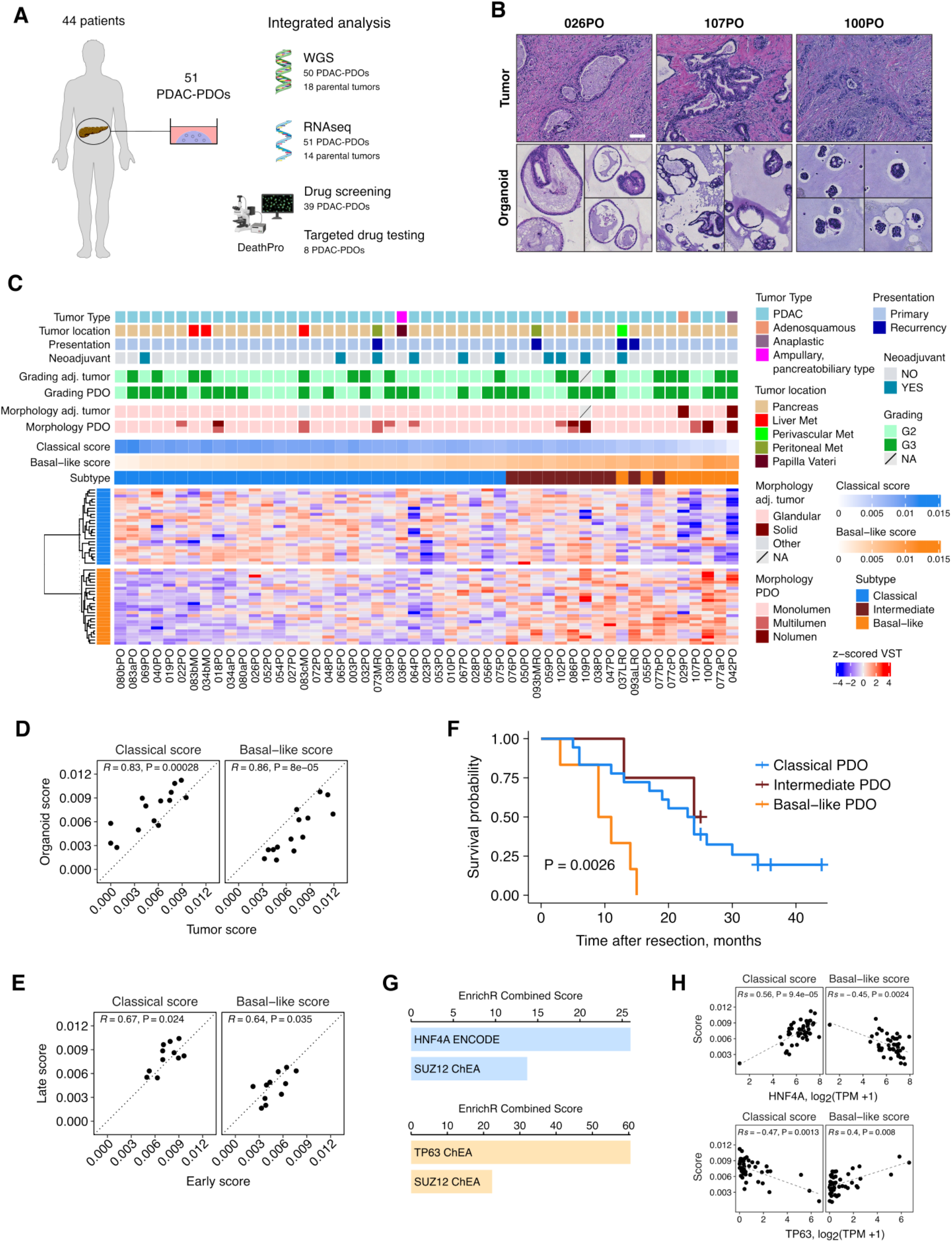
PDAC organoid catalogue: transcriptional subtyping of PDAC-PDOs is prognostic. **A,** Schematic overview of the analyses performed with the established PDAC-PDOs. **B,** H&E staining of PDAC organoids and parental tumor tissues. Organoid growth patterns give rise to mono-luminal, multi-luminal and solid morphologies, showing resemblance to the histological features of the parental tumors. Scale bar 100 µm. **C,** Expression heatmap of Moffitt’s classical and basal-like signature genes. NMF-derived classical and basal-like scores are indicated, as well as the classification of each organoid line into three subtype classes based on the scores. Samples are sorted by the ratio basal-like/classical score. Clinicopathological variables, adjacent tumor grading (Grading adj. tumor), organoid grading (Grading PDO), predominant adjacent tumor morphology (Morphology adj. tumor) and predominant organoid morphology (Morphology PDO) are annotated. The majority of the lines were derived from PDAC, including rare subtypes such as adenosquamous (n = 2), anaplastic (n = 1) and ampullary, pancreatobiliary type (n = 1) carcinomas. **D,** Correlation of classical and basal-like scores between parental tumor tissues and organoids. R: Pearson’s correlation coefficient. **E,** Correlation of classical and basal-like scores for organoid samples at early and later time points in culture. R: Pearson’s correlation coefficient. **F,** Kaplan-Meier curves (log-rank test) for overall survival, stratified by organoid subtypes. Only organoids derived from chemo-naïve patients were included in the survival analysis. **G,** Enriched ENCODE/ChEA consensus transcription factors datasets (adjusted *P* < 0.01) in differentially expressed genes upregulated in classical (top) and basal-like (bottom) organoids. **H,** Expression of *HNF4A* (top) and *TP63* (bottom) according to the classical and basal-like scores. Rs: Spearman’s correlation coefficient. TPM: transcript per million.

Whole-genome sequencing (WGS) and RNAseq were performed for all organoid lines. For analysis of recurrent genomic aberrations in the cohort, one representative sample was randomly selected for cases with multiple samples (**Supplementary Table S1B**). The cohort containing one organoid sample per patient will be referred to as the “main organoid cohort” throughout the text. Alterations in the four main PDAC driver genes (17), i.e., *KRAS* activating mutations*, CDKN2A* inactivation, *TP53* mutations and *SMAD4* inactivation, were observed with frequencies of 98%, 86%, 84% and 51%, respectively, in the main organoid cohort (**Supplementary Fig. S1, Supplementary Table S1C**), which is in line with previous reports for PDAC tumors and organoids (10,12,14,18). *KRAS* mutations were observed in 50 out of 51 cases, exhibiting one single nucleotide variant (SNV) per organoid line (**Supplementary Table S1D)**. In all organoids deriving from the same patient, the same SNV was detected. Similarly, for all cases in which the original tumors were analyzed, *KRAS* mutation was identical between tumor and matched PDO. The only *KRAS* wild-type line, 107PO, carried an activating *BRAF* mutation (p.V600_K601delinsE) as presumed driver mutation.

Investigating mutational signatures (19) as traces of mutational mechanisms active in tumors, seven mutational signatures with variable relative contributions were identified in the PDO cohort (**Supplementary Fig. S2A**). The age-related signatures AC1 and AC5 were the major relative contributors in 66% (33/50) and 14% (7/50) of the samples, respectively. Signature AC3, related to homologous recombination repair (HRR) deficiency, was the predominant signature in 12% (6/50) of the samples. Organoid line 064PO showed predominantly signature AC3 (HRR) due to a germline *BRCA2* mutation (p.E1953X) and concomitant loss of the wild-type allele, while 76PO exhibited mostly signature AC6, consistent with a microsatellite instability phenotype due to a germline frameshift deletion and loss of heterozygosity in the mismatch-repair gene *MSH2* (Lynch syndrome).

IntOGen mutation platform (20) was used to identify potential novel driver genes in the main organoid cohort based on SNVs and small insertions and deletions (indels) (**Supplementary Table S1E**). Beside the four well-known main drivers mentioned above, *ARID1A* was significantly affected by SNVs/indels at a frequency of 12%. Additionally, *DOCK1* (Dedicator of cytokinesis 1; 17% alteration frequency), *SIPA1L1* (Signal Induced Proliferation Associated 1 Like 1; 7% alteration frequency), *ATP2A2* (ATPase Sarcoplasmic/Endoplasmic Reticulum Ca2+ Transporting 2; 4.7% alteration frequency) and *LRBA* (LPS Responsive Beige-Like Anchor Protein; 7% alteration frequency) were identified as potential drivers by Oncodrive-FM (q-value < 0.1) based on analysis of functional impact bias (21).

### The transcriptional PDAC subtypes are conserved in PDOs and have prognostic value

In order to assess whether our PDO catalogue was able to maintain and reflect the subtype identity of the parental tumors, we performed transcriptional subtyping of organoids and a subset of matching tumors. We focused on the classical and basal-like gene signatures described by Moffitt et al. (8), which have been shown to be relatively independent of tumor cellularity (10). Based on the notion that transcriptional subtypes may not be binary, but represent a transcriptional continuum, with classical and basal-like programs co-existing as different tumor subpopulations within the same tumor (22), non-negative matrix factorization (NMF) was utilized to extract the exposures of each sample to both signatures, which were then assigned as classical and basal-like scores (**Fig. 1C**, **Supplementary Table S2A**). Subtype scores correlated well between 14 organoids-tumor pairs, although the organoid samples generally displayed slightly higher classical and slightly lower basal-like scores than tumor samples (**Fig. 1D**). Similarly, subtype scores at early- and late-passages (8-16 passages, i.e., 2-6 months later) of organoid lines were highly consistent, however a slight tendency to higher classical and lower basal-like scores could be observed at later passages (**Fig. 1E**). We hypothesize that this tendency is probably due to the culture conditions, which might either favor the growth of classical subtype tumor cells or push the expression profile to a more epithelial progenitor phenotype.

A principal component analysis (PCA) of the 1000 most variable genes in PDOs shows that the subtype scores correlated with the first principal component, corresponding to most of the observed transcriptional variation (**Supplementary Fig. S3A** and **B**). We additionally performed an independent component analysis (ICA) to decompose PDO transcriptional heterogeneity into 10 robust, reproducible, and biologically meaningful independent components (**Supplementary Table S2B**). Independent component 9 (IC9) showed a strong positive correlation with the classical score (two-sided Spearman correlation test, correlation coefficient = 0.68, P = 2.9 x 10^-7^, power = 0.999). Likewise, gene set enrichment analysis (GSEA) showed a significant enrichment of Moffitt’s classical signature genes among IC9 top genes (**Supplementary Fig. S3C** and **D**). This suggests that PDAC subtypes are associated with a large-scale transcriptional program, supporting the indication by the PCA analysis of PDO transcription.

For categorical analyses a score threshold (see Materials and Methods) was applied to classify the PDOs into three subtypes: classical (63%), intermediate (21%) and basal-like (16%) (**Fig. 1C**, **Supplementary Table S2A**). Different organoid lines derived from the same patient in general had consistent subtype scores and classification, even for cases with samples derived from primary and metastatic sites (**Fig. 1C**). The only exception was found for case 077, for which two organoid lines were identified as basal-like while one was assigned as intermediate. Remarkably, despite the small sample size, this subtype classification of PDOs was prognostic for primary presentation/chemo-naïve cases: patients with basal-like organoids had significantly shorter survival time than patients with classical organoids (**Fig. 1F**). This result is consistent with previous reports based on PDAC tumor tissues (6, 8) and supports the validity of both the PDO technology as representative model for human PDAC and the method used for PDO subtyping.

Comparing RNA expression profiles between classical and basal-like organoids revealed 2175 differentially expressed genes (DEGs; adjusted *P* < 0.1), of which 1092 were upregulated and 1083 were downregulated in basal-like organoids (**Supplementary Fig. S3E**). Functional enrichment analysis for ENCODE/ChEA consensus transcription factor targets and MSigDB Hallmark collections revealed that genes with higher expression in classical organoids were enriched in target genes associated to the endodermal transcription factor HNF4A while genes overexpressed in basal-like PDOs were enriched in TP63 targets (**Fig. 1G**). HNF4A, a proposed determinant of the classical subtype (23) and TP63, previously identified as the master regulator of PDAC squamous subtype (24) were also differentially expressed between the subtypes (**Fig. 1H**). TNFα signaling via NFκB, inflammatory response, KRAS signaling and epithelial-to-mesenchymal transition (EMT) were among the top Hallmark gene sets enriched in genes upregulated in basal-like organoids (**Supplementary Fig. S3F**).

Taken together, these results indicate that subtype identity is highly preserved and remains stable in organoid cultures and that the major transcriptional features previously described in tumor tissues for classical and basal-like subtypes are recapitulated in the organoid *in vitro* system.

### Genomic representability and heterogeneity of PDAC-PDOs

We next investigated whether our PDAC-PDO biobank remains genomically representative of primary tumors and their heterogeneity. For this, a subset of parental tumors with a tumor cell content >20% was selected for WGS to perform comparative analyses, also including cases with multiple samples, such as case 083 (**Fig. 2A-F**, **Supplementary Data**). Additionally, for a subset of organoid lines, a second sample from a later time point in culture (8-16 passages, i.e., 2-6 months later) was subjected to WGS (**Supplementary Table S1B, Supplementary Data**).

**Figure 2.**
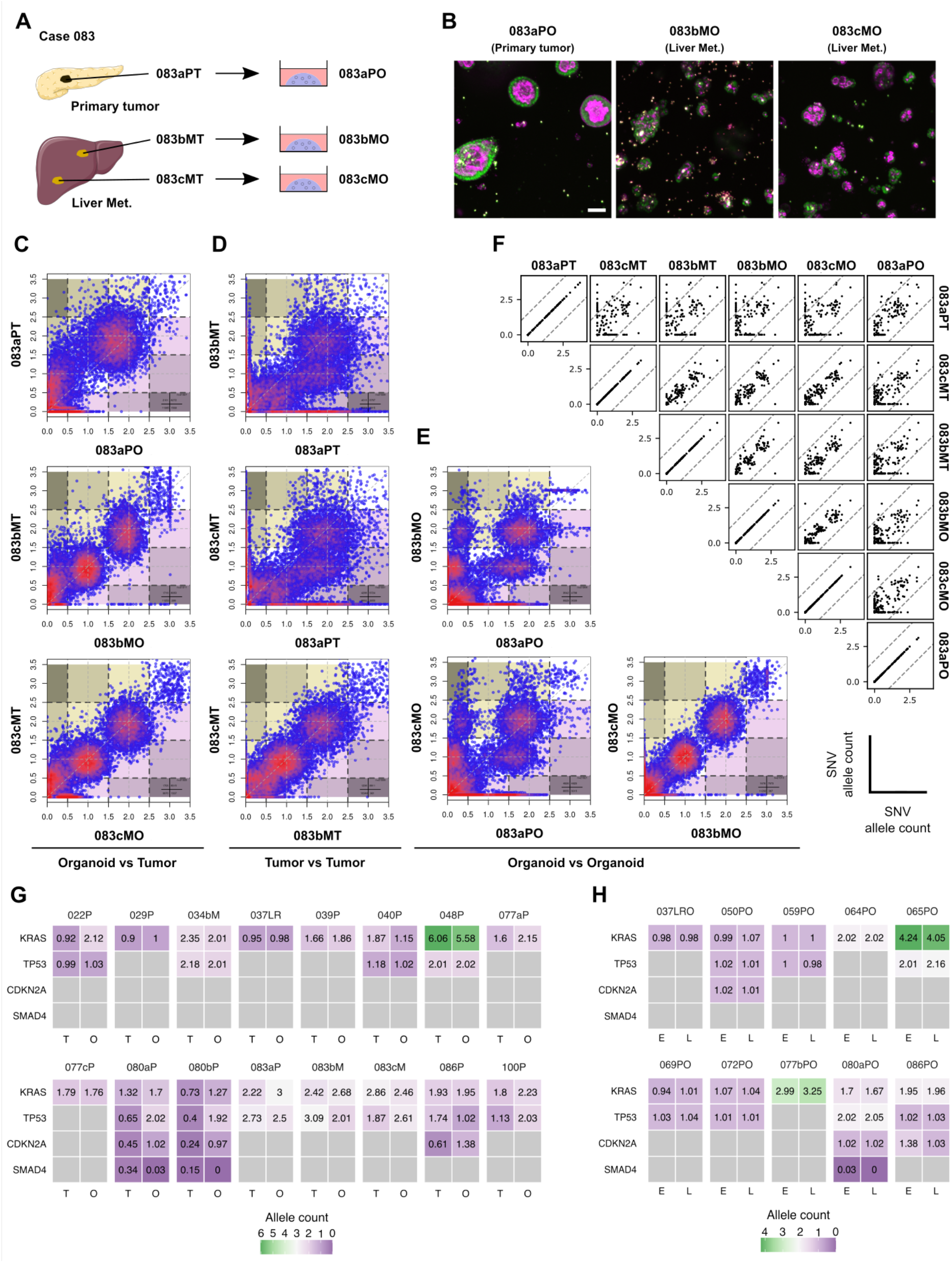
PDAC-PDOs retain the main genomic characteristics of their parental tumors. **A-F,** Analyses for representative case 083. **A,** For patient 083, three organoids lines were generated: one from the primary tumor (083aPO), one from a liver metastasis (083bMO) and one from a second liver metastasis (083cMO). **B,** Representative confocal images of living organoids embedded in Matrigel stained with Hoechst (green) and SiR-Actin (pink) after nine days in culture. Lines 083aPO, 083bMO and 083cMO differ in size and morphology. **C-E,** Correlations of predicted counts for all mutated alleles between each organoid line of case 083 and their corresponding primary tumor/metastatic tissue (**C**), primary tumor and metastatic tissues (**D**) and each organoid line of case 083 (**E**). **F,** Correlations for predicted counts of functional mutated alleles for all sample pairs of case 083. **G,** Comparison of predicted counts of driver gene mutated alleles for parental tumor/metastatic tissues (T) and organoid lines (O). **H,** Comparison of predicted counts of driver gene mutated alleles for the same organoid lines sequenced at different passages (E: early, L: late).

Mutational signature exposures as well as total number of structural variants (SVs) were consistent in organoid cultures compared with their parental tumor samples and remained stable over time in most samples (**Supplementary Fig. S2** and **S4**). Exceptions included case 077c, for which the mutational signature AC13 was particularly enriched in the organoid line (**Supplementary Fig. S2B**), and case 003, for which the number of deletions detected in the early-passage organoid line were notably higher than in the tumor tissue (**Supplementary Fig. S4B**) but decreased again for the later passage sample (**Supplementary Fig. S4C**).

We next focused our comparative analysis on predicted allele counts for non-silent SNVs and indels (**Supplementary Data**). A detailed representative example of the multiple sample case 083 with three organoid/tumor tissue pairs is shown is **Fig. 2A-F**. Comparison of the predicted allele counts for all non-silent (**Fig. 2C**) or functional (**Fig. 2F**) SNVs between organoids and their corresponding tumor tissues revealed that, while the overall genomic representation is well preserved, differences in copy numbers for a fraction of SNVs hints to sub-clonal expansion in organoid cultures. For 083, the observed heterogeneity between primary tumor and metastases was higher than between the two metastases (**Fig. 2D**), which was also recapitulated in the corresponding organoids (**Fig. 2E**). For all tumor-organoid pairs compared, predicted allele counts for recurrently altered PDAC driver genes were usually consistent (**Fig. 2G**). Major discordances between tumors and matching organoids were found only for case 080. While the primary tumor biopsies (080aPT/080bPT) displayed heterogeneity for the driver genes, the matched organoid lines (080aPO/080bPO) did not. Comparison of early and late PDO pairs (**Fig. 2H**) indicated that alterations in driver genes were indeed highly stable during culture.

These results indicate, as may be expected, some variable degrees of sub-clonal expansion during organoid establishment and culture. However, the main functional genomic features of their parental tumors are recapitulated in the PDOs in our cohort and preserved during cultivation.

### Landscape of genomic copy number alterations in PDAC-PDOs

Although the understanding of PDAC genomic features has greatly improved over the past years, a deeper characterization of the copy number landscape remains challenging, mainly due to the reduced sensitivity in the detection of copy number alterations (CNA), in particular deletions, caused by the predominant non-neoplastic components in bulk PDAC tumor tissues. Given that organoid cultures are exclusively composed of epithelial cancer cells, they offer a highly sensitive platform for the characterization of CNAs and for the identification of the underlying drivers.

We investigated the CNA profile of the main organoid cohort, as well as the profile of classical and basal-like organoids separately. Significant chromosomal arm-level aberrations included amplification of 8q (50%) and deletion of 17p (80%), 18q (72%) and 9p (70%) (**Fig. 3A, Supplementary Table S3A**). While these aberrations have been previously reported for PDAC (9, 10) our organoid cohort suggests higher frequencies of occurrence for these chromosome arms.

**Figure 3.**
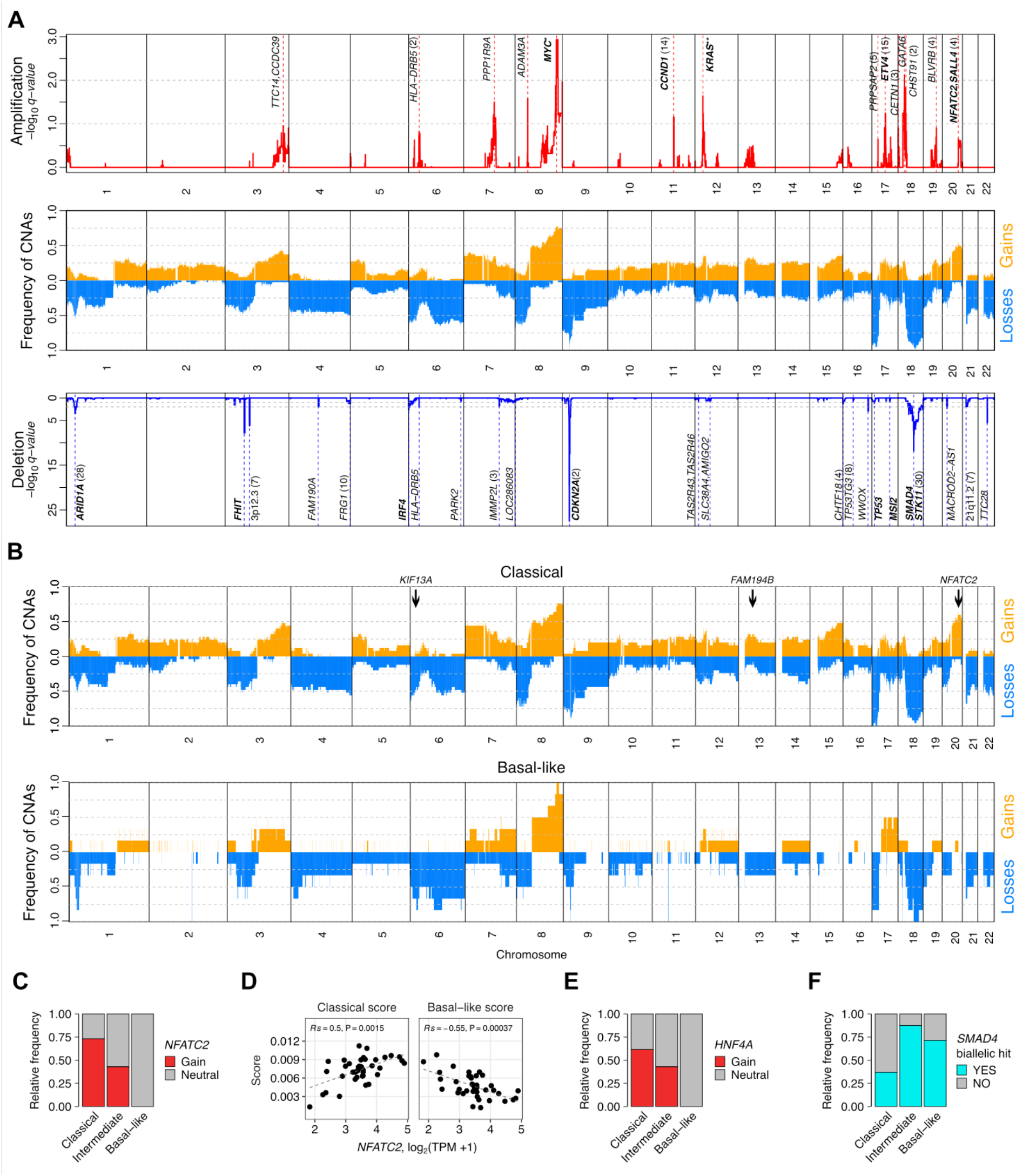
Landscape of genomic copy number alterations in PDAC organoids. **A,** Frequency of somatic copy number gains (orange) and copy number losses (blue) predicted by GISTIC2.0 in the main organoid cohort (mid panel). Significant amplification (top panel) and deletion (bottom panel) peaks (q-value < 0.25). Known oncogenic driver genes are annotated in bold. **\*** The direct hit of the amplification peak on chromosome 8q24.21 was an intergenic locus between *MYC* and *PVT1*. **\*\*** The direct hit of the amplification peak on chromosome 12p12.1 was an intergenic locus close to a cluster of genes including *BCAT1*, *LRMP*, *LYRM5* and *KRAS*. **B,** Frequency of somatic copy number alterations in classical and basal-like organoids. Differential (*NFATC2*) and unique significant amplification peaks for classical organoids (*KIF13A* and *FAM194B*) are indicated. **C,** Fraction of samples carrying *NFATC2* copy number gains, as predicted by ACEseq, according to organoid subtype. **D,** Correlation of *NFATC2* expression with classical and basal-like scores. TPM: transcript per million. **E,** Fraction of samples carrying *HNF4A* (20q13.12) copy number gains, as predicted by ACEseq, according to organoid subtype. Correlation of *HNF4A* expression with classical and basal-like scores appears in Fig. 1I. **F,** Relative frequency of deleterious *SMAD4* alteration by organoid subtype.

In our main organoid cohort, 28 significant deletion and 14 significant gain peaks were identified (**Fig. 3A, Supplementary Table S3B** and **C**). The most significantly amplified peak mapped to the intergenic locus between *MYC* and *PVT1* on 8q24.21 with a frequency of 80%, whereas the most significant deletion peaks were *CDKN2A* (9p21.3) and *SMAD4* (18q21.2), present in 98% and 95% of the samples, respectively. Several of the significant regions contained well-known oncogenes (such as *CCND1* and *ETV4*) or tumor suppressors (such as *CDKN2A*, *SMAD4* and *STK11*), for which the expression levels correlated with CNA status (**Supplementary Fig. S5** and **S6**).

To gain insight of potential genomic alterations contributing to transcriptional subtype determination, we compared copy number patterns in classical and basal-like organoids (**Fig. 3B**) and sought out significant CNAs displaying differential frequencies between the subtypes. We found a significant exclusion of copy number gains for the amplification peak on 20q13.2 in basal-like organoids (Fisher’s exact test, *P* < 0.005) (**Fig. 3C**). Among the four genes located in the amplification peak (*NFATC2*, *ATP9A*, *SALL4*, *MIR3194*), the transcriptional factor and epigenetic modulator *NFATC2* was found to be differentially expressed between classical and basal-like organoids (adjusted *P* < 0.05). Additionally, *NFATC2* expression correlated negatively with the basal-like subtype score (**Fig. 3D**). Although involvement in PDCA tumor growth and progression has been previously shown for *NFATC2* (25, 26), a direct role of *NFATC2* in subtype determination has not been reported yet. Nevertheless, another probable driver candidate of the differential CNA in this region is the classical transcription factor *HNF4A* (**Fig. 1G-I**), located 7 Mb away on 20q13.12, which also displayed exclusion of copy number gains in basal-like organoids (**Fig. 3E**). Additionally, when analyzing only classical PDOs (**Supplementary Fig. S7, Supplementary Table S3D-F**), we identified significantly amplified regions in 6p22.3 (*KIF13A*), a region frequently gained in retinoblastoma (27), and in 13q14.13 (*FAM194B*), both of which have not been associated with PDAC yet. These observations identify potential candidate genes underlying subtype identity according to the Moffitt transcriptional subtypes.

As expected, we observed a high frequency of SMAD4 copy number losses in the main organoid cohort. However, complete loss of *SMAD4,* due to homozygous deletions or mutations combined with monoallelic losses, was significantly under-represented in classical organoids (Fisher’s exact test, *P* < 0.01) (**Fig. 3F**). This result contradicts a recent report linking decreased frequency of intact *SMAD4* to classical tumors (22). Our result is yet in line with a higher *SMAD4* expression observed in classical PDX tumors by Moffitt et al. (8).

### *In vitro* chemosensitivity of PDAC-PDOs is associated with patient survival

We used the DeathPro workflow (28), a confocal imaging-based viability assay that is able to resolve drug-induced cell death and proliferation inhibition, to evaluate the sensitivity of 39 PDOs to the cytostatic and cytotoxic effects of six chemotherapeutic agents commonly used in the treatment of PDAC patients (**Fig. 4A** and **B, Supplementary Table S4A**). Death and proliferation inhibition induced by gemcitabine, 5-fluorouracil (5-FU), irinotecan, oxaliplatin, paclitaxel and erlotinib were determined at two different timepoints: after 72h of drug treatment and after another 72h of drug washout (144h). Area under the curve (AUC) values derived for death (AUCd) and proliferation inhibition (AUCpi) were used to analyze response. Organoid lines displaying AUC values below 0.2 were considered as non-responders.

**Figure 4.**
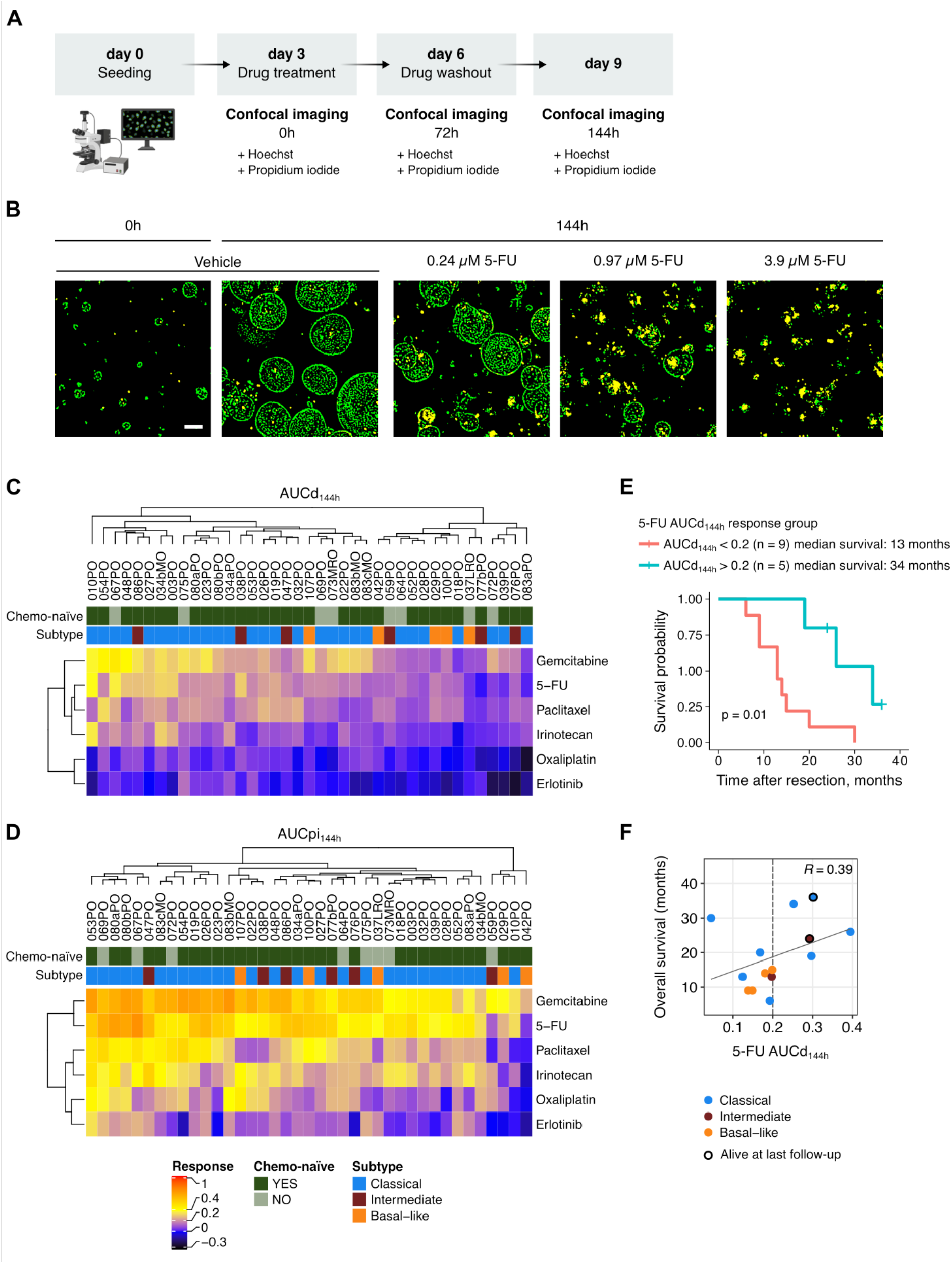
Organoid cytotoxic response to 5-FU treatment is associated with patient survival after adjuvant therapy. **A,** Illustration of the DeathPro assay protocol used for screening PDAC-PDO responses to the clinically relevant chemotherapeutic drugs gemcitabine, 5-FU, irinotecan, oxaliplatin, paclitaxel and erlotinib. Drug-induced cell death and proliferation inhibition were evaluated by confocal imaging after 0h, 72h and 144h (after drug washout) of treatment beginning. **B,** Exemplary DeathPro workflow confocal images of line 067PO before (0h) and after treatment (144h) with the vehicle or with increasing concentrations of 5-FU. Merged Hoechst (green) and propidium iodide (red) channels of maximum intensity projections are shown. Scale bar: 100 µm. **C-D,** Hierarchical clustering of the 39 screened organoid lines according to death (AUCd_144h,_ C) and growth inhibition (AUCpi_144h,_ D) responses. **E,** Kaplan-Meier curves (log-rank test) for the overall survival of patients receiving adjuvant therapy containing 5-FU, grouped by AUCd_144h_ response threshold. Patients with 5-FU-responsive organoids had a longer median overall survival (34 months) than patients with non-responsive organoids (13 months). **F,** Correlation of 5-FU AUCd_144h_ with overall survival of patients receiving adjuvant therapy containing 5-FU. Only organoid lines derived from chemo-naïve patients at resection were included in this analysis. Dotted lines denote the response threshold of 0.2 AUC units. Dot color indicates subtype. Black border indicates patients still alive at the last follow-up.

We observed a strong positive correlation between AUCd (Rp = 0.78) and AUCpi (Rp = 0.89) values at these two different timepoints (**Supplementary Fig. S8A** and **B**). Given that responses were more robust and had a wider dynamic range after the drug washout period, we focused on the readouts 44h after treatment begin for further analyses. Results revealed that for all six drugs cytostatic effects were preponderant over cytotoxic effects (**Fig. 4C** and **D**). LD50 values could consequently not be determined for a large proportion of the samples (**Supplementary Fig. S8C**). Gemcitabine and 5-FU were under our screen settings the most effective cytostatic and cytotoxic drugs for most of the PDOs, followed by paclitaxel **(Fig. 4C** and **D**). Oxaliplatin and erlotinib were particularly inefficient in inducing cell death, therefore only AUCpi values were included in further analyses for these two drugs. In general, we observed correlation between responses of individual organoid lines to different drugs (**Supplementary Fig. S8D**), suggestive of the presence of multidrug resistance mechanisms in PDAC. In particular, strong correlations (Rs > 0.7) were found between 5-FU and paclitaxel AUCpi_144h_ as well as between 5-FU and gemcitabine AUCd_144h_.

To assess whether our PDAC-PDO drug screening platform had clinical predictive value, we correlated the organoid drug sensitivity data with overall survival of the patients receiving the corresponding adjuvant therapy. A total of 27 out of 35 patients in our cohort with relevant follow-up information received adjuvant treatment, most commonly (78%) gemcitabine alone or in combination with 5-FU (or its prodrug capecitabine) (**Supplementary Table S4B**). We analyzed the correlation of organoid drug responses to gemcitabine and 5-FU with the overall survival time of patients that received gemcitabine or 5-FU, respectively, as monotherapy or combined adjuvant treatment after upfront resection. Interestingly, when stratified by PDO 5-FU AUCd_144h_ response, the median overall survival for patients with responsive organoids (AUCd_144h_ > 0.2) was significantly higher than for patients with non-responsive organoids (**Fig. 4E**).

The association of PDO sensitivity with clinical outcome was not solely determined by the transcriptional subtypes. While we observed that basal-like organoids were non-responders to the cytotoxic effects of 5-FU and were associated with shorter patient survival, the response of classical organoids was heterogenous, as was the clinical outcome (**Fig. 4F**). Similarly, 5-FU AUCd_144h_ did not correlate significantly with the subtype scores (**Supplementary Fig. S9A**), although a moderate correlation (Rp > 0.5) was found between classical score and 5-FU AUCpi_144h_ (**Supplementary Fig. S9B**).

In contrast to 5-FU AUCd_144h_, PDO response to gemcitabine treatment did not correlate with survival nor did 5-FU AUCpi_144h_ (**Supplementary Fig. S9C-E**). Collectively, these results suggest that organoid pharmacotyping offers an effective means to identify patients more likely to benefit from 5-FU-based adjuvant treatment.

### Global transcriptomics programs underpin drug responses in PDAC-PDOs

In order to relate drug response of PDAC-PDOs to transcriptomic features, we derived drug-response-specific gene sets by identifying genes whose transcriptional levels correlated positively or negatively with AUCd_144h_ or AUCpi_144h_ parameters using Spearman correlation analysis (*P* < 0.01; **Supplementary Table S5**). As for patient survival analysis, we included only organoids derived from chemo-naïve patients in order to avoid confounders arising from selection pressure by previous chemotherapy. For simplification, gene sets were stratified into two groups based on the directionality of correlation, i.e., genes with lower expression in more resistant organoids are referred to as “down-in-resistant”, while genes with higher expression in more resistant organoids are referred to as “up-in-resistant”.

At the gene level, we observed varying but generally modest overlap among gene sets (**Supplementary Fig. S10A** and **B**). Most of the overlaps occurs among gene sets derived from the same parameter, i.e., either AUCd_144h_ or AUCpi_144h_, suggesting that different genes are associated with cytotoxic and cytostatic responses. In the case of AUCd_144h_, the largest gene overlaps were observed between gemcitabine/irinotecan and gemcitabine/5-FU for both up-in-resistant (**Supplementary Fig. S10A**) and down-in-resistant gene sets (**Supplementary Fig. S10B**), which is intelligible since the mechanism of actions of these three drugs involve direct interference of DNA synthesis. A small number of genes were shared by gene sets related to at least 3 different drugs, which might therefore be implicated in multidrug resistance mechanisms (**Fig. 5A**). Among genes shared by up-in-resistant gene sets, we found genes related to processes with potential therapeutic implications such as mitochondrial protein homeostasis (29) (*SPG7* Matrix AAA Peptidase Subunit, Paraplegin), sphingolipid biosynthesis (30) (*SPTSSA*, Serine Palmitoyltransferase Small Subunit A) and endoplasmic reticulum homeostasis (31) (*H6PD*, Hexose-6-Phosphate Dehydrogenase/Glucose 1-Dehydrogenase).

**Figure 5.**
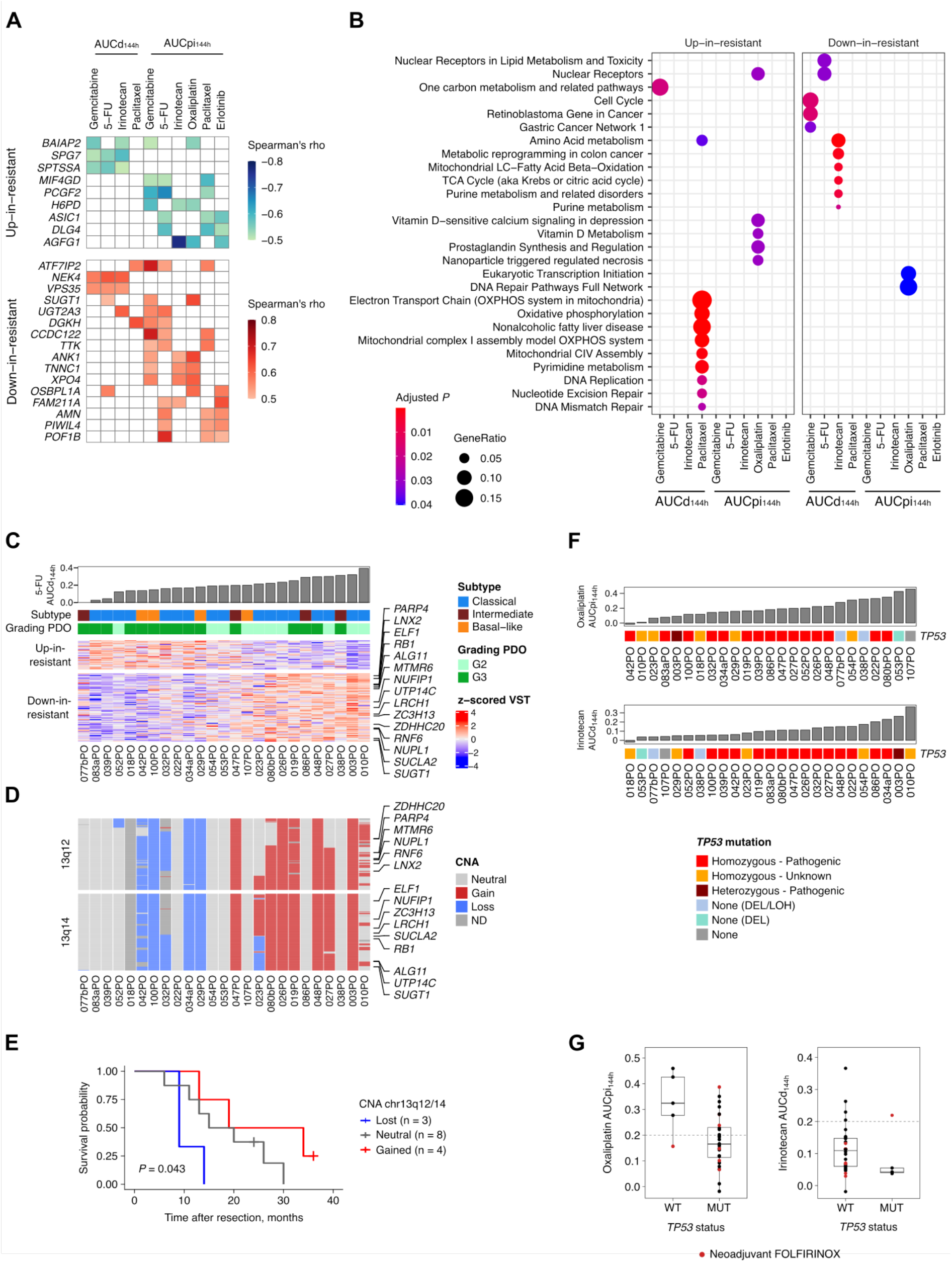
Pharmacotranscriptomic and pharmacogenomic associations revealed by PDO drug screening. **A,** Genes whose expression correlates negatively (up-in-resistant) and positively (down-in-resistant) with the response (AUCd_144h_ or AUCpi_144h_) to at least 3 different drugs. **B,** Significantly enriched Wikipathways terms (adjusted *P* < 0.05) for each gene-drug response correlation gene set. **C,** Heatmap of gene expression correlating negatively (up-in-resistant) or positively (down-in-resistant) with the cytotoxic effect of 5-FU (AUCd_144h_). Samples are sorted by 5-FU AUCd_144h_ and genes are sorted by their Spearman’s correlation coefficient. A significant enrichment (adjusted *P* < 0.0001) of genes located on chromosome 13q12 and 13q14 (annotated genes) was observed for 5-FU AUCd_144h_ down-in-resistant gene set. **D,** Copy number alterations for the enriched cytobands in **C**, 13q12 and 13q14. Samples are sorted as in **C** and genes are sorted by their position along the chromosome. Annotated genes belong to the 5-FU AUC_144h_ down-in-resistant gene set (same as in **C**). **E,** Kaplan-Meier curves (log-rank test) for overall survival of patients receiving adjuvant therapy containing 5-FU, stratified by CNA status for chr13q12/14 in organoids. **F,** Oncoprints of *TP53* mutational status (SNVs/indels) for chemo-naïve PDOs included in the drug screen. Mutation zygosity and functional impact prediction as reported by COSMIC (FATHMM prediction) are indicated. For non-mutated samples, the presence of other alterations (DEL: deletion, LOH: loss of heterozygosity) are reported. Samples are sorted by oxaliplatin AUCpi_144h_ (top panel) and irinotecan AUCd_144h_ (bottom panel). *TP53* wild-type status (n = 4) was associated with increased sensitivity to oxaliplatin proliferation inhibition in chemo-naïve PDOs (top third, Fisher’s exact test, *P* < 0.005) and lower sensitivity to irinotecan cytotoxic effects (bottom third, Fisher’s exact test, *P* < 0.005). **G,** Proliferation inhibition (AUCpi_144h_) induced by oxaliplatin and death induced by irinotecan (AUCd_144h_) in PDOs. PDOs are grouped by *TP53* mutation status. Organoids derived from patients pre-treated with neoadjuvant FOLFIRINOX are indicated.

Surprisingly, functional enrichment analysis for Wikipathways and KEGG collections revealed distinct significantly enriched pathways for each gene set, with almost no overlap (**Fig. 5B** and **Supplementary Fig. S11**) in spite of the moderate or strong correlations in the responses for some of the drugs (**Supplementary Fig. S8D**). Enriched pathways in drug response specific gene sets included terms associated with oxidative phosphorylation, metabolic reprogramming, one carbon metabolism and vitamin D metabolism, all of which have been postulated as potential target for therapeutic approaches in cancer (32–34).

In order to ascertain if any large-scale transcriptional program was associated with drug response we investigated if the independent components obtained from ICA were enriched in any of the gene sets. Interestingly, IC4 was strongly associated with irinotecan AUCd_144h_ gene sets. IC4 was positively enriched for up-in-resistant irinotecan AUCd_144h_ genes while negatively enriched for down-in-resistant irinotecan AUCd_144h_ genes (**Supplementary Fig. S12A**). While the top 100 genes contributing to IC4 were mainly enriched for ribosomal proteins (**Supplementary Fig. S12B**), the bottom 100 genes in IC4 displayed enrichment for metabolic reprograming pathways, comparable to the down-in-resistant gene set enrichment analysis (**Supplementary Fig. S12C**). Remarkably, the top 2 genes of IC4 were to the Mitochondrially Encoded Cytochrome C Oxidase III (*MT-CO3*) and the Mitochondrially Encoded NADH:Ubiquinone Oxidoreductase Core Subunit 1 (*MT-ND1*) genes, both integral components of the mitochondrial respiratory chain. These findings suggest the existence of an oxidative phosphorylation-related transcriptional program playing a complex role in chemoresistance, alongside other transcriptional programs.

For 5-FU AUCd_144h_, which showed relevance for prediction of clinical response in our cohort, no pathways were found to be over-represented in the up-in-resistant gene set, which was the smaller one with only 30 genes, while significantly enriched pathways in the down-in-resistant gene set were limited to “Nuclear Receptors in Lipid Metabolism and Toxicity” and “Nuclear Receptors” (**Fig. 5B** and **Supplementary Fig. S11**). Notably, in the down-in-resistant gene set for 5-FU AUCd_144h_ we observed an enrichment of genes located in chromosome 13q14 (adjusted *P* < 1 x 10^-7^) and 13q12 (adjusted *P* < 1 x 10^-4^), among them the tumor suppressor *RB1* (**Fig. 5C**). In addition, an association at the genomic level was also observed for these regions; monoallelic losses were more frequent in more resistant organoids, while gains were more frequent in more sensitive organoids (**Fig. 5D, Supplementary Fig. S13**). Moreover, chr13q12/14 loss was associated with shorter overall survival for patients receiving adjuvant therapy containing 5-FU (**Fig. 5E**). These results suggest that genes located in these chromosome 13q regions may play a key role in chemoresistance, particularly to 5-FU.

### Wild-type *TP53* status is associated with higher sensitivity to growth inhibition induced by oxaliplatin

Altered *TP53* has been linked to worse overall survival in PDAC (35), yet it is a potential positive predictive factor for gemcitabine efficacy in the adjuvant setting (36). In our main organoid cohort, 83% of the PDOs had mutated *TP53*. Only seven organoid lines carried wild-type *TP53*, five of which were included in the drug screen. Although we did not observe an association with gemcitabine response, wild-type *TP53* was associated with higher sensitivity of PDOs to the cytostatic effect of oxaliplatin (top third, Fisher’s exact test, *P* < 0.005; **Fig. 5F**). Interestingly, these organoid lines also showed low sensitivity to the cytotoxic effects of irinotecan (**Fig. 5F)**. Only one wild-type *TP53* organoid line, 067PO, was a non-responder to oxaliplatin treatment but responded to irinotecan (**Fig. 5G**). This PDO derived from a patient who received the oxaliplatin- and irinotecan-containing FOLFIRINOX regime in a neoadjuvant setting, hinting to selection or development of resistance against oxaliplatin in pre-treated tumors. Unfortunately, none of the patients with wild-type *TP53* organoids received oxaliplatin-containing adjuvant therapy and therefore we were not able to clinically evaluate this finding. Nevertheless, our results suggest that wild-type status of *TP53* is a potential positive predictive factor for oxaliplatin-based chemotherapy efficacy although this observation needs clinical validation. This might have important translational implication since modified FOLFIRINOX has become one of the standard-of-care regimens for palliative, neoadjuvant and adjuvant therapy in pancreatic cancer (37) but up to date, no predictive biomarkers of response have been identified.

### Added value of functional in vitro drug sensitivity testing in PDOs for personalized oncology and target validation

One of the most promising applications of PDOs is their exploitation as personalized oncology tools to identify therapeutic vulnerabilities and functionally test targeted therapies. Within this context, in addition to the sensitivity screen for chemotherapeutics in standard clinical use, we tested whether homologous recombination deficiency (HRD) score, *CDKN2A/B* alteration status and *TP53* loss of individual organoid lines were able to predict response to the PARP inhibitor olaparib, to the CDK4/6 inhibitor palbociclib and to the RNA polymerase II inhibitor α-amanitin, respectively. For this, as a proof of concept, eight PDAC-PDOs were selected based on these genomic features and were stratified accordingly into control or target groups (**Fig. 6A** and **B**).

**Figure 6.**
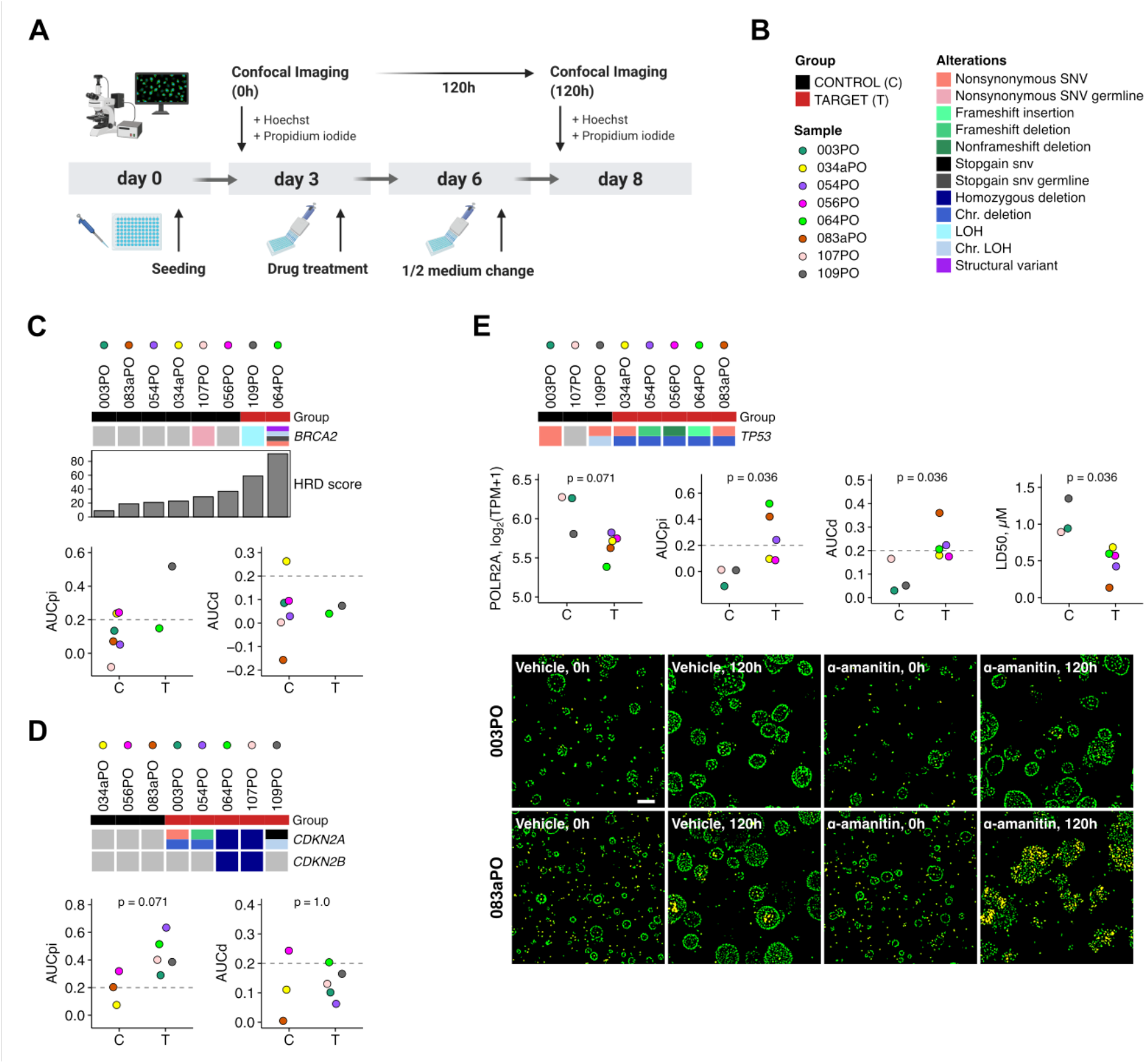
PDOs as platform for testing personalized treatments: PDAC organoids harboring TP53/POLR2A co-deletion display increased sensitivity to RNA polymerase II inhibition by α-amanitin. **A,** Illustration depicting the DeathPro assay protocol for assessing organoid response to the molecular targeted agents olaparib, palbociclib and α-amanitin. **B,** Legends for subfigures **C** to **E** depicting the color codes for control and target groups, individual organoid lines and genomic alterations. **C,** Response of PDCA-PDOs to the PARP inhibitor olaparib. Organoid lines were stratified by their homologous recombination deficiency (HRD) score (middle panel) into control (black) and target (red) groups for olaparib treatment. Alteration status of *BRCA2* is shown (top panel). Comparison of AUCpi and AUCd values between control (C) and target (T) groups (bottom panel), computed from the dose-response curves in **Supplementary Fig. S14A. D,** Response of PDCA-PDOs to the CDK4/6 inhibitor palbociclib. *CDKN2A/B* alteration status (top panel) was used to classify organoid lines into control (black) and target (red) groups for palbociclib treatment. Comparison of AUCpi and AUCd values (bottom panel) between control (C) and target (T) groups, derived from the dose-response curves in **Supplementary Fig. S14B**. *P* values derive from Wilcoxon signed-rank test. **E,** Response of PDCA-PDOs to RNA polymerase II inhibition by α-amanitin. Organoid lines with *TP53* neutral copy number or heterozygous deletion were stratified into control (black) and target (red) groups for α-amanitin treatment, respectively (top panel). Comparison of *POLR2A* expression, AUCpi, AUCd and LD50 values (middle panel) between control (C) and target (T) groups. P values derive from Wilcoxon signed-rank test. Dose-response curves are shown in **Supplementary Fig. S14C**. DeathPro workflow confocal images of a control (003PO) and target organoid line (083aPO) before (0 h) and after treatment (120 h) with the vehicle (water) or 200 nM α-amanitin. Merged Hoechst (green) and propidium iodide (red) channels of maximum intensity projections are shown. Scale bar: 100 µm (bottom panel).

Two lines with a HRD score above 37.8, corresponding to the upper quartile of the main organoid cohort, were stratified as potential responders (target group) to olaparib treatment (**Fig. 6C, Supplementary Fig. S14A**). Notably, only one of the target lines showed high sensitivity to olaparib-induced growth inhibition, while the organoid line with the germline BRCA2 mutation (064PO) was not responsive. Of note, 064PO was derived from a patient who received FOLFIRINOX in a neoadjuvant setting and therefore, the lack of response to olaparib might be explained by development of resistance mechanisms during therapy with oxaliplatin. Olaparib did not induce cytotoxicity in the target lines, although one of the lines in the control group evidenced cell death response. This result suggests that genetic aberrations in fundamental HRD genes might not be sufficient evidence to inform PARP inhibition treatment and that addition of organoid-based functional testing could provide relevant information for therapeutic decisions.

All the tested organoid lines with defective *CDKN2A* were responders to growth inhibition induced by the CDK4/6 inhibitor palbociclib, although cell death response was scarce in both groups (**Fig. 6D, Supplementary Fig. S14B**). Although the difference in cytostatic response between the target and control groups did not reach statistical significance, these results suggest that *CDKN2A/B* status should be further evaluated in context of palbociclib treatment in PDAC.

Co-deletion of *POLR2A* together with *TP53* has been previously described as a potential therapeutic vulnerability in cancer (38). Colon, prostate and breast tumor cells with *TP53*/*POLR2A* loss are more vulnerable to RNA polymerase II inhibition by the fungal toxin α-amanitin than tumor cells with neutral copy number (38–40). In our main organoid cohort, 79% (34 out of 43) presented heterozygous *TP53* loss. In all of the cases we observed the concomitant deletion of *POLR2A*. Therefore, we evaluated if *TP53* loss was also associated with increased sensitivity to RNA polymerase II inhibition by α-amanitin in PDAC-PDOs. Organoids from the target group (*TP53*/*POLR2A* loss) showed significantly higher sensitivity to α-amanitin growth inhibition and death induction, with a median LD50 for the target group 1.7-fold lower than for the control group (**Fig. 6E, Supplementary Fig. S14C**). These results indicate that heterozygous deletion of *TP53* renders tumor organoids more susceptible to RNA polymerase II inhibition, which could be exploited as a potential therapeutic target for PDAC.

Taken together, the results of the targeted drug screen suggest an additional value of functional *in vitro* drug sensitivity testing in PDAC-PDOs over conventional genome-driven personalized oncology and demonstrate the potential of organoids as a platform to functionally validate novel targeted therapy approaches to PDAC.

## Discussion

In this work, we presented a catalogue of 51 PDAC-PDOs generated from resected tumors with an in-depth molecular characterization based on RNAseq and WGS data. For a subset of 39 PDOs, molecular data was complemented with drug response profiling for standard-of-care chemotherapeutics. A major particularity of our cohort, in comparison to previous studies on PDAC organoid pharmacotyping (12,15,41), is the detailed clinical follow-up information that allowed us to directly relate PDO molecular features and drug response to clinical outcomes.

We employed a customized reduced medium composition in order to maintain serum-free conditions as well as to confine the growth to tumor cells. Failure in PDO establishment due to normal ductal cell overgrow is recognized as a major challenge particularly for PDAC tumors. PDAC tumor organoids show independence to some of the niche factors needed by their normal counterpart (16) as well as heterogeneity in their requirements for successful establishment (14, 15). Nevertheless, we reached an establishment efficiency rate within the previously reported range, despite including patients that received neoadjuvant therapy, which indicates that our culture conditions are not exceedingly selective for a subset of tumors. Importantly, we also showed that our organoid cultures remain molecularly representative of the original tumors.

Another point showcasing the representability of our PDOs is the recognition of the subtype spectrum from classical to basal-like phenotypes. Transcriptional profiles expressed as subtype scores were highly concordant between organoids and original tumors. This indicates that subtype heterogeneity is retained in organoid cultures, as we and others have shown also at the single cell level (42, 43). Strikingly, we were able to observe in our rather small cohort the prognostic potential of tumor subtyping based on organoid transcriptomic data. To the best of our knowledge, this is the first study reporting the association of organoid-based subtyping with patient outcome. These results from PDOs as a pure tumor cell culture also validate the signatures suggested by Moffitt et al. (8) as a feature of the tumor epithelium.

Given promising therapeutic implications (23, 44), it has become increasingly relevant to understand molecular players underlying the classical and basal-like phenotypes. Few copy number alterations have been associated with classical or basal-like tumors, such as *GATA6* (10) or *MYC* amplifications (45), respectively. Our CNA profiling of PDOs revealed exclusion of chromosome 20q13.2 gains in basal-like organoids while their frequency in classical organoids was around 60%. The high tumor purity of organoid cultures also allows to identify concordances between CNAs and mRNA levels, which may point out to oncogenic drivers. In particular, *NFATC2*, a known cancer driver located on the chromosome 20q13.2 amplification peak, was upregulated in classical organoids. We were also able to link frequent gains of the classical transcription factor *HNF4A* (23), located on chromosome 20q13.12, to its upregulation in classical organoids. Additionally, our data suggests an association between gains in chromosomes 6p22.3 and 13q14.13 with the classical subtype, which can be further explored.

Organoid drug screening has gained substantial popularity over recent years with the promise of becoming a clinically relevant tool for personalized oncology (46). Seppälä et al. have recently shown that establishment and pharmacotyping of PDAC-PDOs can be completed within a timeframe that would allow to inform adjuvant treatment (41). Previous reports assessing the predictive value of PDAC-PDO pharmacotyping only included a small number of patients with clinical data and focused mainly on advanced disease (12, 15). Given the practical impossibility of directly assessing response to therapy in a post-resection adjuvant setting, we used overall survival as an indicator of clinical response to the adjuvant treatment. Even though all patients receiving adjuvant 5-FU in our cohort were treated with combination regimens for adjuvant chemotherapy, patients with organoids classified as 5-FU-responders had a better outcome than patients with non-responding organoids. Given that the two current standard-of-care regimes in the adjuvant setting contain 5-FU, namely gemcitabine plus capecitabine (5-FU pro-drug) and the modified FOLFIRINOX regime (37), our results are of high clinical relevance and suggest that organoid screening could be used to identify patients for which 5-FU-based adjuvant treatment could be beneficial and patients for which alternative chemotherapy regimens, such as gemcitabine plus nab-paclitaxel (nanoparticle albumin bound paclitaxel), should be pursued. Therefore, prospective validation of PDO pharmacotyping as therapy stratification tool is urgently needed.

Despite an overall correlation in drug responses, integration of drug sensitivity data with transcriptomic profiles recognized distinct pathways associated with the response to the different drugs. Several of these pathways relate to metabolic functional modules including OXPHOS, metabolic reprogramming and one carbon metabolism, suggesting that metabolic plasticity might play a crucial role in PDAC drug resistance. Our drug screen also identified copy number alterations in chromosome 13q as a potential predictive marker of response to 5-FU as well as wild-type *TP53* as a potential predictive marker of sensitivity to oxaliplatin. Mutated TP53 has been previously reported to predict gemcitabine efficacy in PDAC patients (36). Together with our results, this suggests that patients with wild-type *TP53* could therefore benefit from oxaliplatin-based adjuvant therapy, such as the FOLFIRINOX regime, instead of a gemcitabine-based treatment.

Despite great efforts made in individual genomic profiling for developing targeted therapies for solid tumors including PDAC (47, 48), still only a small number of patients benefit from such approaches (49). Reasons include lack of predictive biomarkers and limited predictivity of single genomic features to complex response to targeted therapeutics. Hence, complementation of genomic information with functional testing using PDOs might improve response prediction and biomarker identification (50). HRD is associated with a better prognosis in PDAC (35) and patients with BRCA1/2 alterations have been shown to benefit from PARP inhibition with olaparib, although not all of these patients respond to treatment (51, 52). Our proof-of-concept stratification strategy based on genomic features of eight PDOs evidenced, however, that a deleterious BRCA2 alteration and concomitant HRD was not predictive of olaparib sensitivity in an organoid line derived from a FOLFIRINOX pre-treated tumor. Using the same approach, our results suggest that *CDKN2A/B* status could be a potential predictor of sensitivity to the CDK4/6 inhibitor palbociclib, although a previous report using patient-derived cell lines and xenografts did not find an association of response to palbociclib with genetic alterations (53).

Finally, we could show that PDAC-PDOs with *TP53*/*POLR2A* co-deletions displayed increased sensitivity to RNA polymerase II inhibition by the fungal toxin α-amanitin, a vulnerability initially described for colorectal cancer (38). Due to the high liver toxicity of systemic administration of α-amanitin, clinical use has not been possible yet, however, approaches utilizing antibody-drug conjugates for tumor targeting are being tested in order to overcome toxicity and potentially allow its clinical use (54). Alternatively, targeting other components of the RNA polymerase II transcription machinery might represent a promising therapeutic option for TP53/POLR2A hemizygous tumors (55).

In conclusion, our work demonstrates that PDAC-PDOs are valid surrogate tumor models that not only support molecular characterization of PDAC and investigation of drug resistance mechanisms but also hold potential clinical value as stratification tool for PDAC patients.

## Methods

### PDAC patient cohort

PDAC patients were recruited from the Department of General, Visceral and Transplantation Surgery, Heidelberg University Hospital. The study was approved by the ethical committee of University of Heidelberg (ethic votes 301/2001, 159/2002, S-206/2011, S-708/2019) and was conducted in accordance with the Helsinki Declaration. All patients provided written informed consent prior to acquisition of tissue. Histological confirmation of the tumor entity and grading was performed by an experienced pathologist.

### Establishment and culture of pancreatic tumor organoids

Pancreatic tumor organoid cultures were established using a modified version of the protocol described by Boj et al. (16), with culture conditions adapted to restrict the growth of normal ductal cells. Resected pancreatic tumor specimens were collected in ice-cold basal organoid medium consisting of AdMEM/F12 (Gibco), 2 mM GlutaMAX (Gibco), 10 mM HEPES (Gibco), 1x Primocin (InvivoGen) and processed immediately. Collected tumor tissues were cut in smaller pieces, a portion was snap-frozen and stored at −80°C for DNA and RNA isolation, another portion was fixed in 5% neutral buffered formalin for histological processing and the remaining pieces were used for establishing the organoid culture. For this, the tumor pieces were minced and digested in basal organoid medium containing 1 mg/ml collagenase type IV (Sigma-Aldrich), 100 µg/ml DNase I (AppliChem), 1x B27 (Gibco), 1 mM N-acetylcysteine (Sigma-Aldrich) and 10 µM Y-27632 (Selleckchem) for up to 4 h at 37°C with gentle agitation. The resulting cell suspensions were seeded in growth factor reduced, phenol red-free Matrigel (Corning) and were cultured in organoid growing medium consisting of basal organoid medium supplemented with 1x B27, 1 mM N-acetylcysteine, 10% RSPO1-conditioned medium, 100 ng/ml FGF10 (PeproTech), 100 ng/ml Noggin (PeproTech), 500 nM A83-01 (Tocris) and 10 µM Y-27632. Medium was refreshed every 3-4 days omitting Y-27632. Organoids were routinely passaged by dissociation with TrypLE (Gibco) for 10 min at 37°C. Medium was further supplemented with 50 ng/ml EGF (PeproTech) and, if required, with 50% Wnt3A-conditioned medium only after tumor cell enrichment to avoid overgrowth of normal ductal cells during the initial passages. While Wnt-dependency has been previously reported for a subset of PDAC tumor organoids (14), all our organoid lines were able to grow for at least five passages without the addition of exogenous Wnt ligand, relying only on Wnt signaling activation by R-spondin. For one organoid line (065PO), however, strict Wnt-dependency was evident thereafter. Organoid lines were routinely tested for Mycoplasma contamination using Venor GeM Classic (Minerva Biolabs).

### Histopathological and morphological analysis

Tumor tissue pieces were fixed overnight in 5% neutral buffered formalin and embedded in paraffin. Organoid samples were embedded in low-melting point agarose or HistoGel (Thermo Fisher Scientific) before overnight fixation in 5% neutral buffered formalin followed by paraffin embedding. Routine hematoxylin and eosin staining was performed for 4 µm sections for both organoid and tumor samples. Pathological grading of the tumor tissue adjacent to the source for organoid culture was performed by a pathologist, according to the WHO classification system, 5th edition. Additionally, an analogue organoid grading strategy was applied based exclusively on the amount of gland formation and nuclear features, since none of the organoids showed a significant amount of mucin production via light microscopy.

Tumors were classified according to their predominant growth pattern as glandular, cribriform, micropapillary or solid. Organoid lines were classified according to their lumen structure as monolumen, multilumen or no-lumen organoids. For each line, the frequency of these morphological categories was determined from H&E-staining images, by counting organoids with at least 80 µm diameter. A particular morphology was considered predominant when the difference in frequency was higher than 20%, compared with the second most frequent morphology, otherwise a tie was applied. Predominant morphology of tumor-organoid pairs was considered concordant for the following combinations: glandular-monolumen, cribriform-multilumen and micropapillary/solid-no-lumen.

### Whole-genome sequencing

Prior DNA isolation, organoids were released from Matrigel using Cell Recovery Solution (Corning). Frozen sections of matched tumor tissues were stained with H&E and tumor cell content was evaluated. Only tissue samples with > 20% tumor cell content were selected for whole-genome sequencing (WGS). DNA from organoids and frozen tumor tissues was isolated using the QIAamp DNA Mini Kit (Qiagen) according to the manufacturer’s instructions. DNA from matched blood samples was isolated using the QIAamp DNA Blood Mini Kit or the QIAsymphony DSP DNA Mini Kit (Qiagen). Quality control was performed at the DKFZ-HIPO Sample Processing Laboratory using Quant-iT dsDNA Kit, broad range (Thermo Fisher Scientific) and Genomic DNA ScreenTape on the TapeStation System (Agilent). Library preparation and WGS sequencing was performed at the DKFZ Genomics and Proteomics Core Facility. WGS libraries were prepared using the TruSeq Nano DNA Library Prep Kit (Illumina) and paired-end sequencing (2×150 bp) was performed using the Illumina HiSeq X platform. Tumor tissue, organoid and germline samples were sequenced in four, two and one lanes, respectively.

WGS data from line 028PO was excluded from the analyses because it failed coverage uniformity quality control.

### Whole-genome sequencing analysis

#### Illumina short read alignment

Illumina short reads were aligned using the DKFZ alignment workflow, as used in the ICGC Pan-Cancer Analysis of Whole Genome (PCAWG) projects (56). The workflow is available for download as a Docker container: https://dockstore.org/containers/quay.io/pancancer/pcawg-bwa-mem-workflow. Briefly, bwa mem v0.7.8 was used to align paired reads to the reference human genome sequence (build 37, version hs37d5 with PhiX) with minimum base quality threshold of zero [-T 0] and remaining settings left at default values (57). The alignments were sorted by coordinates using biobambam bamsort (version 0.0.148) with compression option set to fast (1), which was streamed into duplicate read pairs marking using biobambam bammarkduplicates with compression option set to best (9) (58).

#### Somatic small nucleotide variant calling

Somatic single nucleotide variants (SNVs) in matched organoid/tumor-normal pairs were called using the DKFZ SNV calling workflow, as used in the ICGC PCAWG projects. The workflow is available for download as a Docker container: https://dockstore.org/containers/quay.io/pancancer/pcawg-dkfz-workflow:2.2.0. Briefly, the SNVs were identified using samtools and bcftools version 0.1.1957 (59) and then classified as somatic or germline by comparing the tumor sample to the control, and later assigned a confidence which is initially set to 10, and subsequently reduced based on overlaps with repeats, DAC blacklisted regions, DUKE excluded regions, self-chain regions, segmental duplication records as introduced by the ENCODE project (60) and additionally if the SNV exhibited PCR or sequencing strand bias. Only SNVs with confidence 8 or above were considered for further analysis.

#### Somatic small insertion and deletion calling

Somatic small insertions and deletions (indels) in matched organoid/tumor-normal pairs were called using the DKFZ indel calling workflow, as used in the ICGC PCAWG projects. The workflow is available for download as a Docker container: https://dockstore.org/containers/quay.io/pancancer/pcawg-dkfz-workflow:2.2.0. Somatic indels were identified using Platypus (61). Indel calls were filtered based on Platypus internal confidence calls, and only indels with confidence 8 or greater were used for subsequent analysis.

#### Somatic small mutation annotation

The protein coding effect of somatic SNVs and indels from all samples were annotated using ANNOVAR (62) according to Gencode v19 gene annotation and overlapped with variants from dbSNP (build 141) and the 1000 Genomes Project phase 3 database (63, 64).

#### Calculation of allele counts

In order to compare the SSMs between organoids and primary tumor we calculated the allele counts for each variant. The allele count can be interpreted as the number of alleles in the tumor predicted to harbor the variant, modelling the contribution of tumor and diploid normal towards the variant allele fraction (VAF) observed for any given variant. Predicted allele counts were annotated to each variant based on the following formula:

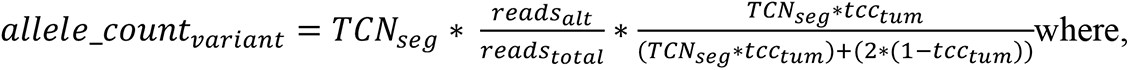

*TCN_seg_* = total copy number of the segment on which the variant lies (as calculated by ACEseq)

*reads_alt_* = the number of reads supporting the variant allele

*reads_total_*= the total number of reads over the position where the variant was called

*tcc_tum_* = tumor cell content (as calculated by ACEseq)

#### Structural variations detection and annotation

Somatic structural rearrangements (SVs) were detected using SOPHIA v.34.0 [https://bitbucket.org/utoprak/sophia/]. Briefly, SOPHIA uses supplementary alignments to identify SV candidates, which are then filtered by comparing them to a background control set. The background control set consists of 3261 normal blood samples from published TCGA and

ICGC studies and published and unpublished DKFZ studies, sequenced using Illumina HiSeq 2000, 2500 and HiSeq X platforms and aligned using the same workflow as in this study. Gencode v19 gene annotations were used to annotate the SV breakpoints and the closest gene up and downstream of the breakpoints. The COSMIC cancer census gene list was used to annotate the overlapping and closest cancer genes to each break point.

#### Copy number alterations

Allele-specific copy number alterations (CNAs) were detected using ACEseq v1.2.8-3 (65). ACEseq determines absolute allele-specific copy numbers as well as tumor ploidy and tumor cell content based on coverage ratios of organoid/tumor to its matched control, and the B-allele frequency (BAF) of heterozygous germline SNPs. SV calls were used to improve genome segmentation. When the control coverage was of insufficient quality due to lack of uniformity, another control sample with matching GC bias and gender was used. ACEseq also reported the genome chromosome instability scores for HRD, LST and TAI.

#### Mutational signatures

Mutational signature exposures were calculated using YAPSA development version 3.13 (66) using R 4.0.0. Briefly, the linear combination decomposition (LCD) of the mutational catalog with known and predefined COSMIC mutational signatures v2 (19) was computed by non-negative least squares (NNLS). The mutational signature analysis was applied to the mutational catalogs for SNVs for all tumor and organoid samples. Signature-specific cutoffs were applied and cohort level analysis was used for detecting signatures as recommended by Huebschmann et al. (66). The cutoff used correspond to “cost factors” of 6 for SNVs.

#### Integrative oncoprint and mutational recurrence analysis

SNVs, indels, SVs and CNAs were integrated into an oncoprint in order to account for all variant types in the recurrence analysis of the main organoid cohort. All genes with somatic SNVs or indels in coding regions (nonsynonymous, stop gain, stop loss, splicing, frameshift and non-frameshift events) and ncRNA (exonic) were included. Any SV directly lying on a gene (SV direct) were considered for oncoprints. SVs were also annotated to a gene when they were either within 100kb of a gene (SV near), or the gene was the closest gene (SV close) for SV recurrence analysis to account for regulatory mutations such as enhancer hijacking events. Any genes between the breakpoints of focal SVs (< 1 Mbp) were considered affected. To capture the precise target of focal CNAs we employed results from GISTIC2 (67).

#### Mutational significance analysis

The IntOGen pipeline (20) was used to identify significant cancer drivers in the main organoid cohort. IntOGen v 3.0.4 was installed via conda from the bbglab anaconda channel. The relevant conda environment setup included explicit definitions of python v3.5.5 (with libraries pandas v0.17). In addition, a local installation of perl v5.16.3 was used, with installation of perl libraries Digest-MD5 v2.58 via cpan and perl-DBI v1.627-4.el7.x86_64 via yum package managers. The background intogen database (bgdata) was automatically downloaded using the command ‘intogen --setup’ which downloaded the 20150729 background databases. The IntOGen run specific parameters included running on 4 cores, Matlab Compiler Runtime v8.1 (2013a) and MutSigCV v1.4. Significance thresholds of 10% FDR were used for oncodrivefm, oncodriveclust and mutsig. Sample thresholds of 2 and 5 were used for oncodrivefm and oncodriveclust respectively. X11 forwarding had to be enabled otherwise MutSigCV would throw and error.

### RNA sequencing

Total RNA was isolated directly from Matrigel-embedded organoids using the RNeasy Plus Mini Kit (Qiagen) using the modification of the standard protocol that allows the purification of total RNA containing small RNAs. Total RNA from snap-frozen tumor tissues was isolated using the AllPrep DNA/RNA/miRNA Universal Kit (Qiagen). Quality control was performed using Quant-iT RNA Kit, broad range (Thermo Fisher Scientific) and RNA ScreenTape on the TapeStation System (Agilent). RNA isolation and quality control was performed at the DKFZ-HIPO Sample Processing Laboratory. Library preparation and RNA sequencing (RNAseq) was performed at the DKFZ Genomics and Proteomics Core Facility. RNAseq libraries were prepared using the TruSeq Stranded mRNA Library Prep Kit (Illumina) and paired-end sequencing was performed using the Illumina HiSeq 4000 (2×100bp) or HiSeq X (2×150bp) platforms.

### RNA sequencing analysis

RNAseq reads were aligned and gene expression quantified as previously described (68). Briefly the RNAseq reads pairs were aligned to the STAR index generated reference genome (build 37, version hs37d5) using STAR v2.5.2b in 2 pass mode (69). Duplicate reads were marked using sambamba v0.4.6 and BAM files were coordinate sorted using SAMtools v1.19. featureCounts v1.5.1 (70) was used to perform reverse strand specific read counting for genes over exon features based on the Gencode v19 gene model. Read duplicates were not excluded.

When both read pairs had an alignment quality score of 255 (indicating a unique alignment) they were used towards gene reads counts. For total library abundance calculations, during TPM expression values estimation, genes on chromosomes X, Y, MT and rRNA and tRNA were omitted as they can introduce library size estimation biases.

### Transcriptional subtyping

RNAseq counts normalization and analysis were performed using DESeq2 (71) with “sample source” (organoid or tumor tissue) as the design after removing all genes with zero counts. Non-negative matrix factorization (NMF) analysis was performed using the NNLM library in R to decompose the variance-stabilizing transformed counts of the 50 Moffitt genes (8). The resulting signature matrix W corresponds to the “classical-like” and “basal-like” subtypes, while the H matrix determines the coefficients or weights of the signatures for each sample, which we defined as classical and basal-like scores. Convergence check of the NMF signatures was performed by running the NMF five times independently with different seeds. For categorical sample classification, a score threshold of 0.005 was applied; in cases where only one of the scores was higher than the threshold, this defined the sample subtype, while in cases where both classical and basal-like scores were higher than the threshold, the subtype was defined as “intermediate”.

The PCA plot was generated based on the top 1000 most variable genes. Heatmap plots were generated using the R package ComplexHeatmap (72).

### Independent component analysis

Independent component analysis (ICA) was performed on the normalized RNAseq count matrix of PDO samples to deconvolute the organoid transcriptome into 10 reproducible, robust, and biologically relevant components using the fastICA v1.2 implemented in Bioconductor-mineICA v1.26.0 (73). Particularly, clusterFastICARuns() was used setting the number of iterations (nbIt) as 500 and alg.type= “parallel”, i.e., the independent components (IC) were extracted simultaneously. The ICA decomposition of the expression matrix X = AS yielded A, the mixing matrix (consisted of the activity of the ICs on the PDO samples) and S, the source matrix (provided the contribution of each gene to the ICs). Hence, the source matrix (S) was used in the biological interpretation of the IC(s) by ranking the genes in descending order of their contributing value. Spearman correlation coefficient test was performed to check the IC correlations to the Moffitt basal-like/classical scores as well as the drug response AUCd/AUCpi parameters using Pingouin-python package.

### Gene set enrichment analysis of independent components

Gene set enrichment analysis (GSEA) was performed using the R package clusterProfiler (74). For each IC, genes were ranked according to their contribution and tested for enrichment in custom gene sets. P values were adjusted using the *fdr* method. The threshold for significant over-representation was set at an adjusted *P* < 0.05.

### Differential expression and enrichment analyses of PDO subtypes

Differential expression testing was performed using the R/Bioconductor package DESeq2 (71). Genes without at least one count in three organoid samples were filtered out before differential expression analysis. An adjusted *P* < 0.1 was considered significant. Functional enrichment analysis for ENCODE and ChEA Consensus TFs from ChIP-X and MSigDB v7.2 Hallmark collections was applied to the significant differentially expressed genes using EnrichR (75). The threshold for significant enrichment was set at an adjusted *P* < 0.01 and significant gene sets were sorted by EnrichR “Combined Score”.

### Survival analysis

Patients with recurrent disease or that received neoadjuvant therapy were excluded from survival analyses. Death occurring within 90 days after surgery was considered peri-operative and therefore also excluded from survival analyses. Overall survival (OS) was defined as the time from surgery to death or last follow-up. Median survival was estimated using the Kaplan– Meier method, patients still alive at the last follow-up were censored. The log-rank test was used to compared survival curves between groups. Differences were considered statistically significant if *P* < 0.05. Survival analysis and plots were performed using the R packages survival and survminer.

### Drug screening and drug response analysis

Drug screen was performed using the DeathPro workflow (28) on a subset of 39 PDAC-PDOs to assess the response to the chemotherapeutic drugs gemcitabine, 5-fluorouracil (5-FU), irinotecan, oxaliplatin, paclitaxel and erlotinib. Drug concentrations, treatment intervals and endpoints were chosen according to published studies or determined in pilot experiments (**Supplementary Table S4A**). All image data was used and analyzed. To assess reproducibility, some drug tests were performed in two independent biological replicates. Organoids were seeded at different time points and treated with drugs using different plate layouts. Biological variability in all tested conditions was assessed by imaging two positions per well and no other technical replicates were included. For drug testing 1000-2500 organoid forming units were mixed with 10 µl growth-factor reduced, phenol red-free Matrigel (Corning, >9 mg/ml protein) per well and seeded in 96-well Angiogenesis µ-Plates (ibidi). Organoids were grown for three days in growth medium, then stained with 1 µg/ml Hoechst (Invitrogen) and 1 µg/ml propidium iodide (Sigma), incubated for 6 h and imaged (0 h timepoint). After imaging, the dye-containing medium was substituted with drug-containing medium. After 72 h of drug treatment, organoids were stained with Hoechst and propidium iodide, incubated for 6 h and imaged again (72 h timepoint). After imaging, the dye-containing medium was substituted with growth medium. Organoids were grown for additional 72 h and then stained with Hoechst, propidium iodide and 500 nM SiR-Actin (Cytoskeleton, Inc) for 6 h and imaged again (144 h timepoint). Cells were exposed maximally to 1% DMSO in the highest drug concentrations and corresponding controls were included in the assay. Organoids were imaged at similar positions at 0 h, 72 h and 144 h after start of drug treatment using a Zeiss LSM780 confocal microscope, 10x objective (EC Plan-Neofluar 10x/0.30 M27) and 405 nm, 561 nm and 633 nm diode lasers in simultaneous mode. Imaging was performed in an incubation chamber at 37 °C, 5% CO2 and 50-60% humidity using the Visual Basic for Applications macro ‘AutofocusScreen’ (76). Image processing and drug response analysis was performed with DeathPro workflow as described and published in Jabs et al., 2017 (28). For segmentation, local threshold parameters were adjusted (Hoechst: rad=5, c=-5; PI: rad=100, c=-20) and objects were filtered according to size (7 pixel < object < 40,000 pixel). LD50 (median lethal dose) values were filtered to retain only robust fits: LD50 values smaller or larger than the minimum or maximum drug concentration, respectively, and LD50 values with confidence intervals larger than a factor of 1000 were removed.

### Correlation analysis of drug response with gene expression

Normalized RNAseq counts from 44 PDTOs were filtered for protein-coding genes. After inspecting the distribution of counts and not available values, further filtering was performed for the correlation analysis: only genes that were detected in at least 41 of 44 organoid samples were included, to avoid comparing correlations stemming from different sample sizes. In addition, a bimodal distribution of RNAseq counts was observed and the analysis was focused to medium to highly expressed genes (log_2_counts > 7) to ease potential follow-up analyses. No further filtering was performed and genes expressed with high and low variation across organoids were included. In total, 12,236 genes were used for calculating spearman correlation estimates and *P* values at a confidence level of 0.95 in a two-tailed test. *P* value distributions were checked for unexpected behavior and to compare association of response and gene expression between drugs. Genes sets containing genes correlating positively or negatively (*P* < 0.01) with each response parameter (AUCd_144h_ or AUCpi_144h_) were generated for each drug. For downstream analyses, gene sets derived for PDOs from chemo-naïve patients were included.

Gene overlaps for the different gene sets were depicted using the UpSet plot implementation within the Complexheatmap package (72), setting the mode to *intersection*.

### Over-representation analysis

Over-representation analysis for Wikipathways and KEGG pathways was conducted for drug response genes sets using the clusterProfiler package (74). *P* values were adjusted using the *fdr* method. The threshold for significant over-representation was set at an adjusted *P* < 0.05.

For over-representation analysis of IC genes was conducted for the top 100 or bottom 100 genes according to their loading values using the same parameters described above.

### Targeted drug testing

For testing of targeted agents, eight organoid lines were selected based on their molecular status. The drug tested included palbociclib (Selleckchem), olaparib (Selleckchem), and the fungal toxin α-amanitin (Sigma-Aldrich). Organoids were dissociated into single cells and small aggregates by enzymatic dissociation with TrypLE. Cells were resuspended in Matrigel at a density of 2000 cells/10 µL, seeded into the inner chamber of µ-Chamber Angiogenesis 96-well-plates (ibidi) and cultured with 70 µl organoid medium. The drug or vehicle treatment was applied three days after seeding. Dilution rows of the drugs (1:3, α-amanitin 1:2) were freshly prepared before application from frozen stocks. After 72 hours of treatment, half of the medium was replaced by fresh medium. Drug-induced cell death and proliferation inhibition were quantitively assessed using the DeathPro workflow (28) 8 days after seeding (120 h after treatment). At days 3 and 8, organoids were stained with 1 μg/ml Hoechst (Invitrogen) and 1 μg/ml propidium iodide (Sigma) in organoid medium. Automated confocal imaging was performed according to the DeathPro assay image acquisition procedures described above. Images of two positions per well were acquired and each organoid line was screened once. Image stacks were processed to Maximum Intensity Projections (MIPs) and dose-response-curves were generated from the MIPs. Dose-response curves were manually checked for plausibility by comparison with MIPs and severe outliers were excluded. Given that we observed a column-wise growth bias most likely due to a seeding effect, we used the lowest drug concentration conditions, which were located in the same column as the treatment conditions, as individual baselines (controls) for each drug. Area under the curve values were calculated for cell death (AUCd) and proliferation inhibition (AUCpi) from the corresponding dose-response curves. LD50 values were derived as described above. Wilcoxon signed-rank test was used for comparisons between groups.

### Mutual exclusivity analysis

Mutual exclusivity analysis was performed on all genes with a minimum recurrence threshold of 5. We applied the Fisher’s exact test and the CoMET test (77). Fisher’s right tailed test was used to support co-occurrence when the number of samples with alterations in both genes is significantly higher than expected by chance. The Fisher’s left tailed suggests mutual exclusivity when the number of samples with alterations in both genes is significantly lower. A *P* value of < 0.01 was used as cut-off for mutual exclusivity and inclusivity.

### Data availability

All raw sequencing data has been deposited at the European Genome-Phenome Archive under the accession number EGAS00001005474. All somatic mutation calls and integrated mutations tables on which the analysis was performed will be available upon peer-reviewed publication in Zenodo (10.5281/zenodo.5208420).

## Acknowledgments

This work received financial support from NCT 3.0 Extension Programm (NCT3.0_2015.17 PrecO-Panc). We thank the NCT Molecular Precision Oncology Program for technical support and funding through HIPO-015 K20K. S.L.B. and O.S. were supported by the National Center of Tumor Diseases (NCT) Heidelberg (NCT3.0_2015.17 PrecO-Panc). Human specimens and clinical data were obtained from the EPZ-Pancobank (Biobank of the European Pancreas Center at the Department of General Surgery, University Hospital Heidelberg) in accordance with the regulations of the tissue biobanks and an approval by the Ethics Committee of Heidelberg University (ethic votes 301/2001, 159/2002, S-206/2011, S-708/2019). Activities of the EPZ-Pancobank were supported by the Heidelberger Stiftung Chirurgie and in part by the German Ministry of Science and Education (BMBF) grants 01ZX1305C, 01ZX1605C, 01KT1506. The EPZ-Pancobank is a member of the BioMaterial Bank Heidelberg/BMBH (coordinator: Prof. P. Schirmacher, BMBF grant#01EY1701) belonging to the German Biobank Alliance. We thank the DKFZ-HIPO Sample Processing Laboratory and DKFZ Genomics and Proteomics Core Facility for excellent support in sample processing and sequencing. We thank ODCF for sequencing data management. We thank K. Felix for helpful discussions. We thank K. Schneider, E. Lederer, K. Ruf and M. Fischer for the excellent technical support.

## Author’s contributions

S.L.B: experimental design, organoid establishment, sample generation, drug screening experiments and targeted testing, data analysis, integration, interpretation and visualization, writing original manuscript. N.I: genomic and transcriptomic data analysis, mutational signatures, mutual exclusivity analysis, data integration, interpretation and visualization, manuscript edition. J.J: drug screening experiments and data analysis, drug-transcriptomic correlation analysis, data integration, interpretation and visualization, manuscript edition. T.B: genomic and transcriptomic data analysis, mutational signatures. S.S: targeted drug testing and data analysis. Q.H: Drug screening and targeted drug testing. O.D: independent component analysis. F.W.T: RNAseq data normalization and NMF. C.S.L: patient follow-up, clinical data, scientific support, manuscript edition. A.K.K: tissue sample acquisition, clinical data. M.B: genomic data analysis. C.E: pathological analyses. M.M.G: pathological analyses. M.V: Sample analysis, scientific support. D.H: mutational signatures. M.S: tissue sample acquisition and clinical data. R.O: funding acquisition, scientific and infrastructural support. N.A.G: tissue sample acquisition and clinical data, scientific and infrastructural support. M.S: funding acquisition, scientific and infrastructural support, genomic data analysis. M.W.B: patient inclusion, funding acquisition, scientific and infrastructural support. R.E: project conception, design and supervision, funding acquisition, scientific and infrastructural support. C.C: project conception, design and supervision, funding acquisition, data interpretation. O.S: project conception, design and supervision, patient inclusion, funding acquisition, data interpretation, manuscript writing.

All authors reviewed and approved the final manuscript.

---

**Supplementary Figure S1.**
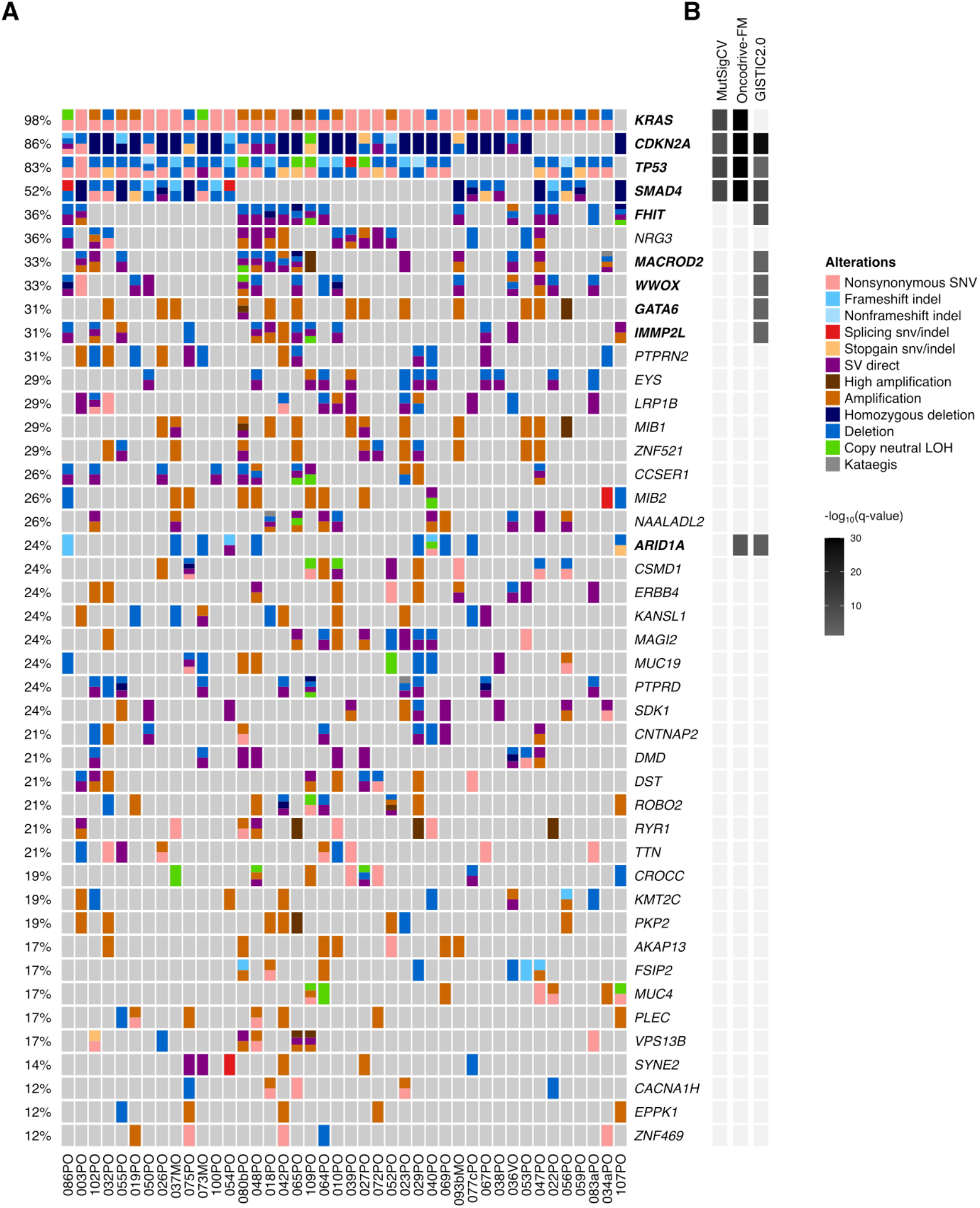
Mutational landscape of the main organoid cohort. **A,** Oncoprint for the most recurrent altered genes (>12% frequency). **B,** Significance of the alterations according to MutSigCV, Oncodrive-FM and GISTIC2.0.

**Supplementary Figure S2.**
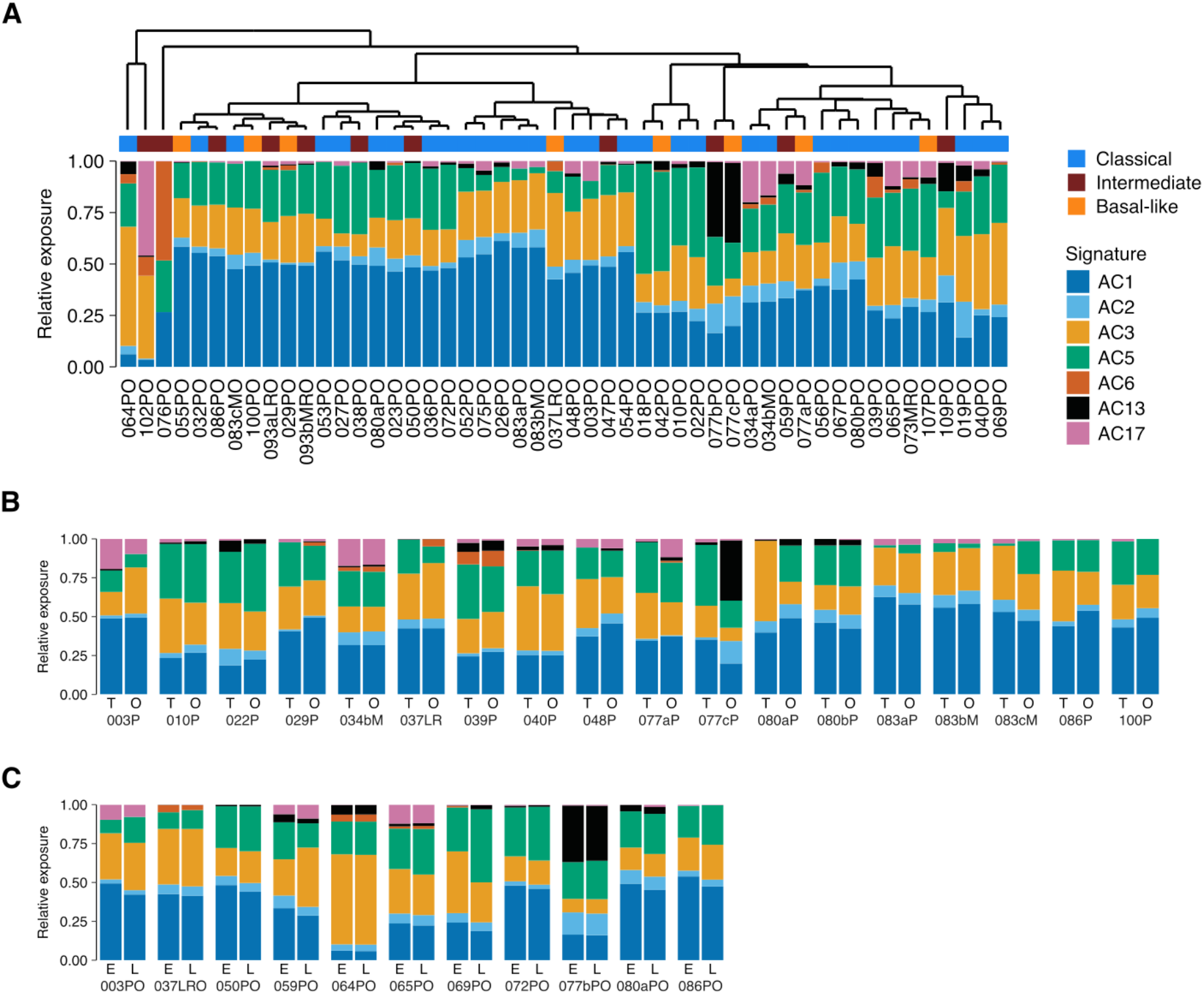
Mutational signatures profiles are preserved in organoid cultures. **A,** Relative mutational signature exposures identified in the PDAC-PDO catalogue. **B,** Relative mutational signature exposures in tumors (T) and matched organoids (O). **C,** Relative mutational signature exposures in organoids at early (E) and later (L) passages. AC1: clock-like, AC2: APOBEC, AC3: HRR deficient, AC5: clock-like, AC6: MMR deficient, AC13: APOBEC, AC17: unknown.

**Supplementary Figure S3.**
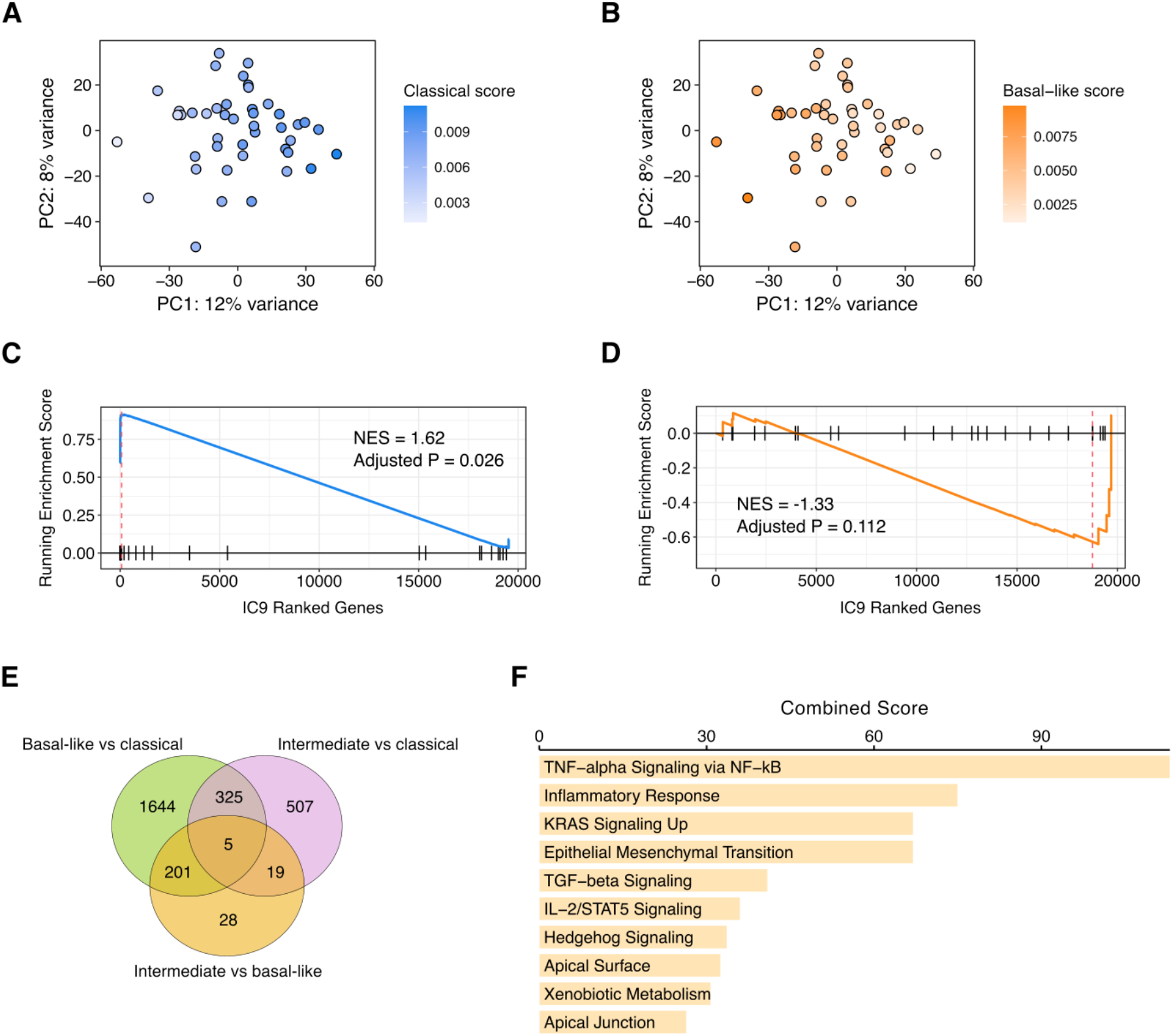
PDAC transcriptional subtypes in organoids. **A-B,** Principal component (PC) analysis of the 1000 most variant genes expressed by organoids colored according to classical and basal-like scores. **C-D,** GSEA for Moffitt’s classical (C) and basal-like (D) signature genes in the independent component IC9. Genes are ranked according to their contribution. NES: normalized enrichment score. **E,** Venn diagram indicating the overlap of differentially expressed genes (adjusted *P* < 0.1) in organoids for the indicated subtype comparisons. **F,** Top 10 enriched Hallmark gene sets (adjusted *P* < 0.01) in basal-like organoids versus classical organoids.

**Supplementary Figure S4.**
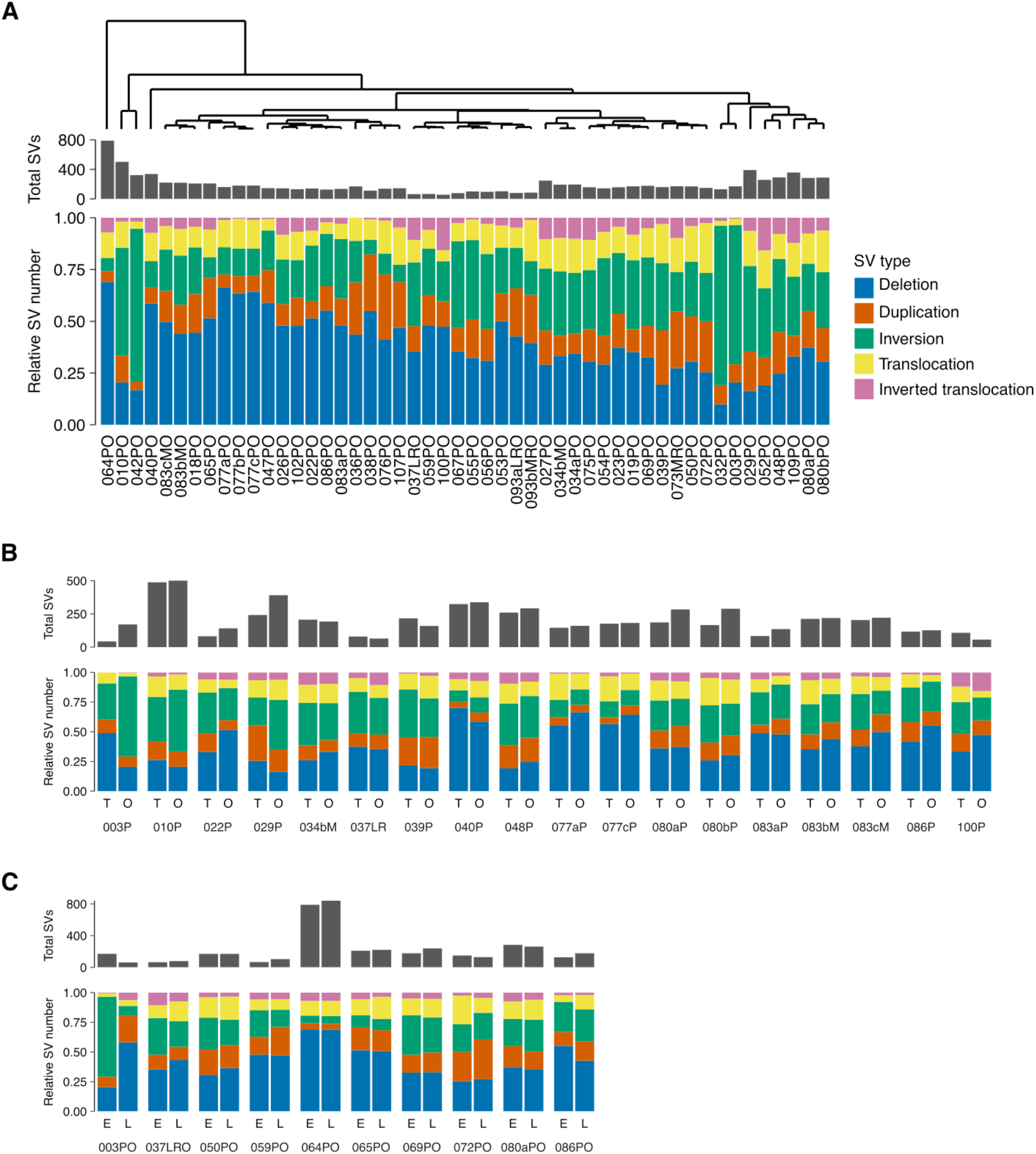
Structural variant (SV) profiles of organoids and parental tumors. **A,** Total and relative number of structural variants detected in the PDAC-PDO catalogue. **B,** Structural variants in tumors (T) and matched organoids (O). **C,** Structural variants in organoids at early (E) and later (L) passages.

**Supplementary Figure S5.**
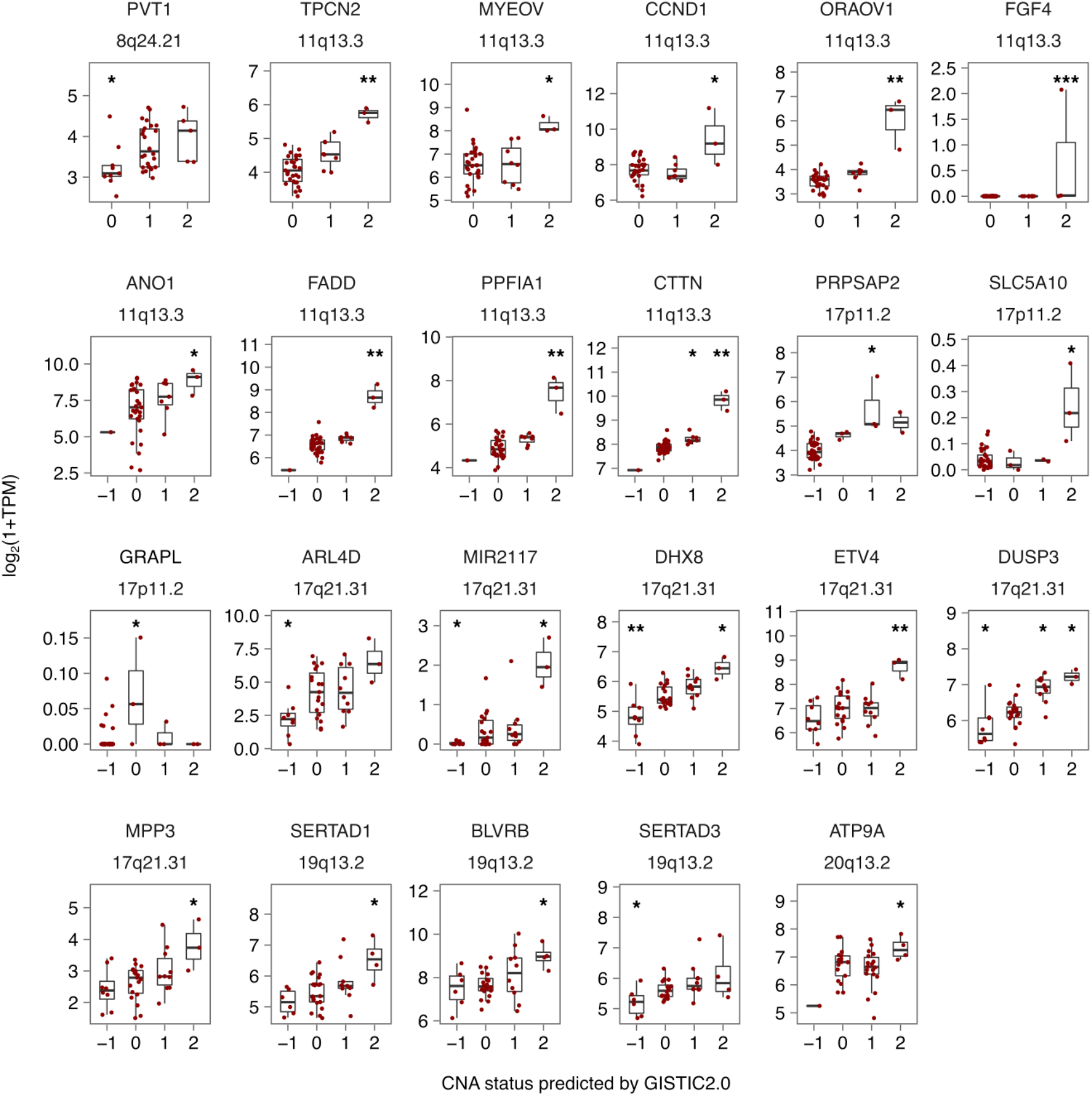
Expression levels of genes in significantly amplified regions categorized by their CNA status as predicted by GISTIC2.0. Only genes with a significant difference in expression in any of the CNA groups are displayed (Wilcoxon test, comparisons against base-mean as reference group, *P* < 0.05). TPM: transcript per million. * *P* < 0.05, ** *P* < 0.01, *** *P* < 0.001.

**Supplementary Figure S6.**
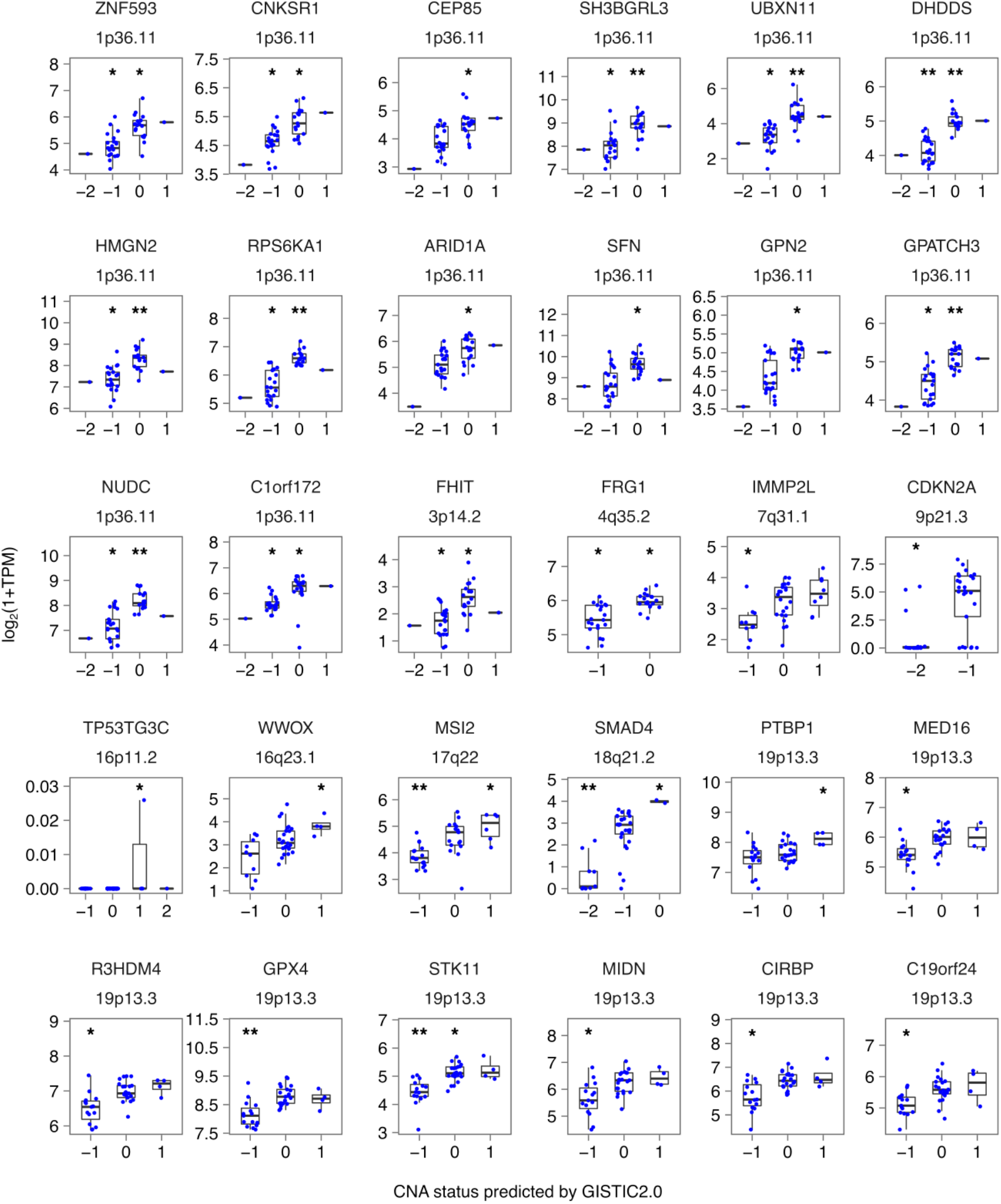
Expression levels of genes in significantly deleted regions categorized by the CNA status as predicted by GISTIC2.0. Only genes with a significant difference in expression in any of the CNA groups are displayed (Wilcoxon test, comparisons against base-mean as reference group, *P* < 0.05). TPM: transcript per million. * *P* < 0.05, ** *P* < 0.01, *** *P* < 0.001.

**Supplementary Figure S7.**
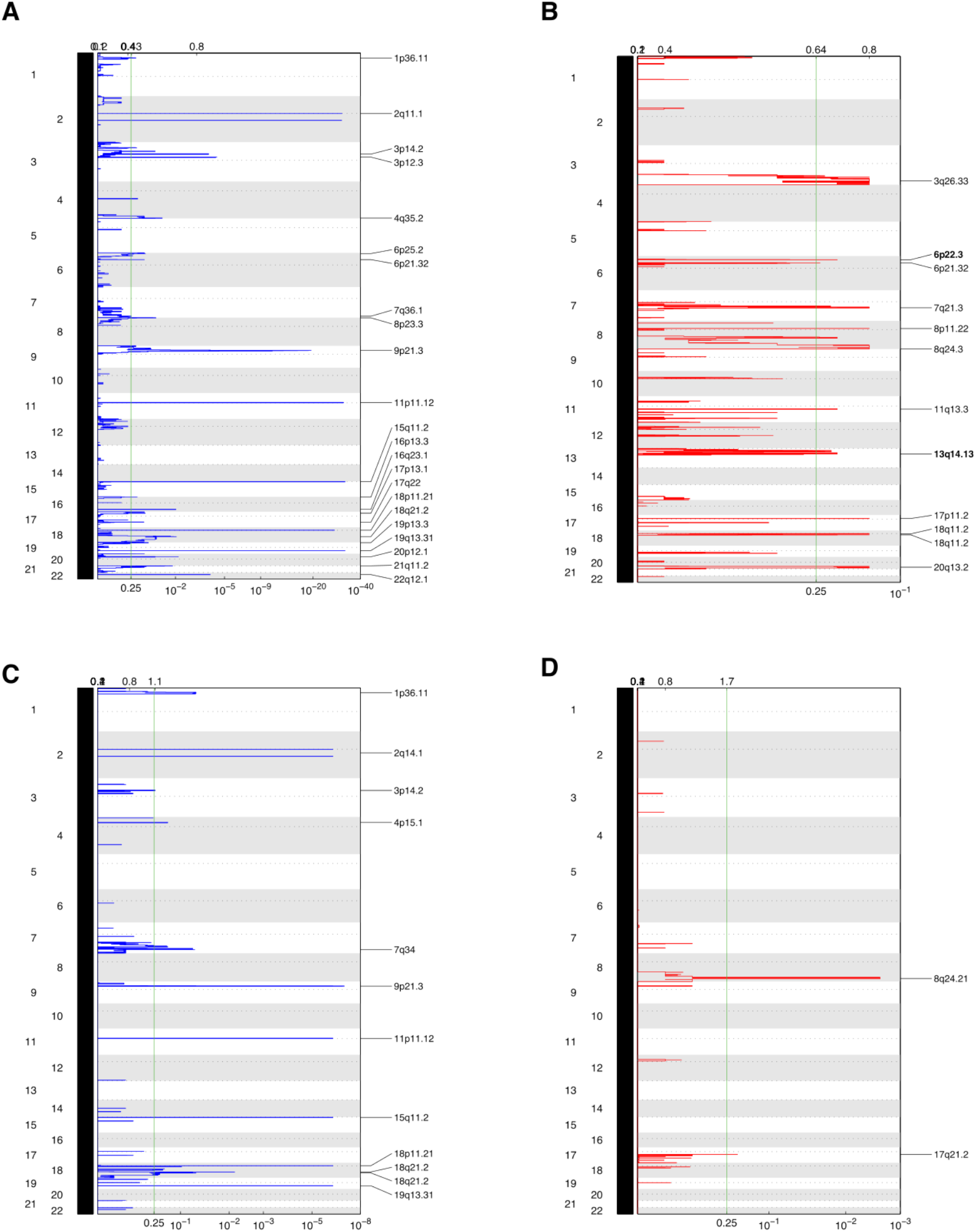
Significant CNAs in classical and basal-like organoids predicted by GISTIC2.0. **A,** Deletion and **B,** amplification peaks in classical organoids. **C,** Deletion and **D,** amplification peaks in basal-like organoids.

**Supplementary Figure S8.**
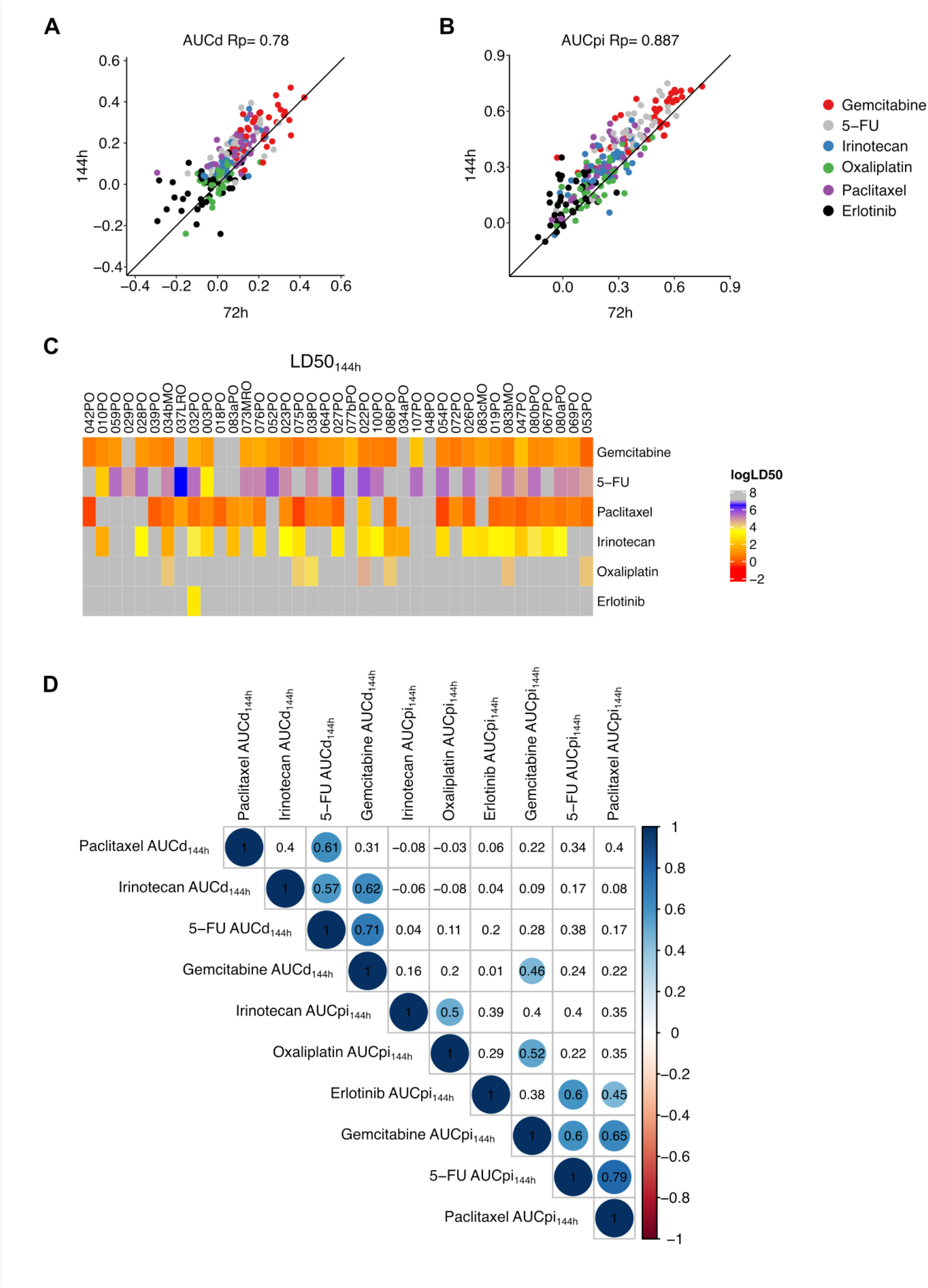
DeathPro drug screening of PDOs. Correlation of **A,** AUCd and **B,** AUCpi between DeathPro assay readouts at 72 h and 144 h. Rp: Pearson’s correlation coefficient. **C,** LD50 values derived from death dose-response curves for the 144h time-point. **D,** Correlation matrix for drug responses of the 39 organoid lines included in the drug screen. Spearman’s correlation coefficients are displayed. Significant correlations are denoted by a colored circle according to the Spearman’s correlation coefficient.

**Supplementary Figure S9.**
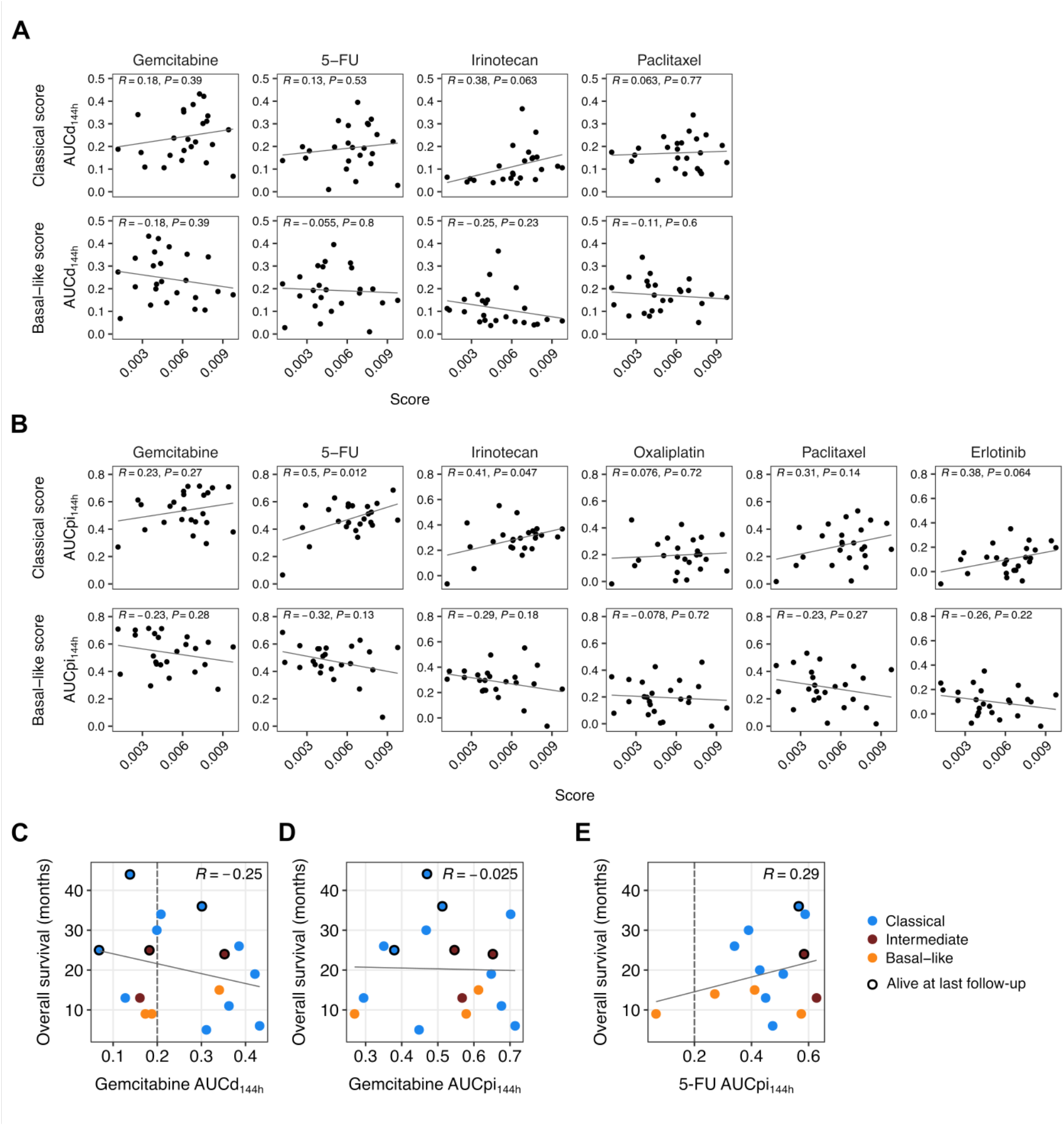
Association of PDO drug response with subtype scores and patient survival. Correlation of **A,** AUCd_144h_ and **B,** AUCpi_144h_ with classical and basal-like scores. Only organoid lines derived from chemo-naïve patients at resection were included in this analysis. Correlation of organoid drug responses for **C,** gemcitabine AUCd_144h_, **D,** gemcitabine AUCpi_144h_ and **E,** 5-FU AUCpi_144h_ with overall survival for patients receiving adjuvant therapy containing the corresponding drug. Only organoid lines derived from chemo-naïve patients at resection were included in this analysis. Dotted lines denote the response threshold of 0.2 AUC units.

**Supplementary Figure S10.**
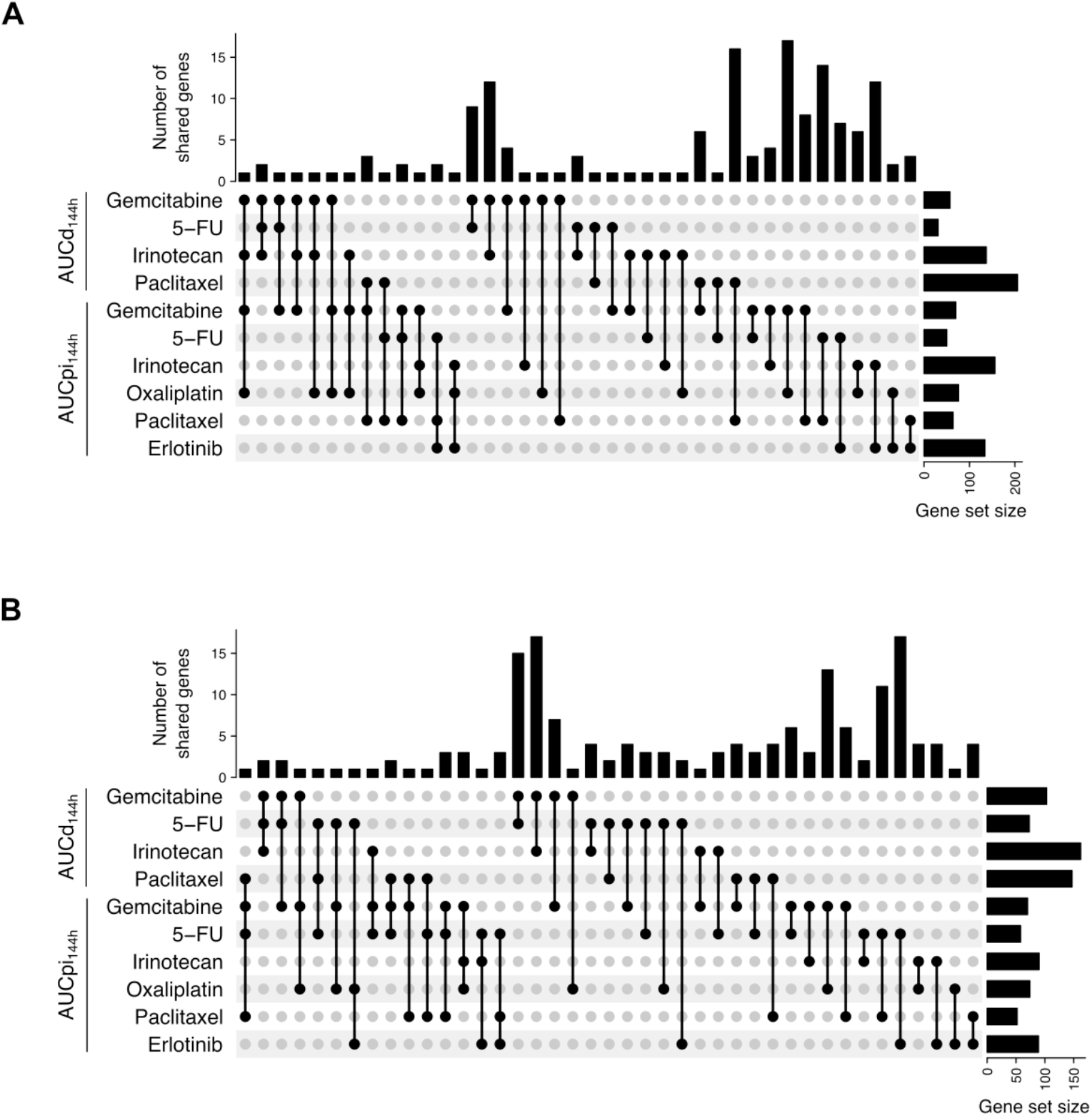
UpSet plots for the overlap (intersection) of genes among gene-drug response correlation gene sets. **A,** Number of genes shared by the indicated up-in-resistant gene sets (expression negatively correlated with AUCd_144h_ or AUCpi_144h_). **B,** Number of genes shared by the indicated down-in-resistant gene sets (expression positively correlated with AUCd_144h_ or AUCpi_144h_).

**Supplementary Figure S11.**
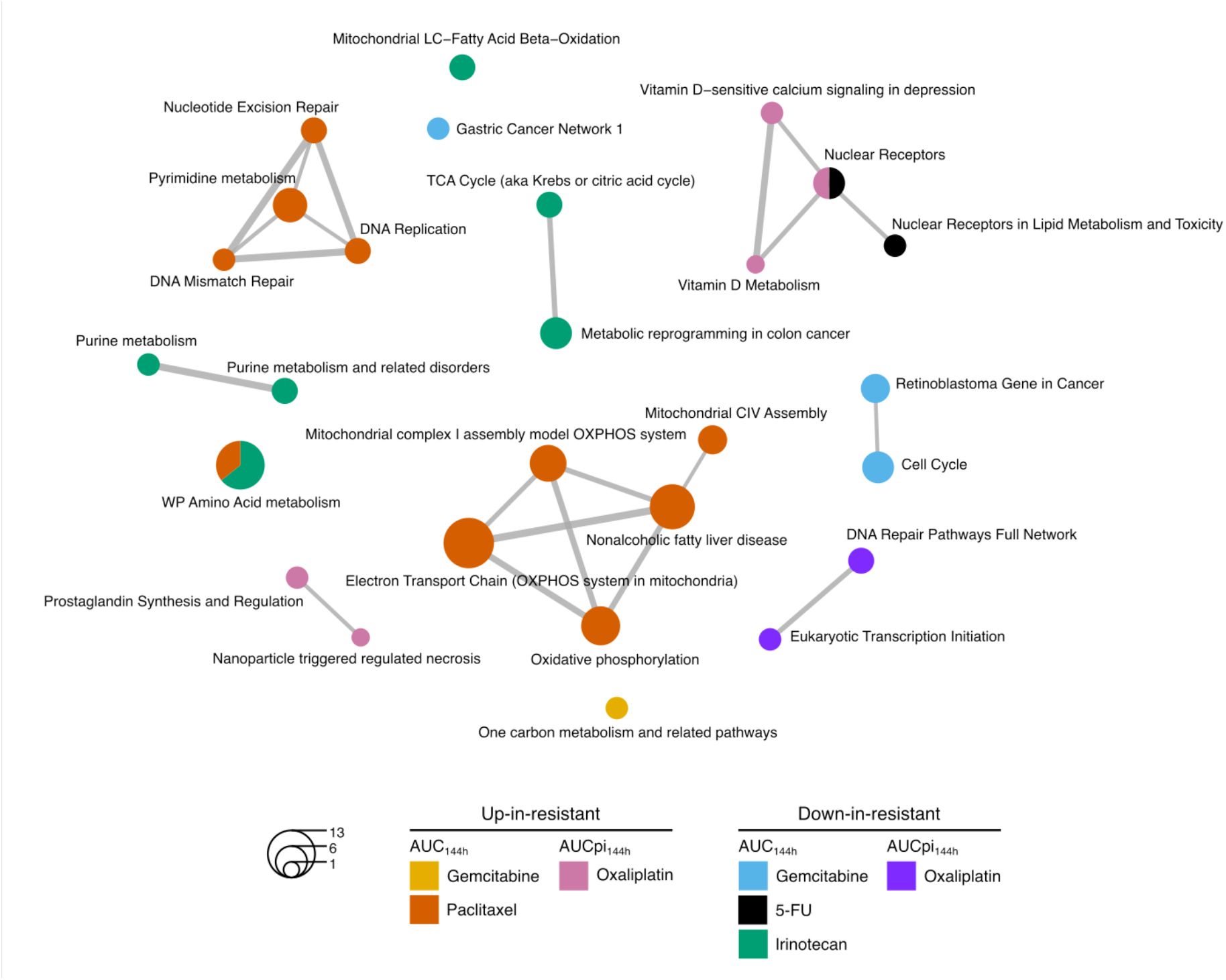
Enrichment map for significantly enriched Wikipathways terms (nodes; adjusted *P* < 0.05) for each gene-drug response correlation gene set as shown in Fig. 5B, depicting overlapping gene sets. Each color represents an individual gene set. Nodes sizes represent the number of genes and the edge thickness the degree of overlap between two terms.

**Supplementary Figure S12.**
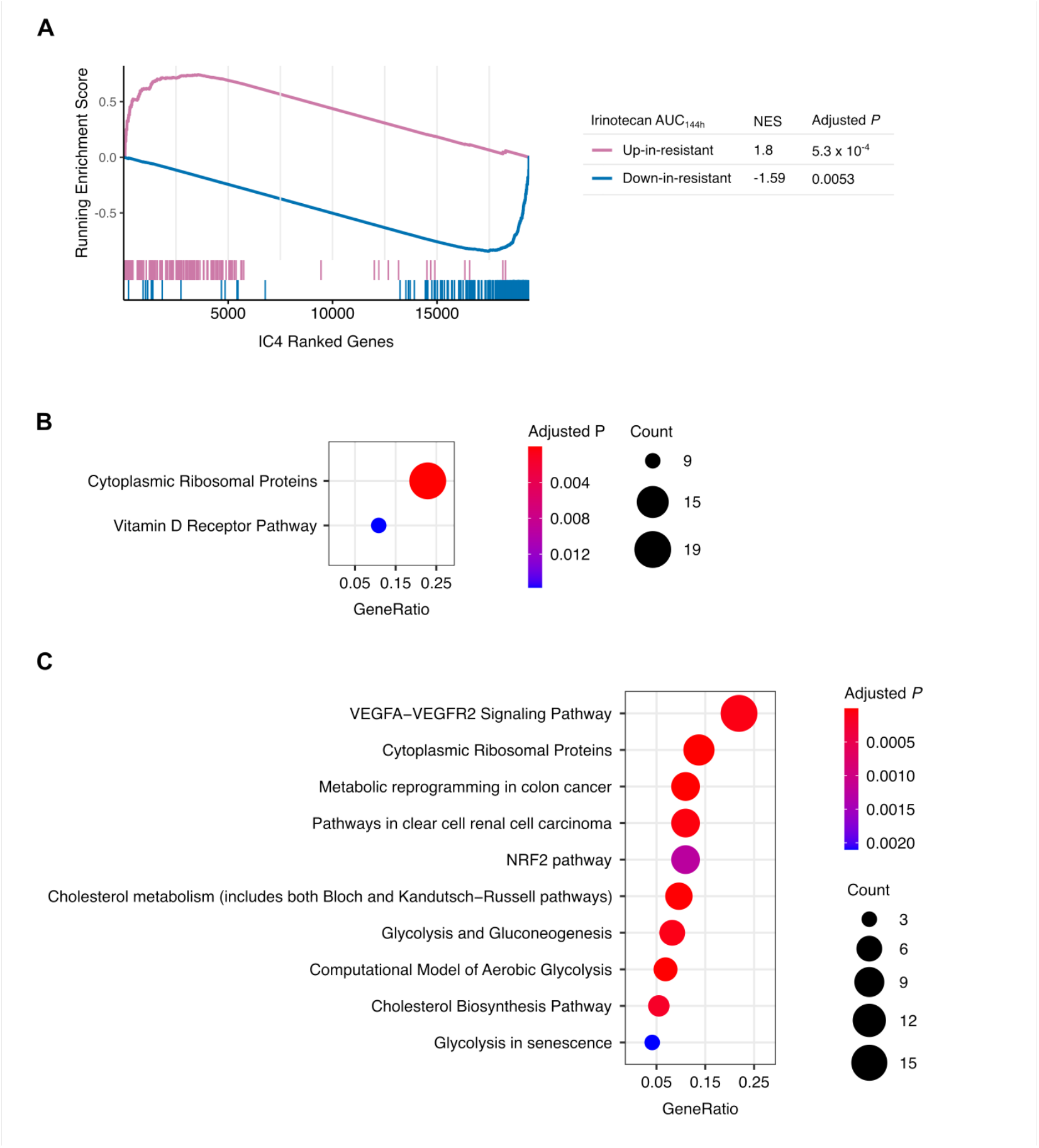
Association of irinotecan AUCd_144h_ gene sets and IC4. **A,** Gene set enrichment analysis of irinotecan AUCd_144h_ up-in-resistant and down-in-resistant gene sets in IC4. Genes are ranked according to their contribution. NES: normalized enrichment score. **B,** Over-represented Wikipathways (adjusted *P* < 0.05) for the top 100 IC4 genes. **C,** Top 10 over-represented Wikipathways (adjusted *P* < 0.05) for the bottom 100 IC4 genes.

**Supplementary Figure S13.**
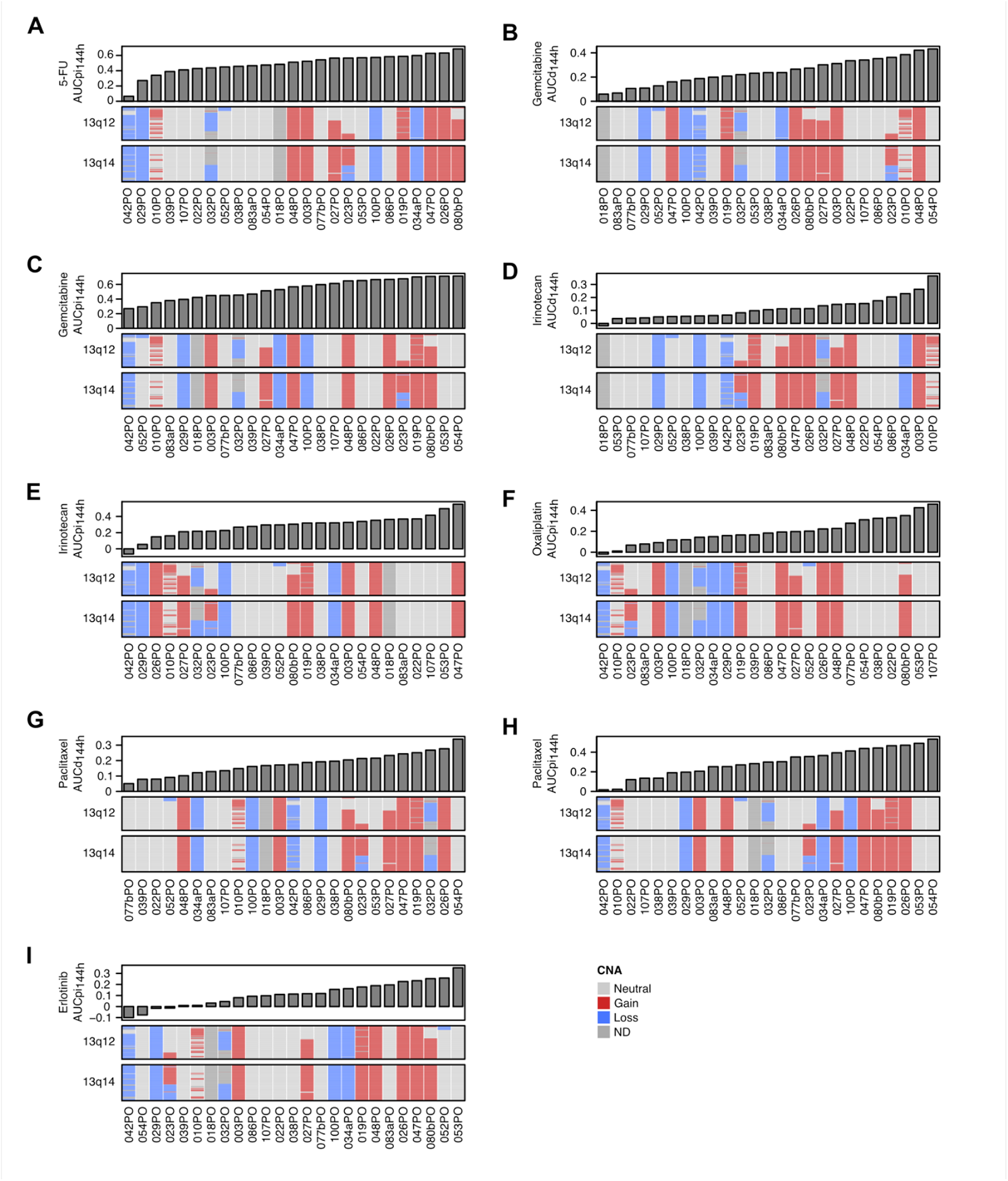
Copy number alterations in chr13q12/chr13q14 and PDO drug responses. **A,** 5-FU AUCpi_144h_. **B,** Gemcitabine AUCd_144h_. **C,** Gemcitabine AUCpi_144h_. **D,** Irinotecan AUCd_144h_. **E,** Irinotecan AUCpi_144h_. **F,** Oxaliplatin AUCpi_144h_. **G,** Paclitaxel AUCd_144h_. **H,** Paclitaxel AUCpi_144h_. **F,** Erlotinib AUCpi_144h_. Samples are sorted by drug response. Genes are sorted by their position along the cytobands.

**Supplementary Figure S14.**
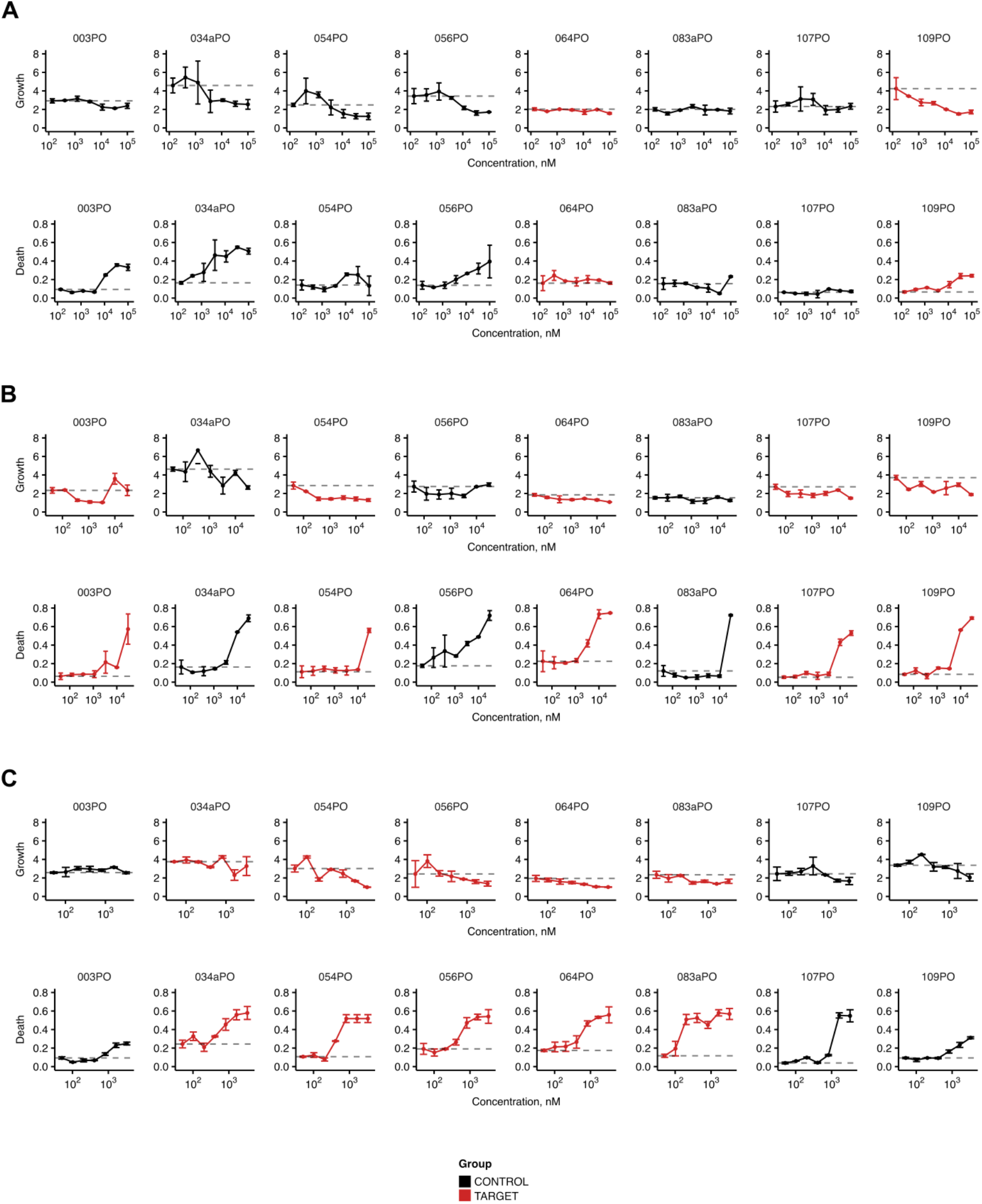
Individual dose-response curves for growth and death for targeted therapies. **A,** Olaparib. **B,** Palbociclib. **C,** α-amanitin.

