## Supplementary Data for "Deep molecular characterization linked to drug response profiling of pancreatic ductal adenocarcinoma using patient-derived organoids"

**Supplementary data: Genomic representability and heterogeneity of PDAC-PDOs**

Related to Figure 2.

Plots on the left: Correlations of predicted allele counts for non-silent SNVs and indels between pairs of related primary tumor/metastatic tissue and organoid samples.

Plots on the rights: Predicted allele count displayed according to their chromosomal location. Purple dots represent SNV/indels with higher allele count in the sample of the upper panel (or x-axis on the left plot) and yellow dots represent SNV/indels with higher allele count in the sample of the lower panel (or y-axis on the left plot). Gray dots represent SNV/indels with equivalent allele count in both samples.

### Patient 003

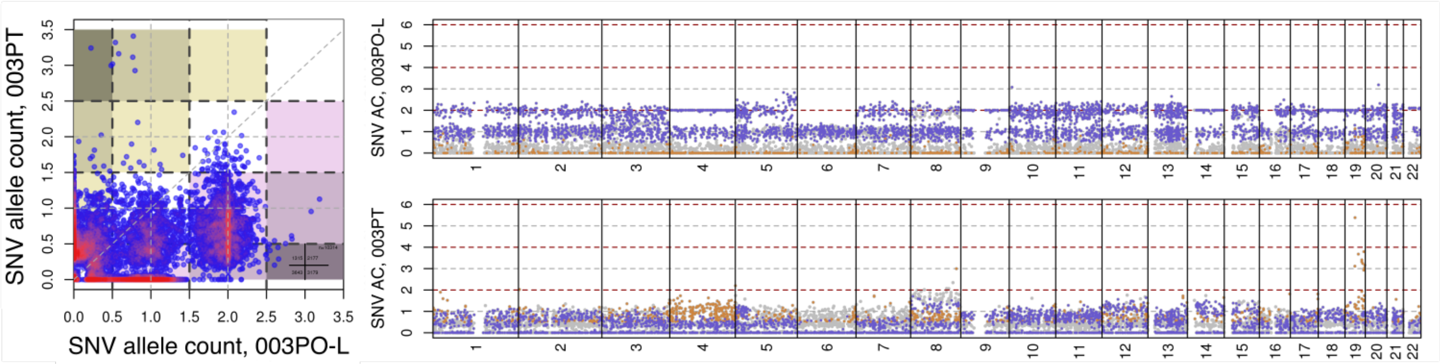


### Patient 010

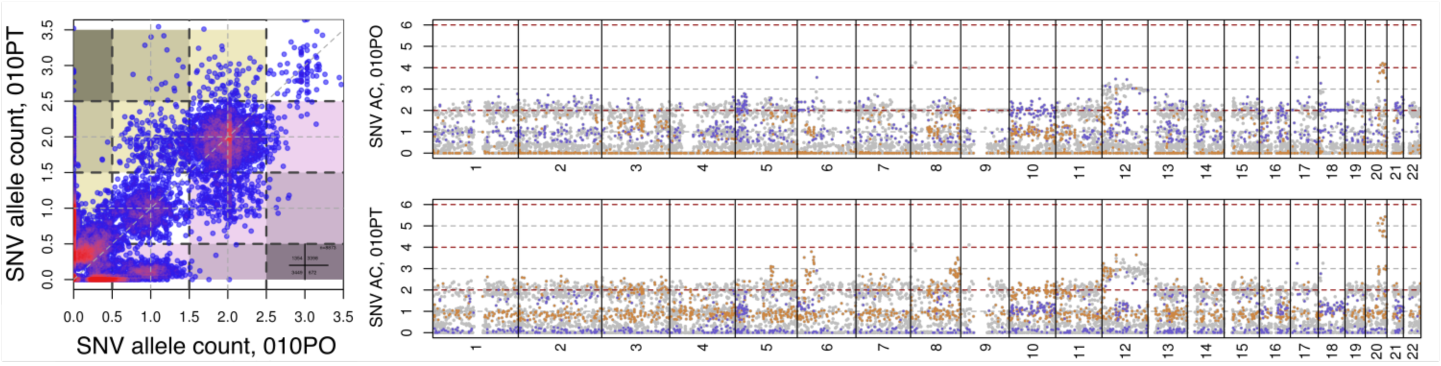


### Patient 022

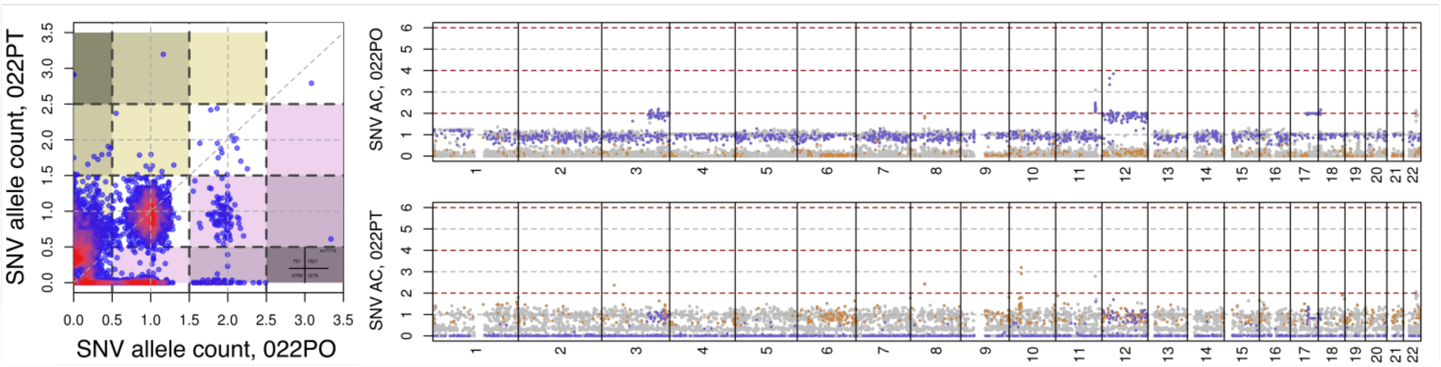


### Patient 029

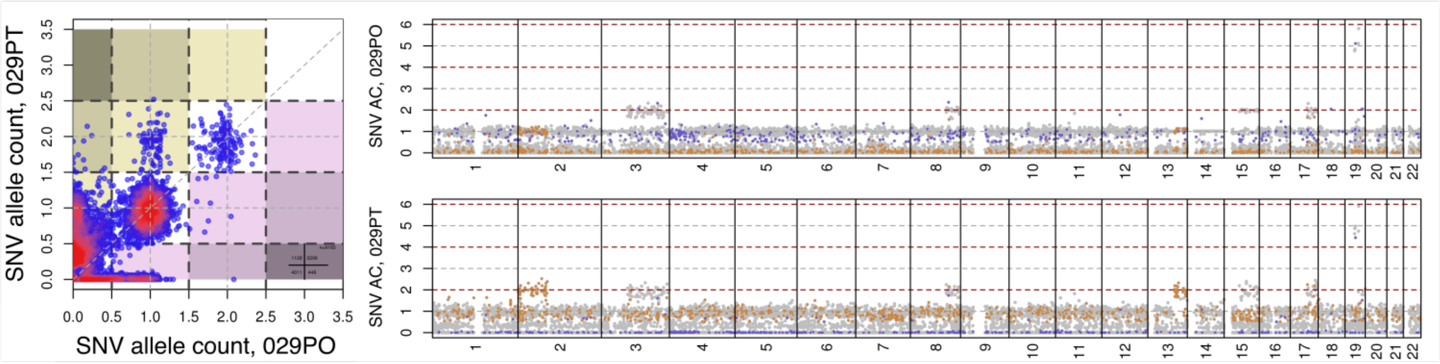

### Patient 034

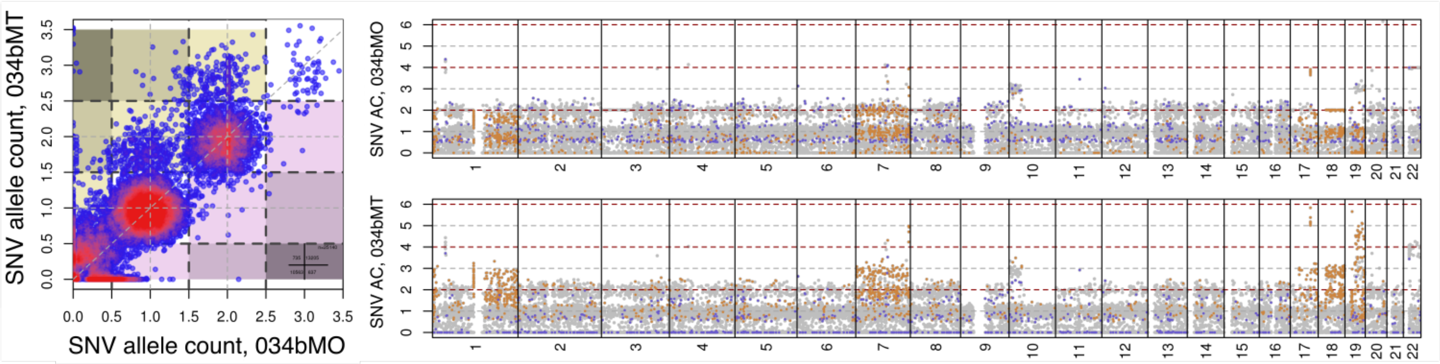


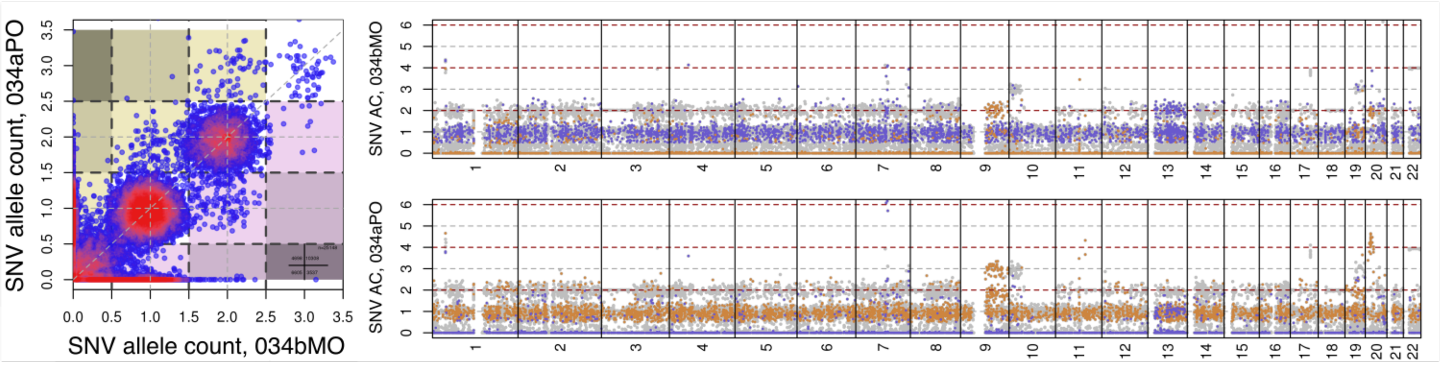


### Patient 037

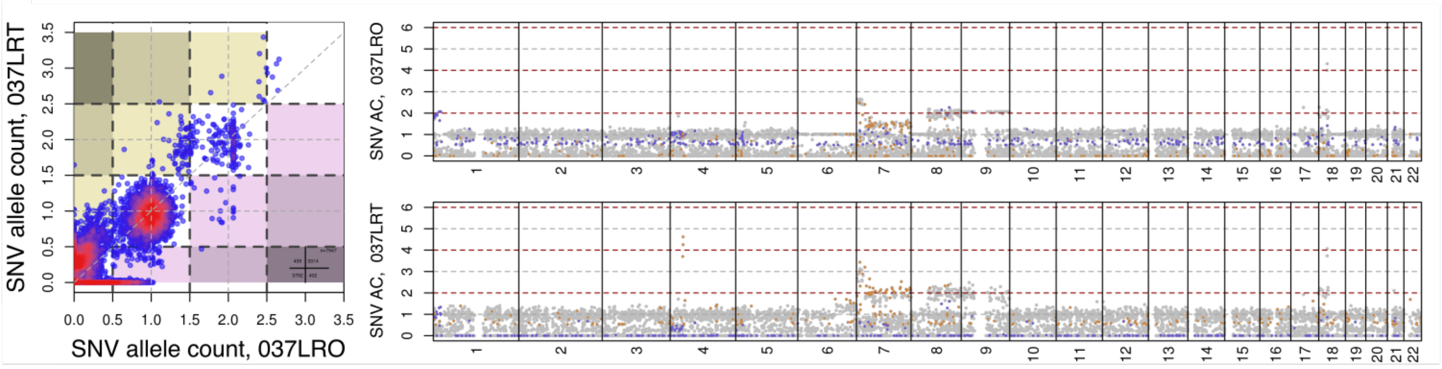


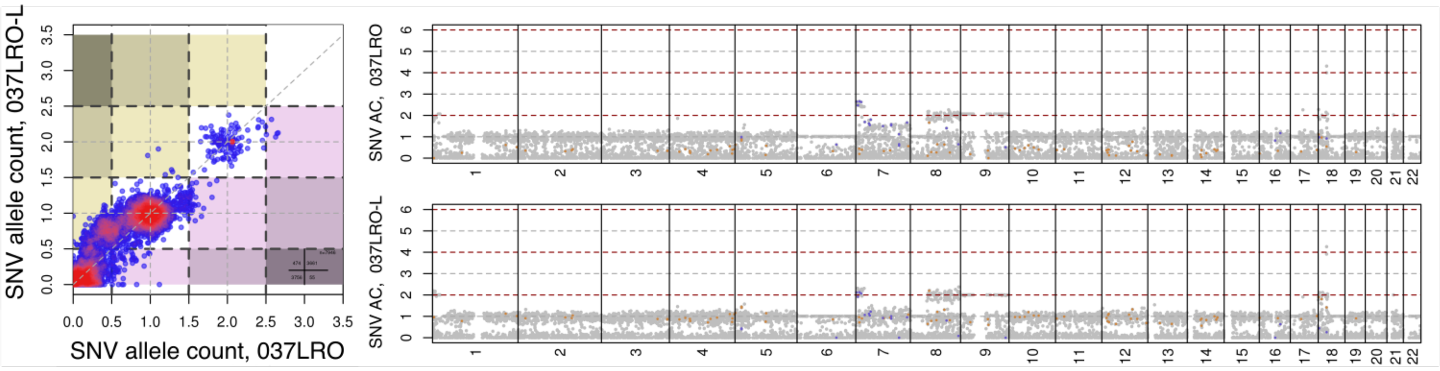

### Patient 039

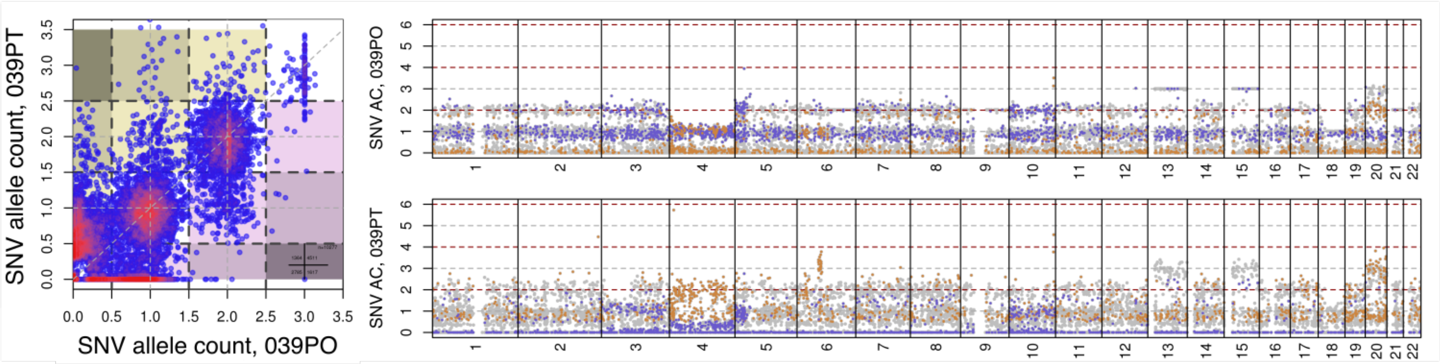


### Patient 040

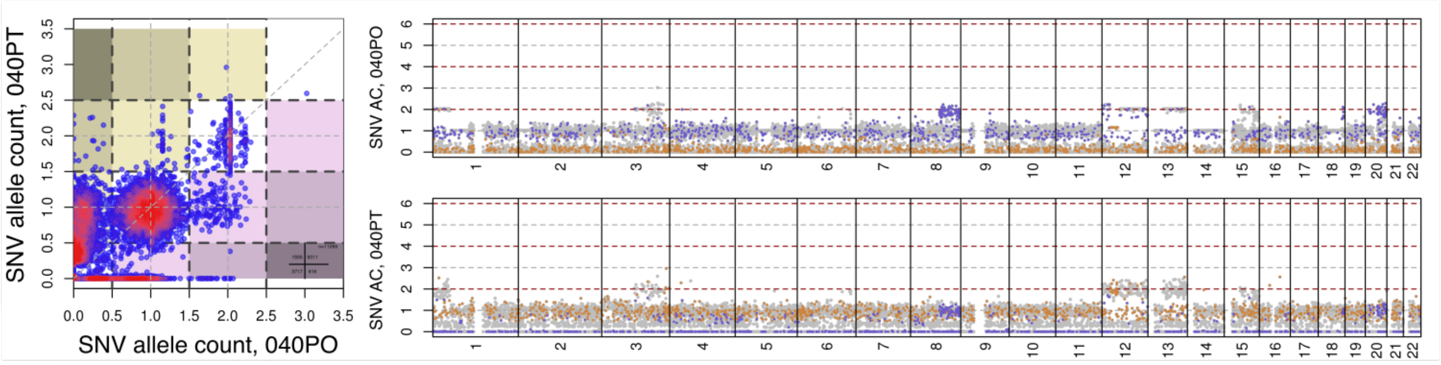


### Patient 048

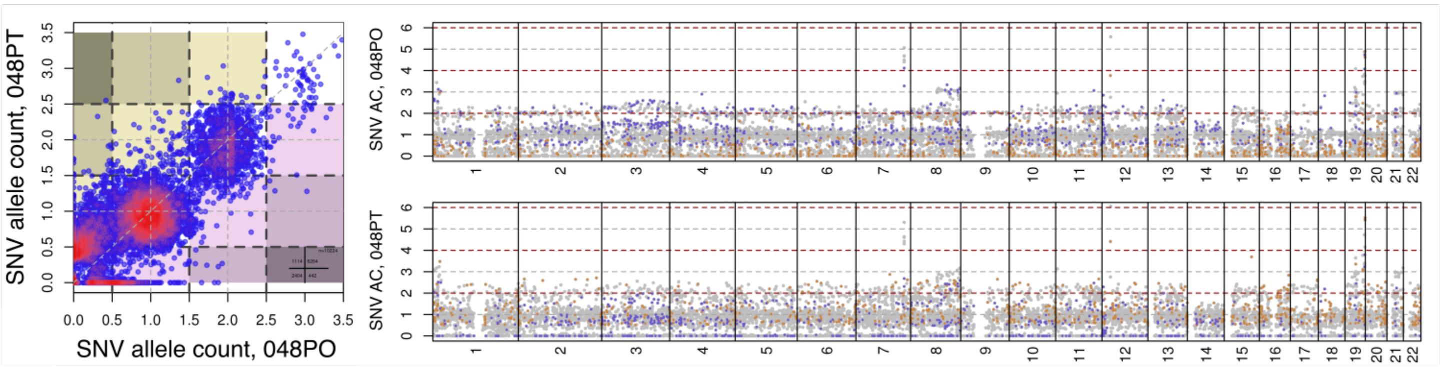


### Patient 050

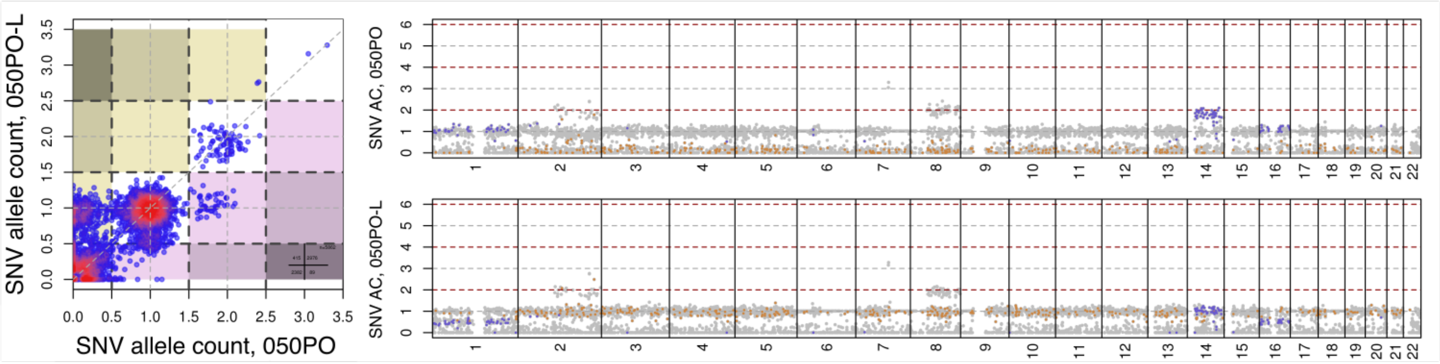

### Patient 059

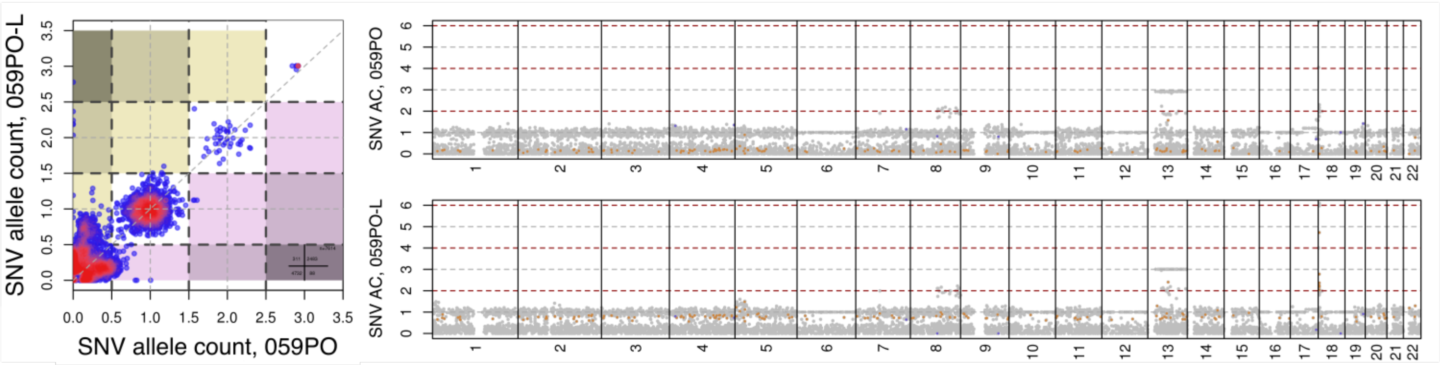


### Patient 064

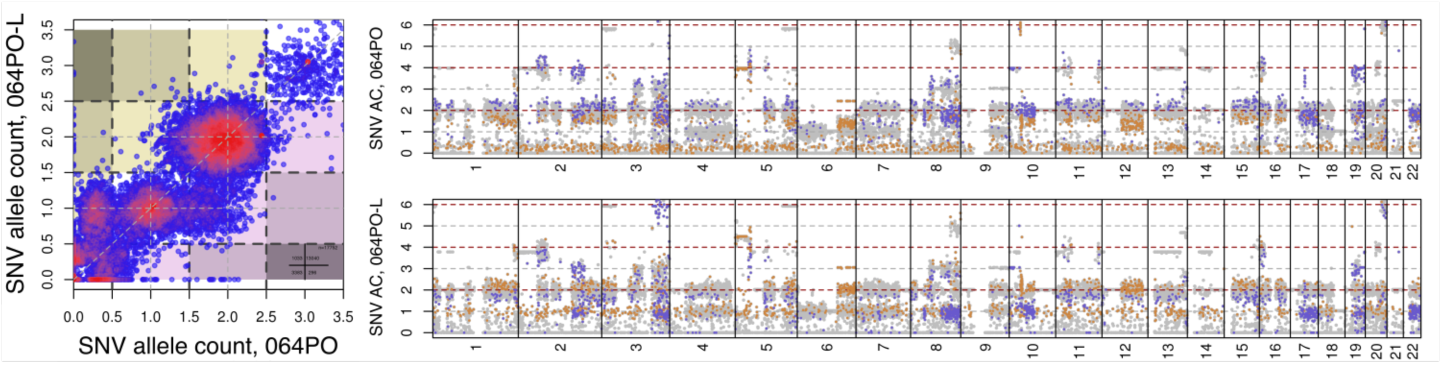


### Patient 065

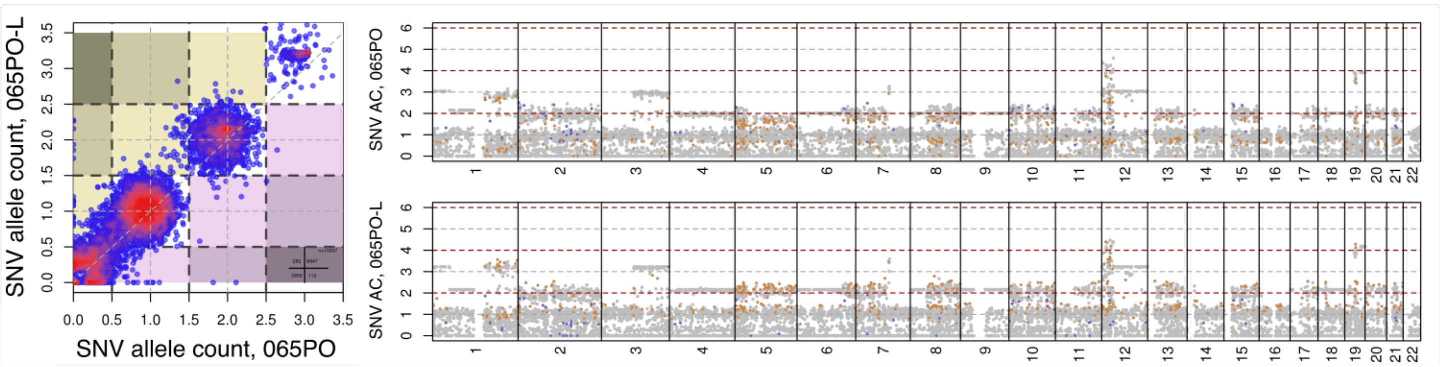


### Patient 069

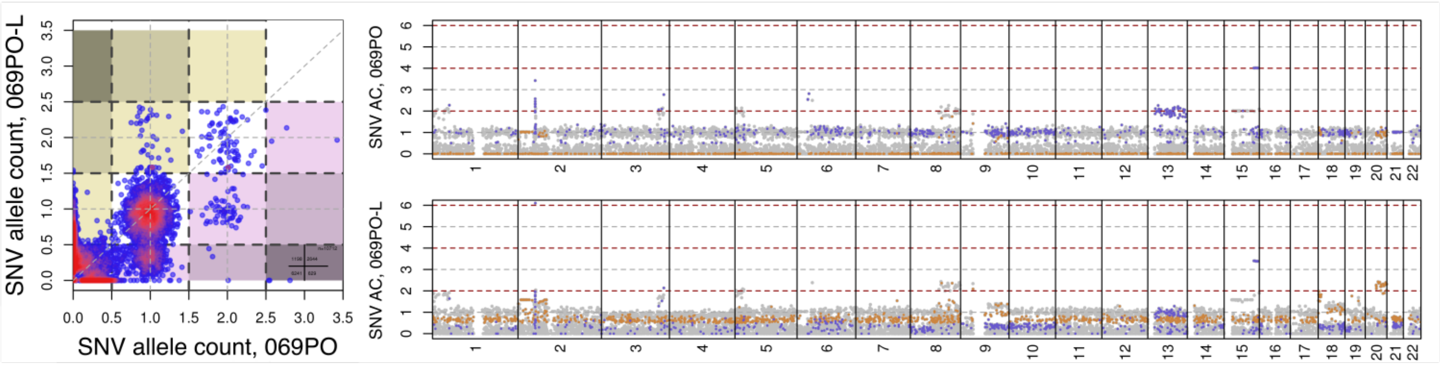

### Patient 072

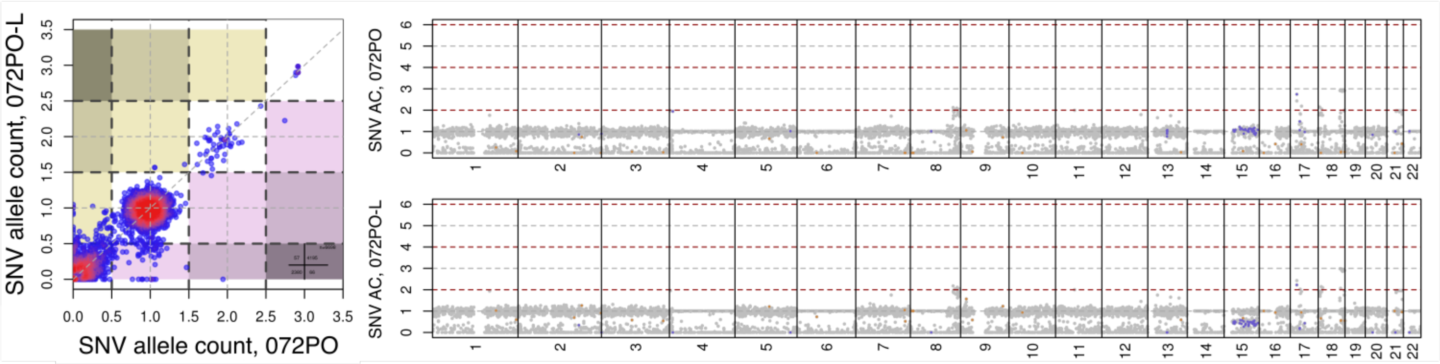


### Patient 077

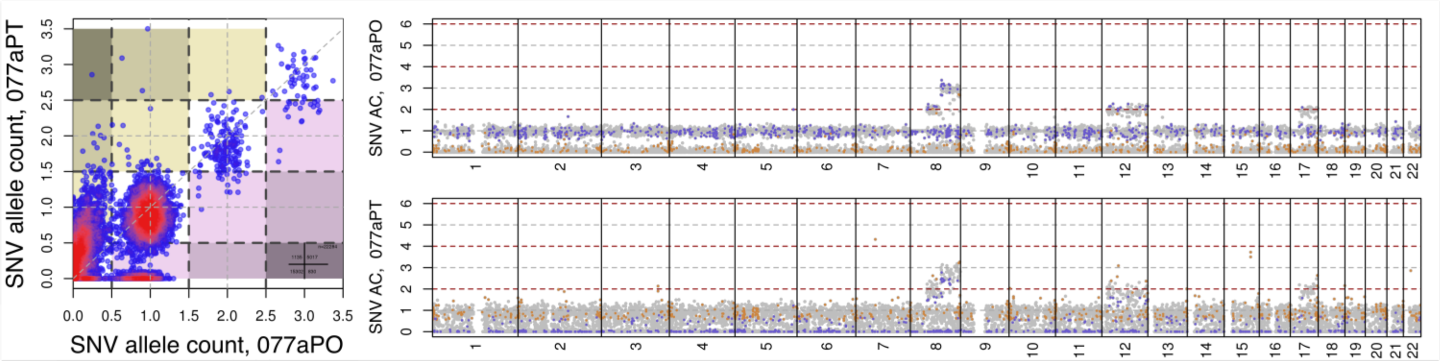


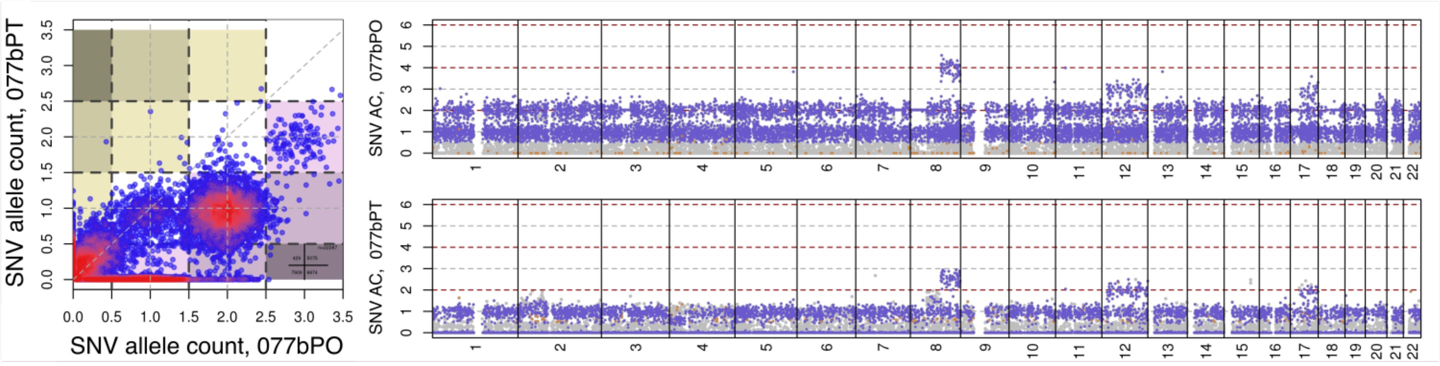


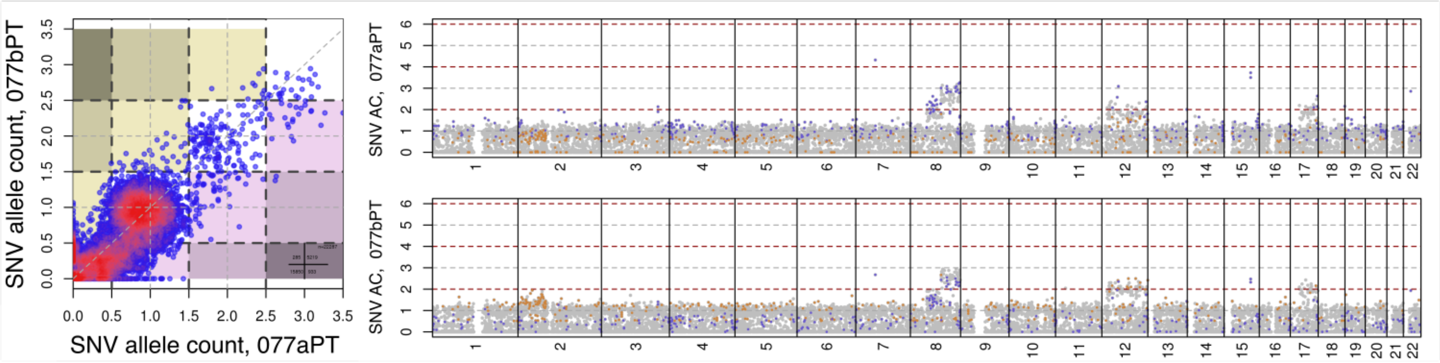

**Patient 077 cont.**

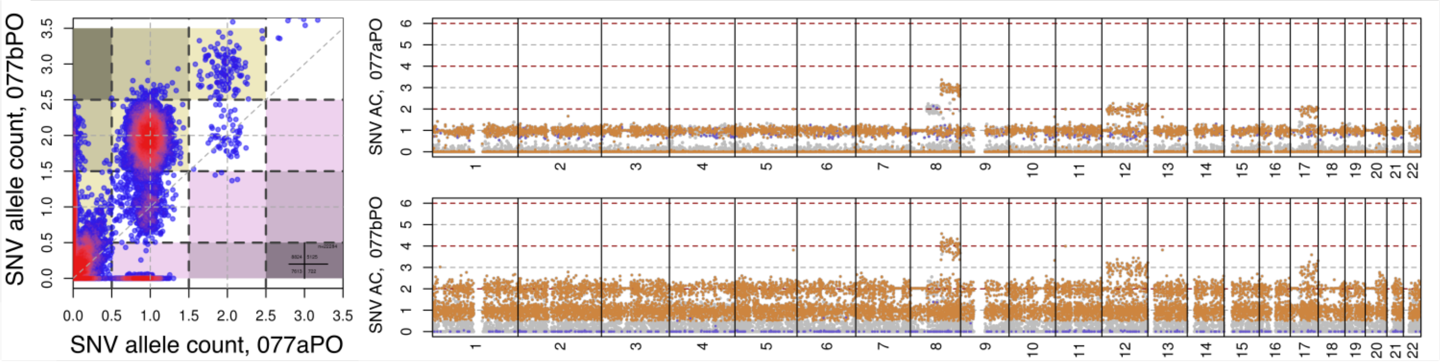


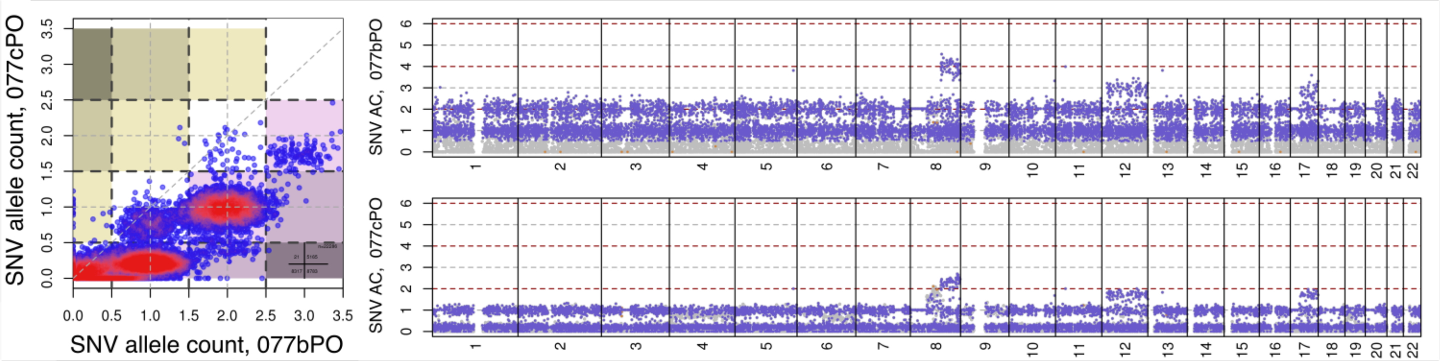


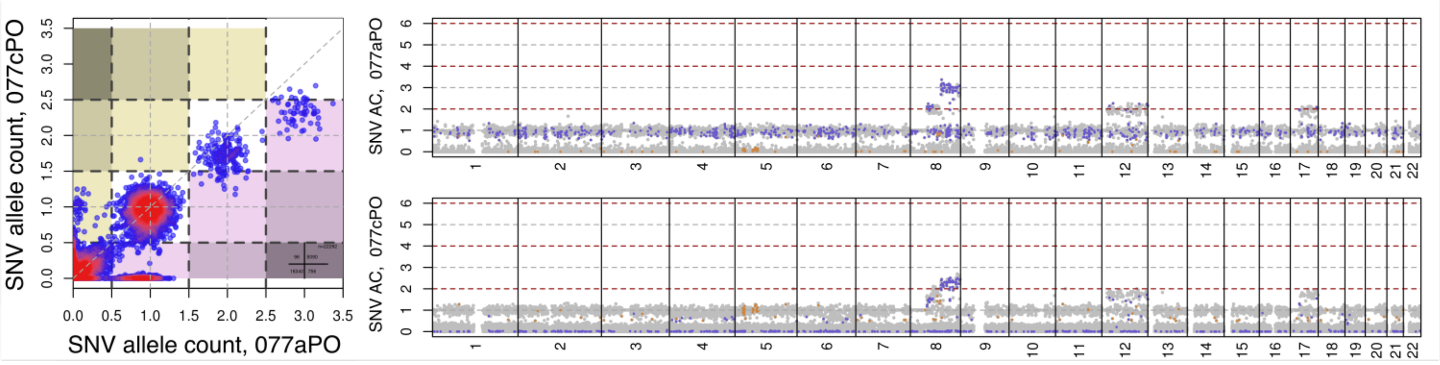


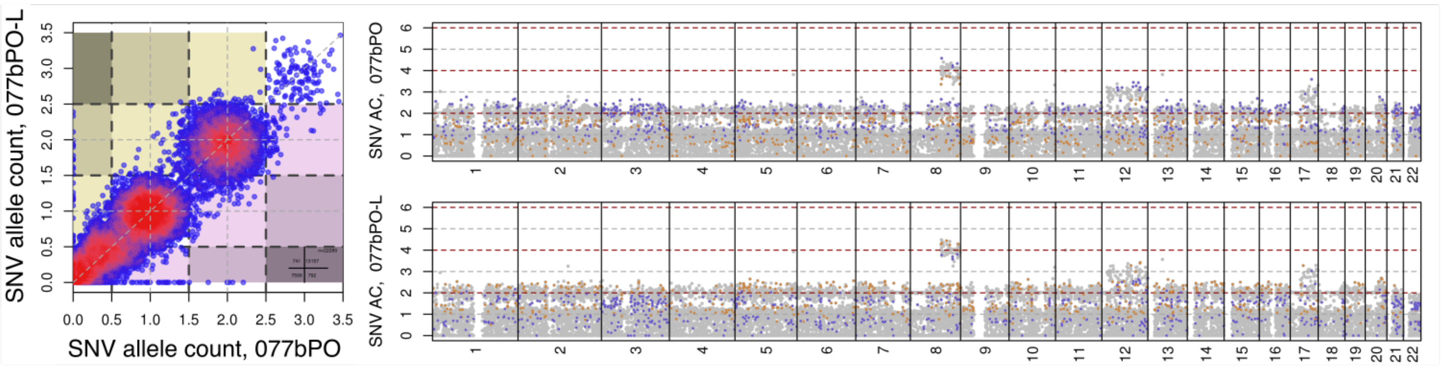

### Patient 080

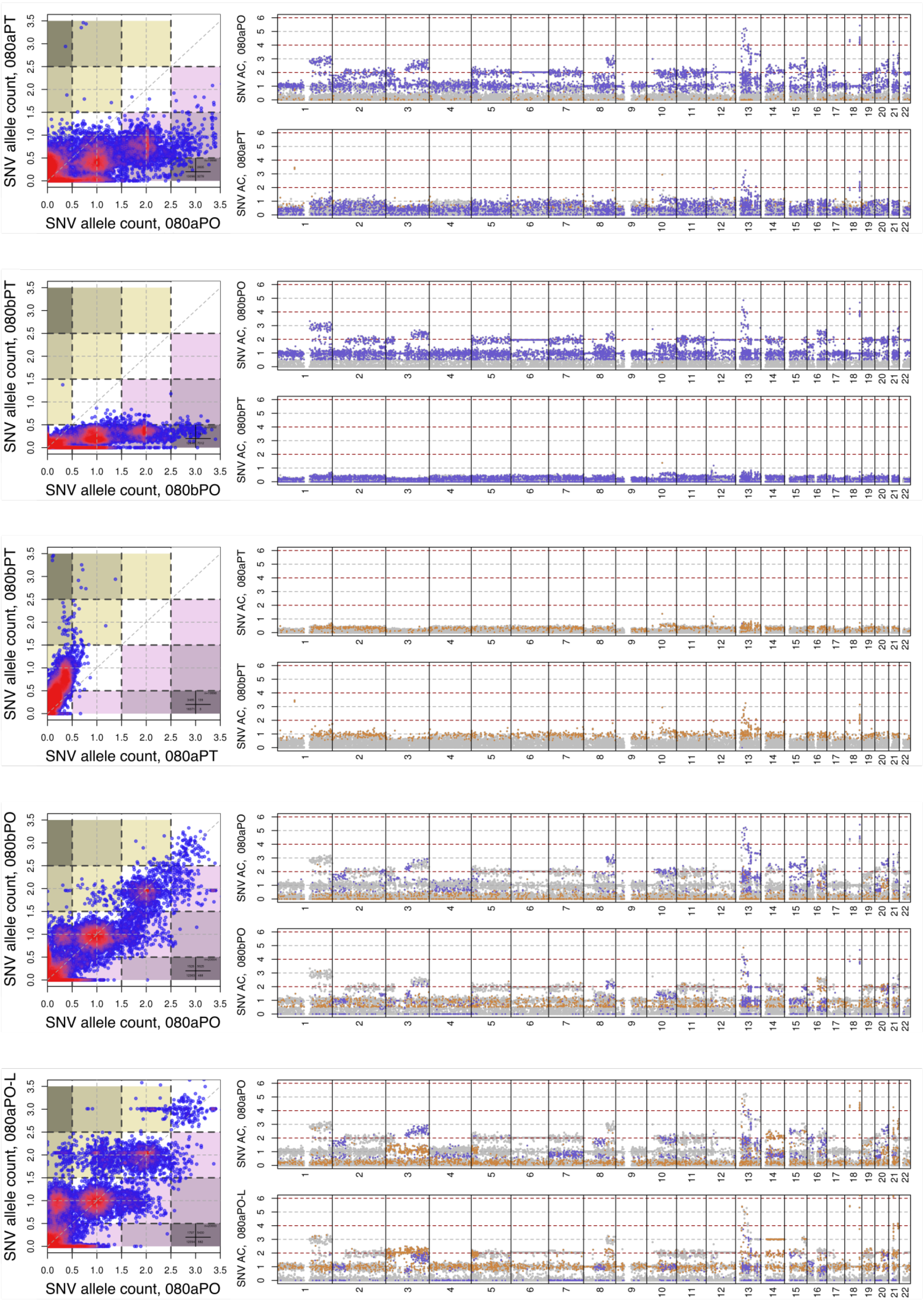

SNV allele count, 080aPT
SNV allele count, 080aPO
SNV AC, 080aPO
SNV AC, 080aPT
SNV allele count, 080bPT
SNV allele count, 080bPO
SNV AC, 080bPO
SNV AC, 080bPT
SNV allele count, 080bPT
SNV allele count, 080aPT
SNV AC, 080aPT
SNV AC, 080bPT
SNV allele count, 080bPO
SNV allele count, 080aPO
SNV AC, 080aPO
SNV AC, 080bPO
SNV allele count, 080aPO-L
SNV allele count, 080aPO
SNV AC, 080aPO
SNV AC, 080aPO-L

### Patient 083

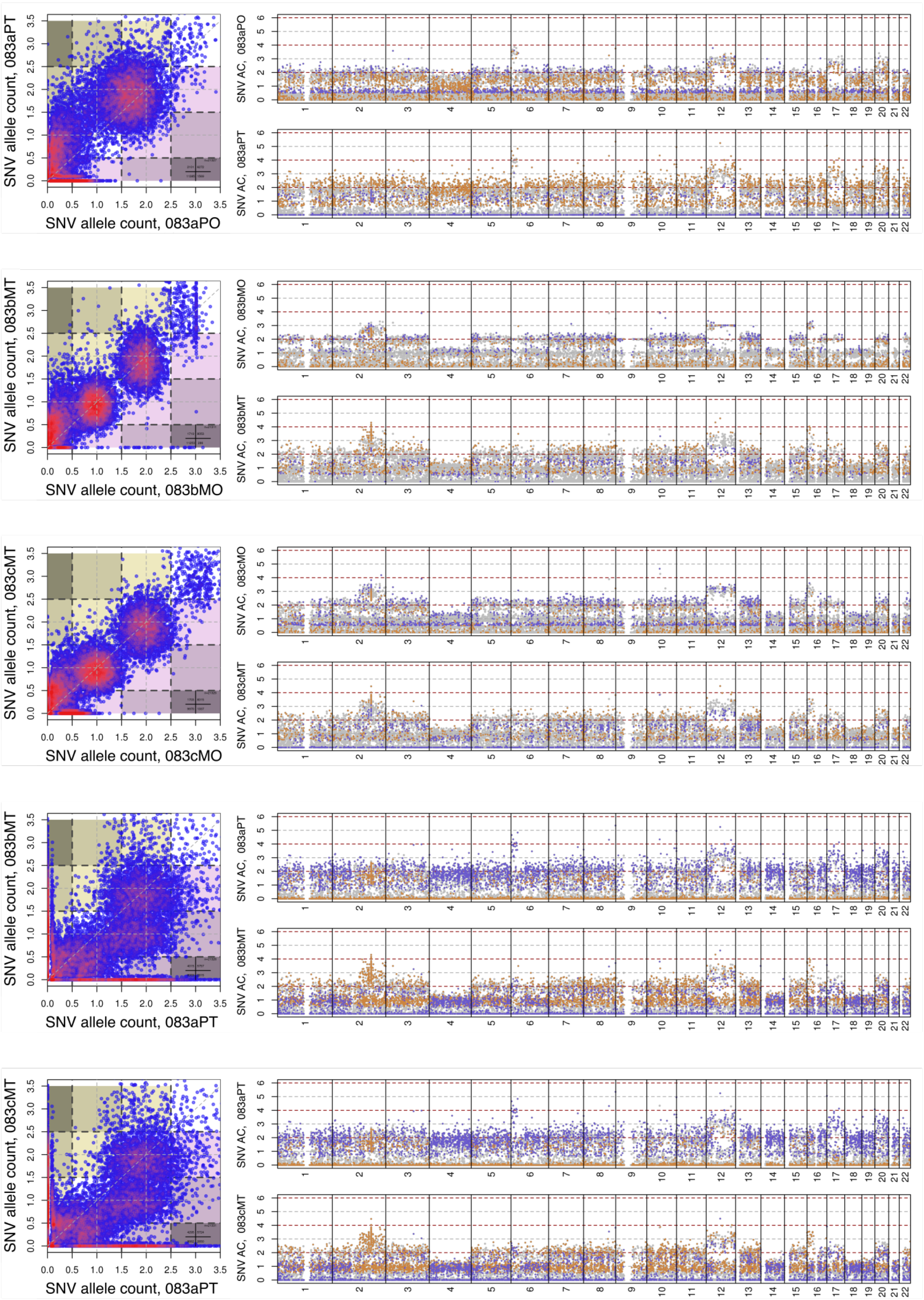
SNV allele count, 083aPT
SNV allele count, 083aPO
SNV AC, 083aPO
SNV AC, 083aPT
SNV allele count, 083bMT
SNV allele count, 083bMO
SNV AC, 083bMO
SNV AC, 083bMT
SNV allele count, 083cMT
SNV allele count, 083cMO
SNV AC, 083cMO
SNV AC, 083cMT
SNV allele count, 083bMT
SNV allele count, 083aPT
SNV AC, 083aPT
SNV AC, 083bMT
SNV allele count, 083cMT
SNV allele count, 083aPT
SNV AC, 083aPT
SNV AC, 083cMT

### Patient 083 cont.

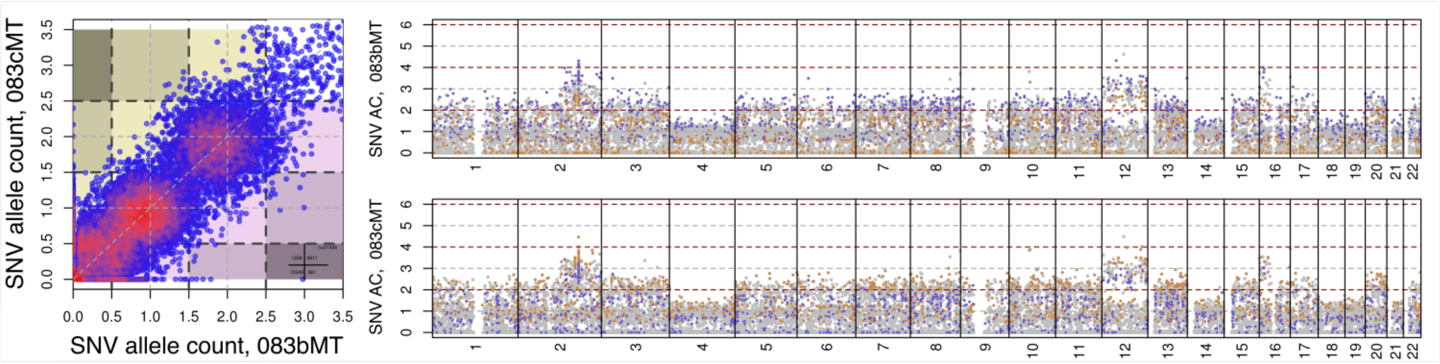


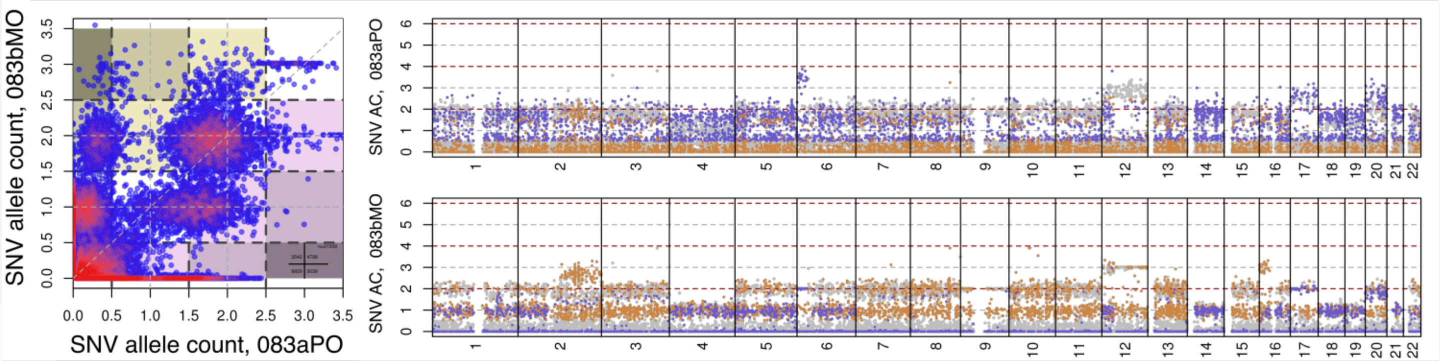


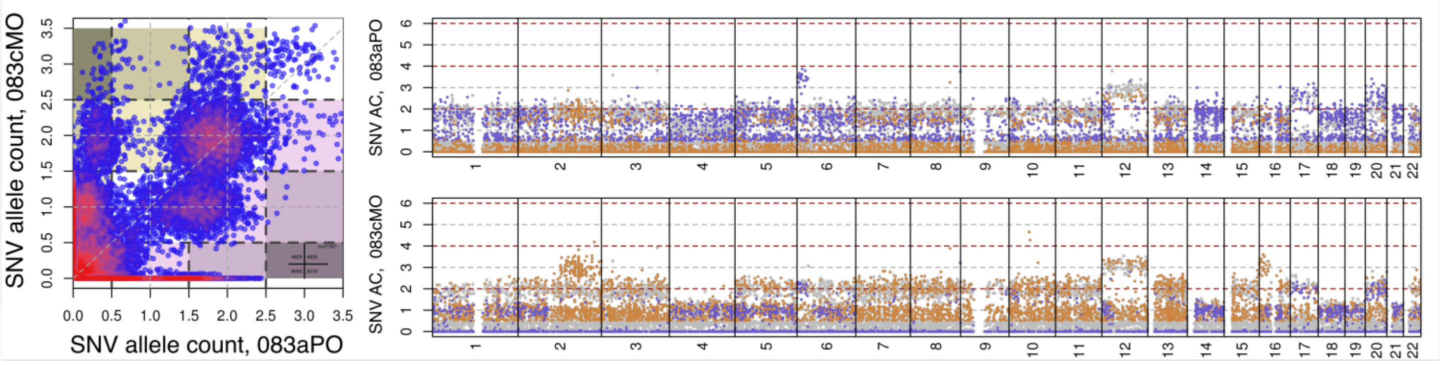


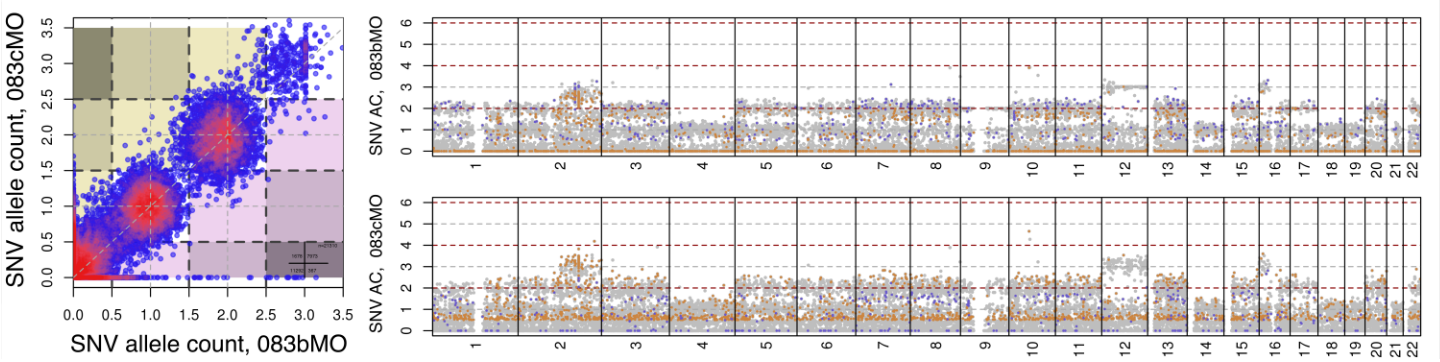

### Patient 086

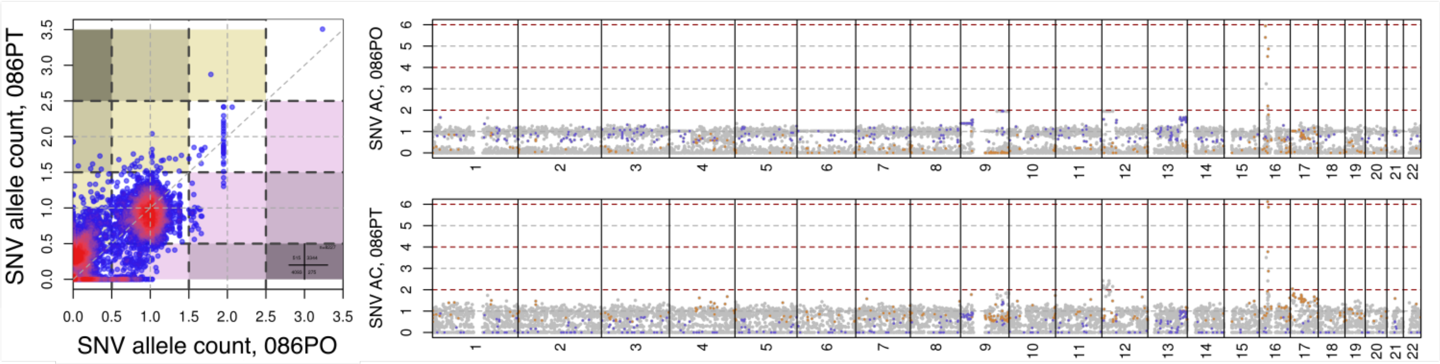


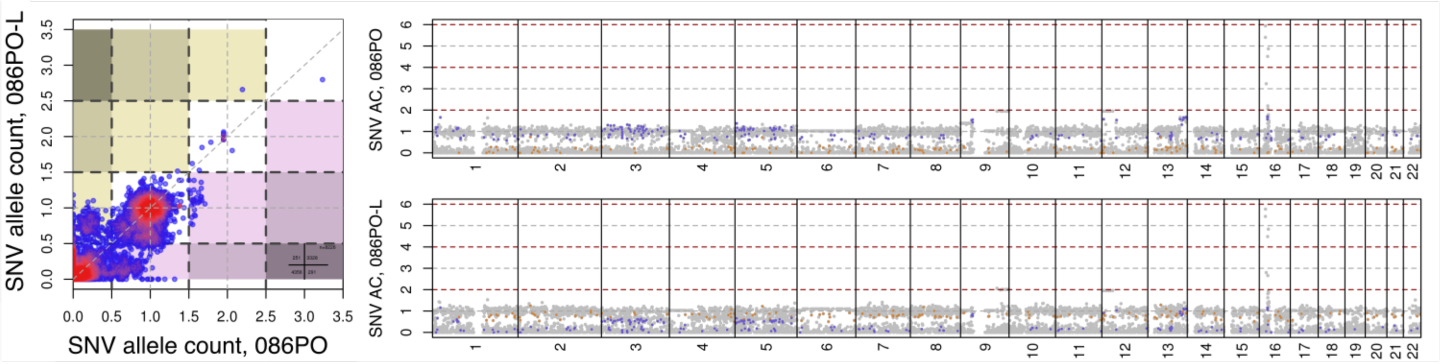


### Patient 100

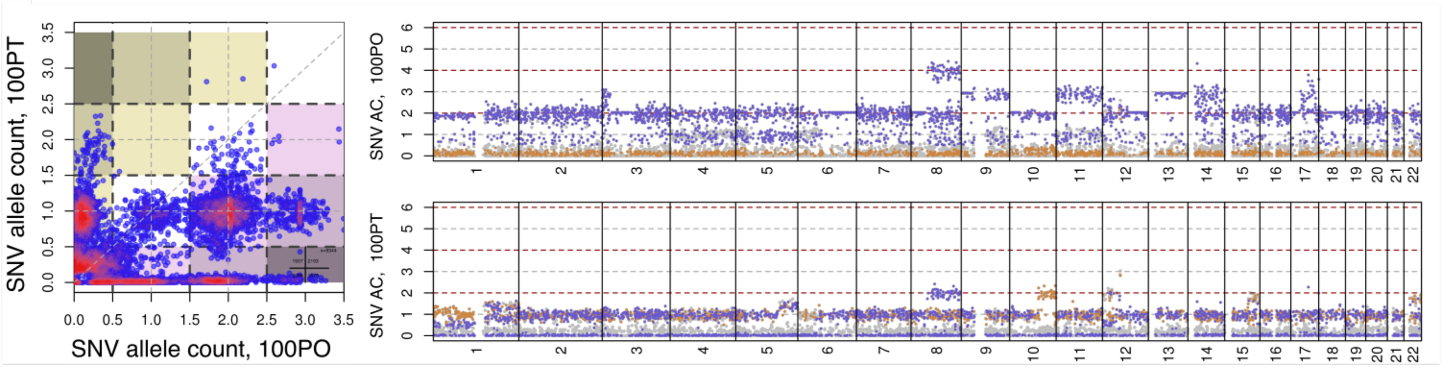
